# Inter-domain distance prediction based on deep learning for domain assembly

**DOI:** 10.1101/2022.12.23.521752

**Authors:** Fengqi Ge, Chunxiang Peng, Xinyue Cui, Yuhao Xia, Guijun Zhang

## Abstract

AlphaFold2 achieved a breakthrough in protein structure prediction through the end-to-end deep learning method, which can predict nearly all single-domain proteins at experimental resolution. However, the prediction accuracy of full-chain proteins is generally lower than that of single-domain proteins because of the incorrect interactions between domains. In this work, we develop an inter-domain distance prediction method, named DeepIDDP. In DeepIDDP, we design a neural network with attention mechanisms, where two new inter-domain features are used to enhance the ability to capture the interactions between domains. Furthermore, we propose a data enhancement strategy termed DPMSA, which is employed to deal with the absence of co-evolutionary information on targets. We integrate DeepIDDP into our previously developed domain assembly method SADA, termed SADA-DeepIDDP. Tested on a given multi-domain benchmark dataset, the accuracy of SADA-DeepIDDP inter-domain distance prediction is 11.3% and 21.6% higher than trRosettaX and trRosetta, respectively. The accuracy of the domain assembly model is 2.5% higher than that of SADA. Meanwhile, we reassemble 68 human multi-domain protein models with TM-score ≤0.80 from the AlphaFold protein structure database, where the average TM-score is improved by 11.8% after the reassembly by our method. The online server is at http://zhanglab-bioinf.com/DeepIDDP/.

## 1 Introduction

Protein structure prediction methods based on deep learning have made significant progress over the past decades [1-4]. These methods have been proven to be effective in improving protein structure prediction accuracy, especially in the absence of homologous templates [5]. In general, protein structure prediction methods based on deep learning can be divided into three categories. First is inter-residue contact prediction methods, where 8Å is used as a criterion to classify residue contact and non-contact relationships. The representative methods include PSICOV [6], CCMpred [7], MSA Transformer [8], and others. PSCOV proposes the use of sparse inverse covariance estimation techniques to deal with the coupling effects. CCMpred is a PLM-based contact prediction method that integrates GPU and CPU to accelerate prediction efficiency. MSA Transform is an unsupervised contact prediction method that proposes a variant of axial attention, which shares a single attention map across rows. The second category is composed of inter-residue distance prediction methods, which not only contain more abundant information than contact prediction but also have more physical constraints on protein structure [9]. RaptorX is the first method to apply residual network (ResNet) to predict residue distance [9], which integrates sequence and paired features. trRosetta is the representative distance prediction method, which describes additional geometrical constraint information by predicting three orientations between residues [10]. Based on trRosetta, trRosettaX further improves the accuracy of distance prediction, which proposes a new multi-scale network Res2Net and an attention module while automatically integrating homologous template information [11]. Aside from these, many other distance prediction methods, such as MULTICOM [12], tFold [13], TripletRes [14] and DMPfold [15], are available. The third category consists of the end-to-end structure prediction methods, which uses attention and rotation-equivariant networks to directly generate the 3D structure of proteins [11]. These methods are known as RoseTTAFold and AlphaFold2. RoseTTAFold explores network architecture incorporating related ideas and obtains the best performance with a three-track network in which information at the 1D sequence level, 2D distance map level, and 3D coordinate level are successively transformed and integrated [16]. In AlphaFold2, the attention-based module (Evoformer) is used to extract co-evolutionary information, and a structural module based on invariant point attention is used to directly generate atomic coordinates [17]. In CASP14, the average GTD-TS score of AlphaFold2 is up to 92.4, and even on hard proteins, the average GTD-TS score is 87.0 [18].

The greatly improved prediction of protein 3D structures achieved by AlphaFold2 [19], but challenges remain on multi-domain protein structure modelling. On the one hand, many multi-domain proteins absence co-evolutionary information and full-length templates [20], which may be one of the main reasons for the reduced prediction accuracy. On the other hand, AlphaFold2 has high computer hardware requirements [21]. Therefore, domain assembly techniques are considered effective methods to model the full-chain structure of multi-domain proteins, which are almost unaffected by the length of the full-chain protein [22]. AIDA, DEMO and SADA are the representative assembly methods. AIDA developed an *ab*-*initio* folding fast docking algorithm for full-length modelling by changing the linker configuration to simulate the domain assembly of multi-domain protein structures [23]. DEMO proposed a rigid body assembly method based on constructing multi-domain structures from a single domain, which determines the domain orientation through the template distance profile detected by local and global templates [24]. In our previously proposed SADA approach, a multi-domain protein structure database (MPDB) was constructed for the full-chain analogue detection using individual domain models. Based on the detected analogue, an energy function assisted by deep learning network GeomNet is designed to guide domain assembly [25]. The results show that SADA can be an effective complement to AlphaFold2 in multi-domain protein modelling [21]. However, we consider that the performance of SADA could be further improved if a network could be trained to specifically predict inter-domain distances for multi-domain proteins.

In this work, we first propose a data enhancement strategy termed domain-pair multiple sequence alignment (DPMSA). Then, two new features including inter-domain residue coupling score (IMCP) and inter-domain average contact potential (ICOP), are designed to focus on inter-domain residue pairs. Finally, a deep network, which combines attention mechanism and deep residual block, is constructed to predict the inter-domain residue distance distribution of multi-domain proteins. We integrate DeepIDDP into the SADA domain assembly method, termed SADA-DeepIDDP. The experimental results show that SADA-DeepIDDP outperforms the comparative method in inter-domain distance prediction and multi-domain protein modelling.

## 2 Materials and Methods

### 2.1 Benchmark dataset

To train a deep neural network to predict inter-domain residue pair distance distributions, we constructed a non-redundant dataset from the MPDB (up to July 2022, there were 48,224 multi-domain proteins). First, the CD-HIT [26] sequence clustering tool was used to obtain 10,593 multi-domain proteins with 40% sequence identity cut-off. Then we further filtered the proteins based on the following criteria:(i) minimum resolution of 2.5Å, (ii) limit of 50–1,024 residues for each multi-domain protein and (iii) limit of 6 for each multi-domain protein. Finally, the dataset contained 9,248 multi-domain proteins (8,820 for training and 464 for validation).

### 2.2 Domain-Pair MSA

As shown in Figure 1(A), MSA consists of basic MSA and DPMSA. The basic MSA uses the HHblits [27] tool to search in the Uniclust30_2018_08 and BFD datasets [28]. The detailed parameters of the HHblits are shown in Supplementary Table S1. During the iterative search, we gradually relax the boundaries of the e value, and the search stops when at least 2,000 sequences cover 75% or 5,000 sequences cover 50% meet either of the two criteria.

**Figure 1.**
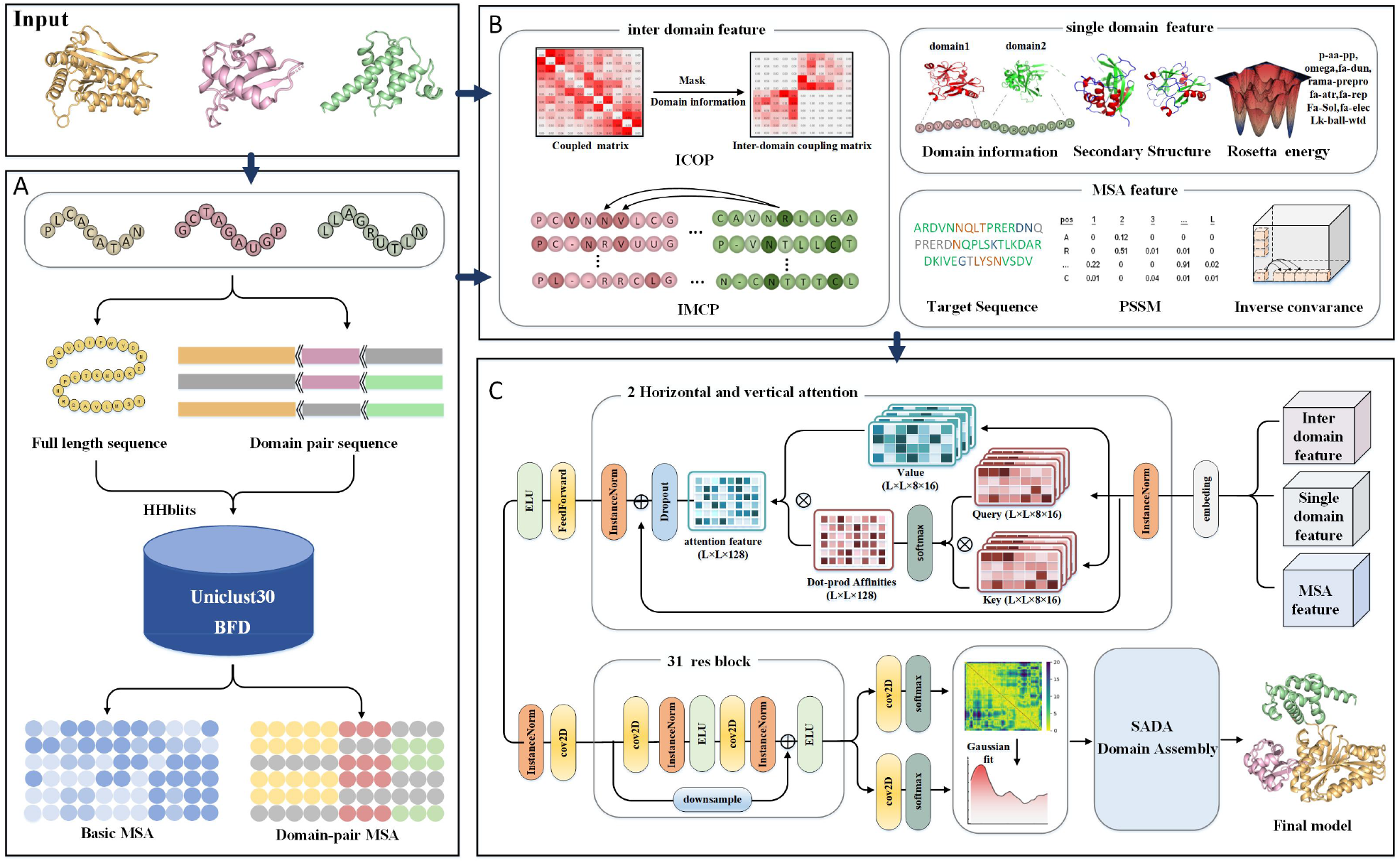
**(A)** Pipeline of DeepIDDP. **(A)** Domain-pair MSA data augmentation method where MSA is searched from full length and domain-pair sequences. **(B)** Feature extraction through which inter-domain, single-domain and MSA features are extracted from the single-domain structure and enhanced MSA data. **(C)** Network architecture. Contains an attention module and a deep residue blocks module.

We use the following two strategies to generate DPMSA for data enhancement:

1. Performing mask operation (replacing the amino acid symbol with “-”) on a domain randomly on the full-length sequence, generating a domain-pair sequence, and then performing MSA search.
2. Performing MSA search on single-domain sequences, and then clustering and concatenating MSA from various domains to generate a domain-pair MSA.

Detailed data enhancement methods can be found in Supplementary Figure S1. The search criteria are the same as above except that the sequence coverage is 90%. In addition, the number of basic MSA and DPMSA is kept at 1:1 to ensure that the network does not favour either side.

### 2.3 Feature extraction

As shown in Figure 1(B), inter-domain features (ICOP and IMCP), single-domain features (domain information, secondary structure and Rosetta centroid energy terms), and MSA features (amino acid sequence, location-specific frequency matrix and inverse covariance matrix) are extracted and fed into the deep neural network. All the details of the features are listed in Supplementary Table S2.

#### 2.3.1 Inter-domain feature

Residues within each domain usually have strong interactions with each other, stabilising the conserved domain structure, while only a fraction of all residues are involved in the interactions with other domains [29]. Thus, we design two inter-domain features (ICOP and IMCP) to characterise these inter-domain residues that may interact with each other, which can allow the network to focus more on these residue pairs.

ICOP is an *L*×*L* dimensional feature, and for every element in the ICOP, coupling score from the output of the pre-trained MSA Transformer [8] model is used in calculation based on the following formula:

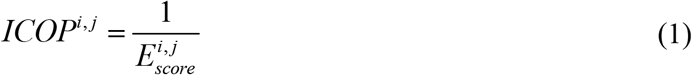

where 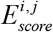 is the coupling fraction of residue *i* and residue *j* of MSA Transform output. We calculate only the residue pairs between different domains.

IMCP is an *L*×*L* dimensional feature, which is used to characterise the tendency of some residues to form mutual residue pairs. The feature is calculated as follows:

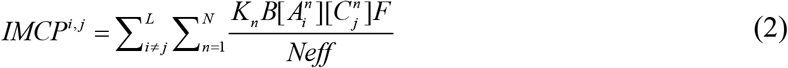

where *K*_*n*_ is the weight of the n-th MSA, 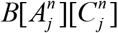 is the residue contact potential energy between residues at positions *i* and position *j* of the n-th MSA in the interaction matrix *B* [30], *Neff* is the number of valid sequences for MSA, and *F* is a flag indicating whether the *i* and *j* residues belong to the same domain.

#### 2.3.2 Single-domain features

Single-domain features are important for determining the spatial position of residues within domains, which can indirectly affect the prediction of inter-domain distances. In this work, we take domain information, secondary structure and Rosetta centroid energy terms as single-domain features. The domain information is obtained according to the sequence alignment of the target sequence and individual domain structure. It is used to annotate the residues of sequence, where residues belonging to different domains are labelled differently in order. The secondary structure information is obtained by DSSP [30], which divides the structure into *α*-fold, *β*-helix and loop. The Rosetta centroid energy terms [31, 32] are introduced as the third feature of single domain, which includes the one-body-terms (omega, p-aa-pp, fa-dun and rama-prepro) and the two-body-terms (fa-atr, fa-rep, fa-sol, lk-bal-wtd and fa-elec).

#### 2.3.3 MSA feature

We extract four conventional features from MSA, including one-hot encoding of target sequence, position frequency specificity matrix (PSSM), position entropy and inverse covariance matrix. PSSM represents the evolutionary trend of each amino acid column [33]. The position entropy is a measure of sequence randomness. The inverse covariance matrix represents the marginal relationship between various positions in a protein sequence, calculated by the frequency of residues and frequency of residue pairs at the given location in MSA [34].

### 2.4 Network architecture

Figure 1C shows the overall network architecture that includes two modules: a horizontal and vertical attention module and a residual module. The features mentioned in section 2.3 are fed into the network to predict the inter-domain residue pair distance (ranging from 2Å to 22Å in 0.5 steps into 41 equally spaced bins, plus one bin indicating that the residues are uncorrelated). Firstly, the axial attention module integrates all input features through the embedding layer and converts the number of input features to 128 through the normalisation and the linear layers. Secondly, the axial attention module applies two layers of self-attention along the rows and columns, each layer containing eight heads. Queries, Keys, and Values are obtained from the input through the linear layer. The attention map is the product of values and results of the softmax after the product of queries and keys. Finally, the output is fed to the residual module through a feedforward network including a normalisation and a dropout layer. In the residual module, the number of input features is converted to 64 (2D convolution with filter size 1) at the first layer, and then a stack of 31 basic residual blocks with dilation is applied. Expansion cycles through 1, 2, 4, 8 and 16 (a total of six full cycles). After the last residual block, the network branches into two independent paths (one per objective) each consisting of a 2D convolution followed by a softmax activation. All convolution operations, except the first and last, use 64 3 × 3 filters. ELU activations are applied to the entire network.

### 2.5 Model training

During training, we use categorical cross-entropy to measure the network loss, The calculation formula is as follows:

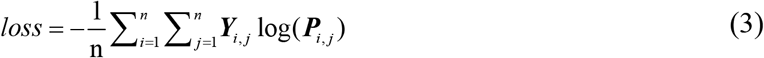

where *Y*_*i, j*_ is the true distance vector of residue pairs *i* and *j*, and *P*_*i, j*_ is the distance distribution vector predicted by the network of residue pairs *i* and *j*.

Meanwhile, the protein is randomly sliced over 320 amino acid domains to accommodate the 320 residues limit. All training proteins are randomly executed in each training epoch, and a total of 100 epochs are performed. Adam optimizer with a learning rate of 1e-3 is used. All trained parameters are subject to the *L*_2_ penalty of 1e-4 weights. The 15% dropout rate is used to maintain the probability. It takes about 7 days to train the network on an NVIDIA Tesla V100s GPU.

## 3 Results

In order to test accuracy of the predicted inter-domain distance, DeepIDDP is compared with three other methods, including trRosettaX [11], trRosetta [10] and SADA [21]. Meanwhile, to test the effectiveness of inter-domain distance on domain assembly method, we integrate the distance of DeepIDDP into SADA and compare it with the original SADA method. Finally, by comparing with AlphaFold2 on 17 CASP14 multi-domain targets and 68 multi-domain proteins with TM-score ≤0.80, we further demonstrate that inter-domain distance can improve the domain orientations of the multi-domain protein models.

### 3.1 Evaluation of inter-domain distance prediction

In this section, we compare the performance of DeepIDDP with that of representative distance prediction methods trRosettaX, trRosetta and SADA in a benchmark multi-domain protein set. The multi-domain dataset is obtained from published SADA papers and consists of 166 2-domains(2dom), 69 3-domains(3dom), 40 ≥4-domain(m4dom) and 81 discontinuous domains(2dis). The results of the comparative methods are obtained from the executable program of the published paper. Meanwhile, all inter-domain residual pairs with a true distance of less than 15Å [9] is considered because the number of contacts (true distance is less than 8 Å) of inter-domain residual pairs is small. The comparison results are summarised in Table 1 and the detailed results are shown in Supplementary Table S3.

**Table 1.** Summary of the results of inter-domain distance prediction for 356 multi-domain proteins. SADA-DeepIDDP represents the distance predicted by DeepIDDP and SADA represents the distance predicted by GeomNet. MAE represents the mean absolute error of all residue pairs between domains, where the distance of the residue pairs is less than 15 Å. Top-*L* represents the prediction accuracy of the top *L* for the inter-domain residues.

| Method | MAE | | | | | Top-$L$ | Precision | Recall | F1 |
| --- | --- | --- | --- | --- | --- | --- | --- | --- | --- |
|  | 2dom | 3dom | m4dom | 2dis | All |  |  |  |  |
| **SADA-DeepIDDP** | **2.65** | **2.78** | **3.21** | **2.78** | **2.77** | **0.692** | **0.545** | 0.674 | **0.577** |
| SADA | 3.45 | 3.66 | 3.63 | 3.51 | 3.52 | 0.572 | 0.418 | 0.637 | 0.471 |
| trRosetta | 3.41 | 3.55 | 3.49 | 3.45 | 3.47 | 0.569 | 0.439 | 0.577 | 0.472 |
| trRosettaX | 3.36 | 3.56 | 3.48 | 3.32 | 3.41 | 0.622 | 0.432 | **0.714** | 0.502 |

Overall, SADA-DeepIDDP outperforms the other three methods on the multi-domain protein test set. The mean absolute error (MAE) value of SADA-DeepIDDP is 2.77 Å, which is 21.3%, 18.7% and 20.1% lower than that of SADA, trRosettaX and trRosetta, respectively. Among them, SADA-DeepIDDP performed better on 2dom and 3dom than SADA, trRosettaX, and trRosetta, respectively. The most likely reason is that the training set consisted mainly of 2dom and 3dom (93.7% of the total training proteins), resulting in the neural network biased towards predictors of both types of multi-domain proteins. The top-*L* accuracy of SADA-DeepIDDP is 0.692, which is 21.0% higher than that of SADA, 11.3% higher than that of trRosettaX and 21.6% higher than that of trRosetta. Figure 2 shows the head-to-head comparison of SADA-DeepIDDP versus SADA, trRosettaX and trRosetta on top-*L*, respectively. In addition, we use three machine metrics, precision, recall and F1, to comprehensively measure the predictive performance of the three methods. In both precision and F1, SADA-DeepIDDP is superior to the other two methods, which are 26.0% and 14.9% higher than the suboptimal method. However, the recall of trRosettaX is higher than that of SADA-DeepIDDP. This is most likely because trRosettaX considers a number of inter-domain residue pairs, resulting in high recall and low accuracy. In summary, SADA-DeepIDDP outperforms the other three methods in inter-domain distance prediction.

**Figure 2.**
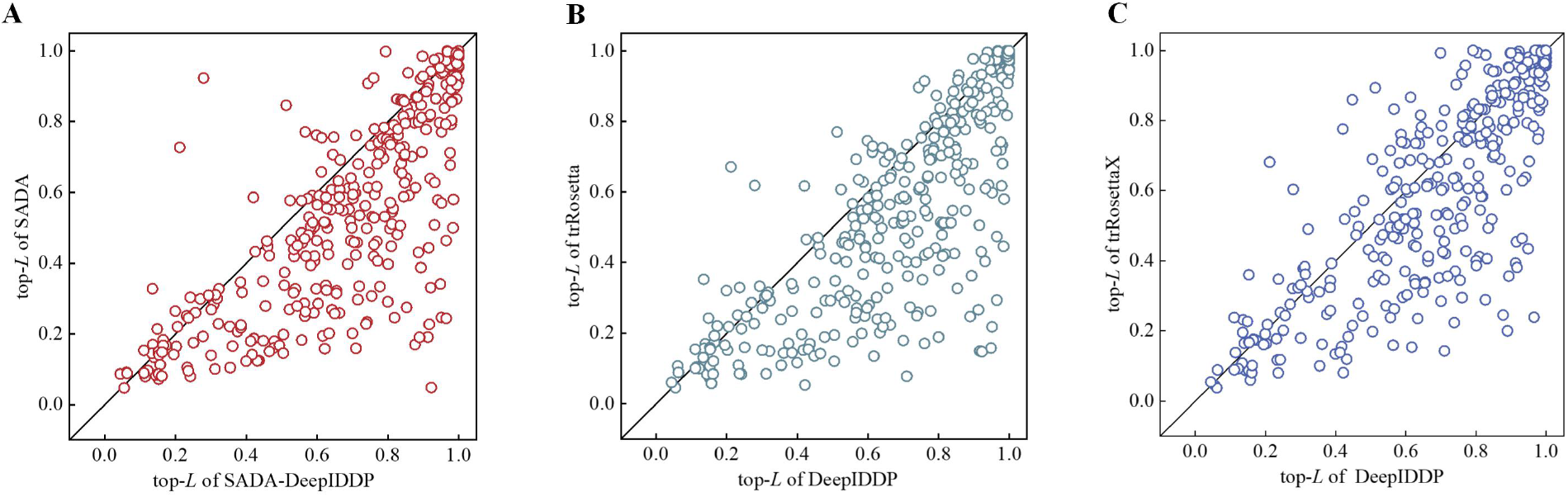
Comparison of SADA-DeepIDDP, SADA, trRosettaX, and trRosetta. **(A)** Head-to-head top-*L* accuracy comparison of SADA-DeepIDDP and that created by SADA. **(B)** Head-to-head top-*L* accuracy comparison of SADA-DeepIDDP and that created by trRosetta. **(C)** Head-to-head top-*L* accuracy comparison of SADA-DeepIDDP and that created by trRosettaX.

### 3.2 Effectiveness of inter-domain distance on the assembly method

To further test the performance of DeepIDDP, we integrate it into our previously developed domain assembly method SADA and reassemble the proteins on the multi-domain dataset. Table 2 summarises the results of SADA-DeepIDDP and the original SADA method. The detailed results are listed in the Supplementary Table S4.

**Table 2.** Summary of modelling results for the benchmark multidomain set. TM-score represents the average TM-score of final full-chain models. #TM-score ≥ 0.5 represents the number of proteins with TM-score ≥ 0.5[35]. iRMSD [36] represents interface RMSD. The values in the last column are the results of the student’s *t*-test based on the comparison with the TM-score of SADA-DeepIDDP.

| Method | TM-score | | | | | #TM-score<br>$\geq 0.5$ | iRMSD | $p$-value |
| --- | --- | --- | --- | --- | --- | --- | --- | --- |
|  | 2dom | 3dom | m4dom | 2dis | All |  |  |  |
| **SADA-DeepIDDP** | **0.85** | **0.77** | **0.66** | **0.90** | **0.82** | **339** | **2.71** | **-** |
| SADA | 0.83 | 0.72 | 0.60 | 0.89 | 0.80 | 328 | 3.17 | 4.59E-4 |

**Table 3.** Summary of modelling results for 68 human multi-domain proteins.

| Method | TM-score | #TM-score$\geq 0.5$ | iRMSD | *p*-value |
| --- | --- | --- | --- | --- |
| **SADA-DeepIDDP** | **0.68** | **53** | **4.56** | **-** |
| AlphaFold2 | 0.60 | 49 | 5.72 | 7.98E-7 |
| SADA | 0.66 | 51 | 5.13 | 9.86E-3 |

Overall, the average TM-score of the SADA-DeepIDDP model on all proteins is 0.82, which is 2.5% higher than that (0.80) of SADA. Among them, the TM-score of SADA-DeepIDDP for 2dom, 3dom, m4dom and 2dis are respectively 2.4%, 6.9%, 10.0%, and 1.1% higher than that of SADA. Obviously, for 3dom and m4dom, SADA has lower accuracy, because it is difficult to detect effective structural analogues when the number of structural domains increases. However, the DeepIDDP can provide more inter-domain information than distance prediction methods (such as GeomNet) to capture the orientation between domains, and further improve the accuracy of the final model [25].

Figure 3 intuitively reflects the comparison of TM-scores of the two methods on different types of multi-domain proteins. SADA-DeepIDDP correctly assembles 339 of the 356 proteins (TM-score ≥ 0.5), representing 95.2% of the total and 3.39% higher than SADA. In addition, we use iRMSD to evaluate domain-domain assemble poses, which is almost unaffected by the size of the domains (the details of iRMSD can be found in Supplementary Equation S1) [37]. The iRMSD of SADA-DeepIDDP is 14.5% lower than that of SADA, which further indicates that DeepIDDP can effectively capture the inter-domain orientation. The difference of *p*-value (4.59E-4) reflects the statistical significance of the two methods. Figure 4 is an example (PDB ID:4c0aB). Although SADA correctly assembles domains 1 (orange) and 2 (green), it has a large deviation in the domain 3 (cyan) position. SADA-DeepIDDP correctly captures the orientations between domains 1, 2 and 3. The right side of Figure 4 uses the PAE (predicted alignment error) plot to reflect the inter-domain accuracy[38].

**Figure 3.**
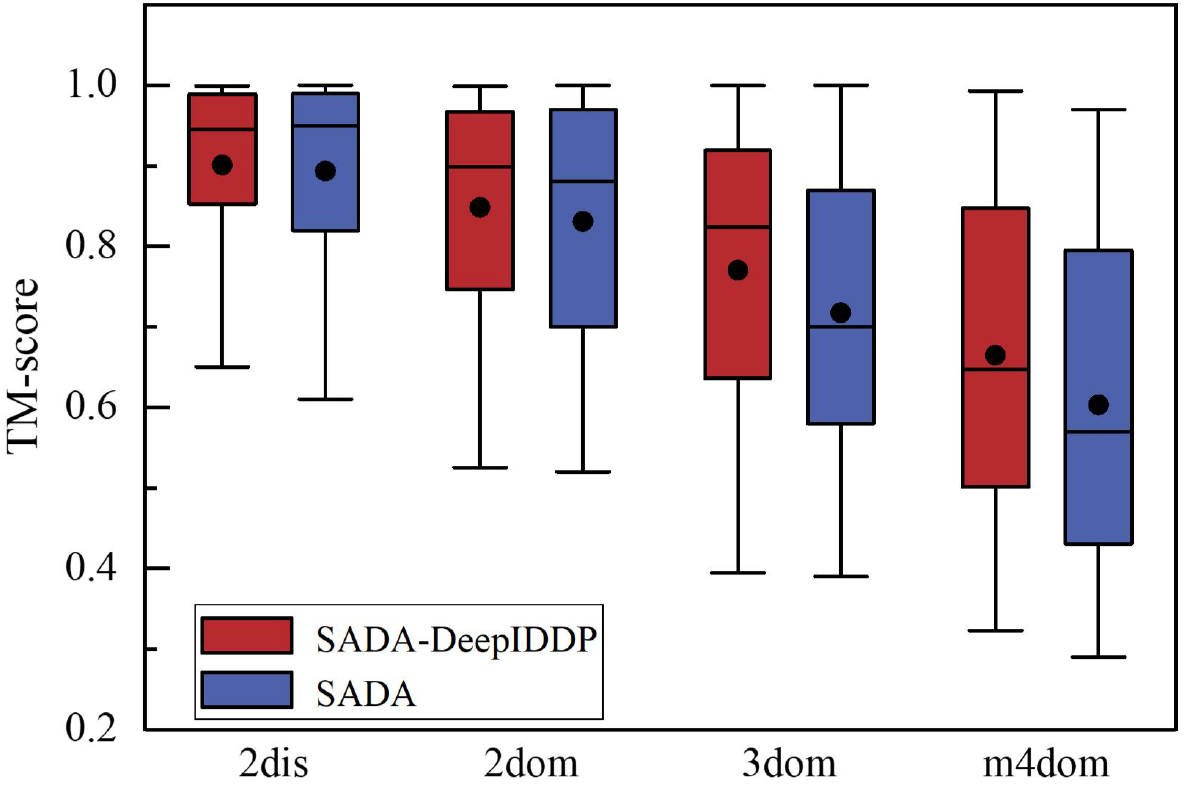
Boxplot of TM scores for SADA-DeepIDDP and SADA assembly models. where the circle and horizontal lines in the box represent the mean and median top-*L*, and the horizontal lines on the top and bottom are the maximum and minimum top-*L*, respectively.

**Figure 4.**
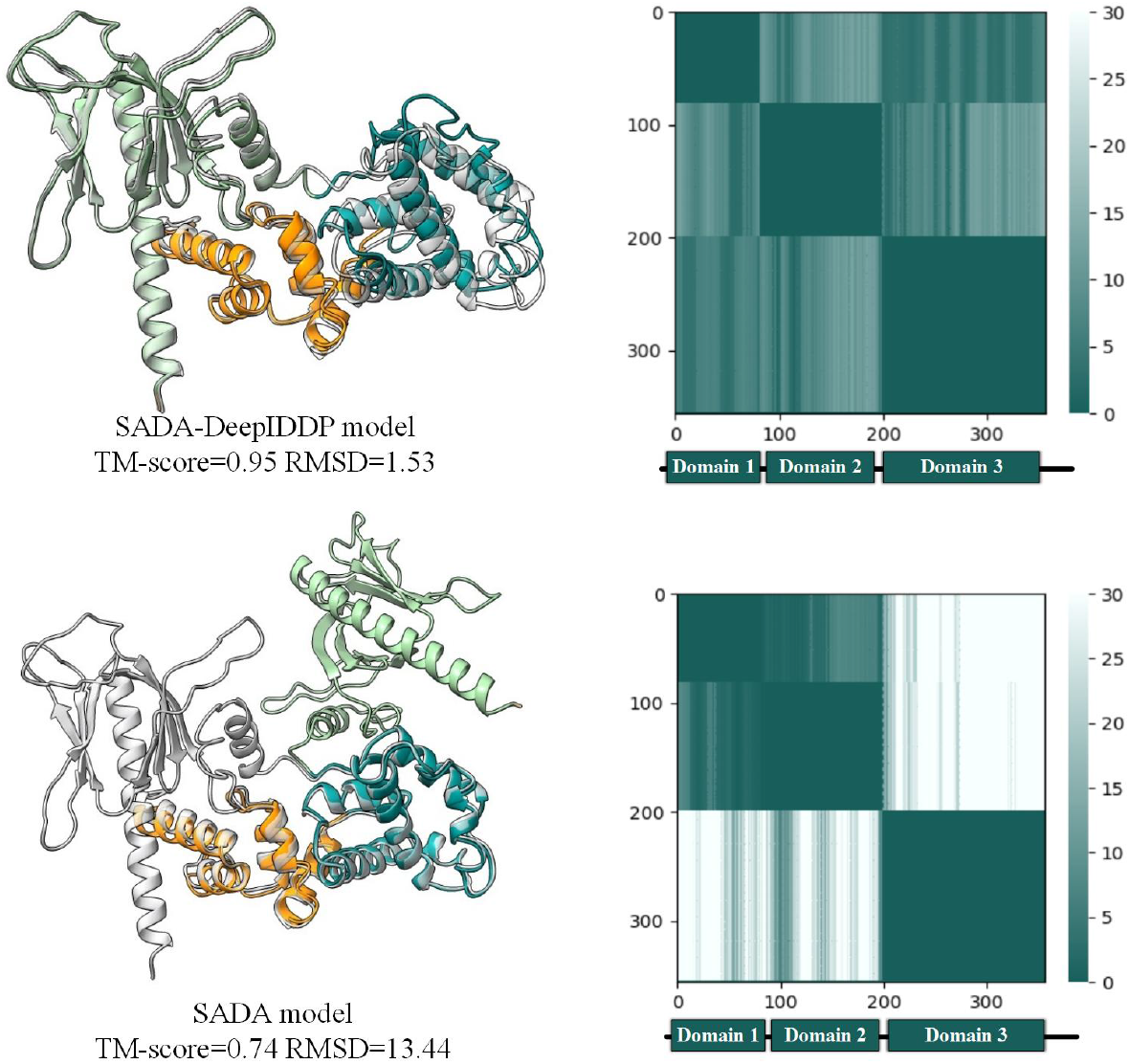
Modelling comparison of SADA-DeepIDDP and SADA on representative examples on the left side where grey represents the native structure, and various colours represent different domains. The corresponding PAE plot is shown on the right.

### 3.3 Compare with the state-of-art full-chain modeling method

There is still room to further improving the performance of AlphaFold2 for multi-domain proteins. We reassemble 68 human multi-domain proteins, which randomly selected from AlphaFold DB according to three standard criteria. The sequence identity with the DeepIDDP training set is ≤ 40%, the sequence identity with each other is ≤ 30% and the TM-score score is ≤ 0.80. Table 4 summarises the modelling results of SADA-DeepIDDP, SADA, and AlphaFold2. The detailed results are shown in Supplementary Table S5.

**Table 4.**
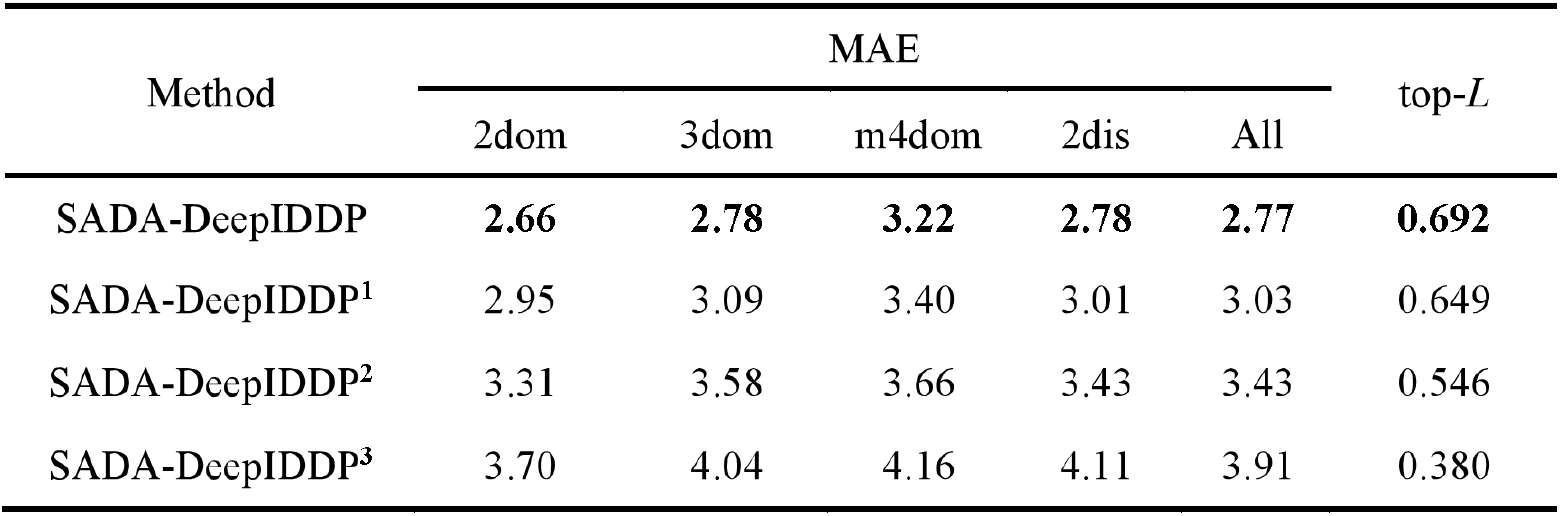
Summary of the results of inter-domain distance prediction for 356 multi-domain proteins.

| Method | MAE | | | | | top-$L$ |
| --- | --- | --- | --- | --- | --- | --- |
|  | 2dom | 3dom | m4dom | 2dis | All |  |
| SADA-DeepIDDP | **2.66** | **2.78** | **3.22** | **2.78** | **2.77** | **0.692** |
| SADA-DeepIDDP<sup>1</sup> | 2.95 | 3.09 | 3.40 | 3.01 | 3.03 | 0.649 |
| SADA-DeepIDDP<sup>2</sup> | 3.31 | 3.58 | 3.66 | 3.43 | 3.43 | 0.546 |
| SADA-DeepIDDP<sup>3</sup> | 3.70 | 4.04 | 4.16 | 4.11 | 3.91 | 0.380 |

Overall, the average TM-score of SADA-DeepIDDP model is 0.68, which is 13.33% and 3.03% higher than that of AlphaFold2(0.60) and SADA (0.66), respectively. The Student’s t-test *p*-values indicates statistically significant differences among the methods. The iRMSD of our method is 20.4% and 11.1% lower than that of AlphaFold2 and SADA, respectively, which indicates that DeepIDDP can further capture the interdomain orientations. Figure 5A shows the head-to-head comparison of SADA-DeepIDDP and AlphaFold2 on TM-score. Figure 5B is a representative three-domain example (PDBID:1dt9A), where the TM-scores for the models decomposed from the full-chain structures of AlphaFold2 for domains 1, 2 and 3 are 0.94, 0.95 and 0.94, respectively. However, because the inter-domain orientation is not correctly predicted, the TM-score of the full-length structure is only 0.80. The same problem occurs with the deep learning approach used in SADA beacuse it is not specifically processed and optimised for multi-domain proteins. DeepIDDP correctly predicts the inter-domain orientation and obtains a high-accuracy full-chain model (TM-score:0.95). Figure 5C is another example (PDBID:1dt9A) where the TM-scores of domains 1, 2 and 3 are 0.78, 0.92 and 0.77, respectively. However, the accuracy of Alphafold2 and SADA full-length models are only 0.63 and 0.60, respectively. The full-length model of SADA-DeepIDDP achieves a high accuracy (TM-score: 0.84).

**Figure 5.**
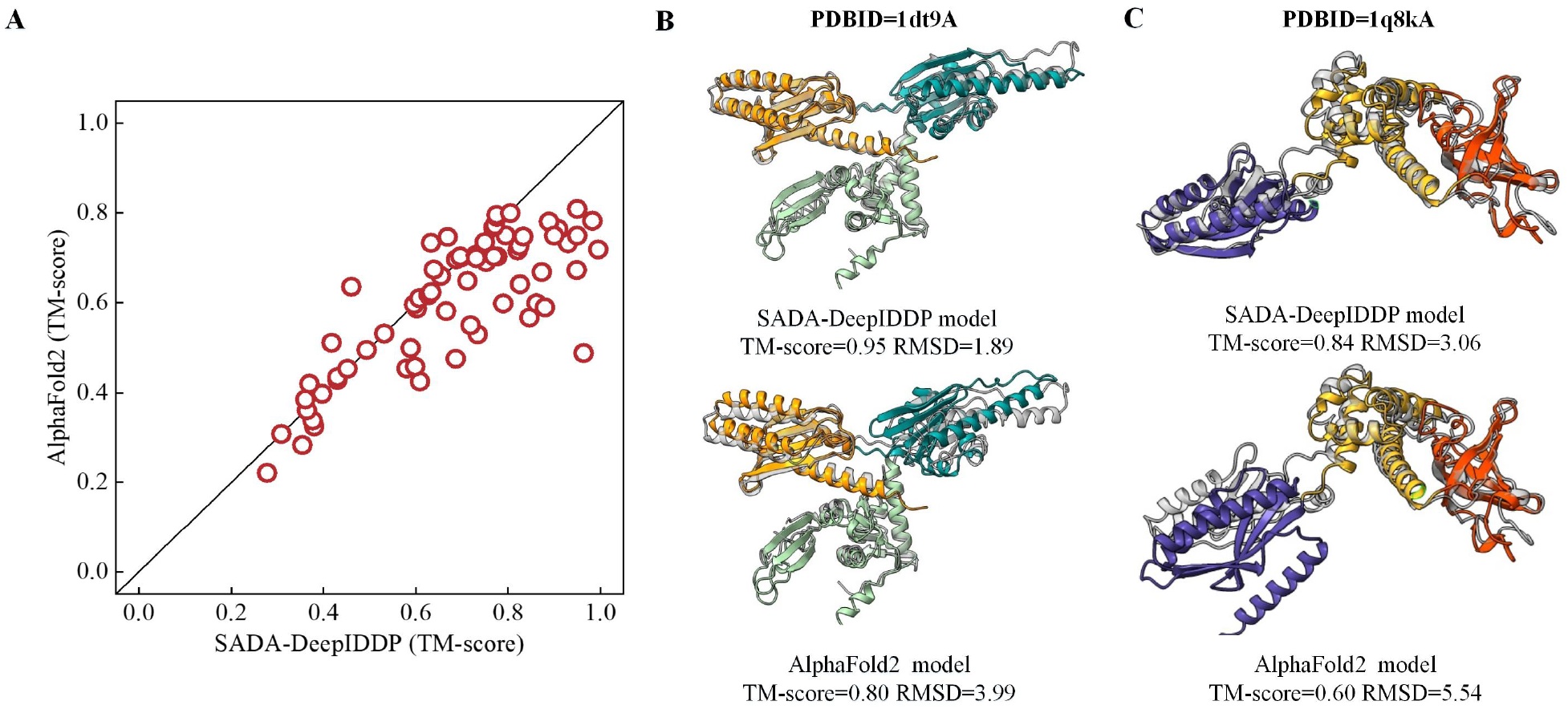
Comparison of SADA-DeepIDDP and AlphaFold2. **(A)** head-to-head TM-score comparison of models generated by SADA-DeepIDDP and that built by AlphaFold2 (**B** and **C)** representative examples are showing SADA-DeepIDDP builds better full-length models than SADA and AlphaFold2. Gray represents natural structures, and different colors represent different domains. **(B)**1dt9A **(C)** 1q8kA.

### 3.4 Performances on CASP14 targets

we also test the performance of SADA-DeepIDDP on 17 multi-domain proteins of CASP14 targets and compare it with AlphaFold2. AlphaFold2-CASP14 is the result of a download from the CASP website (http://predictioncenter.org). AlphaFold2-local is the result of executing of the localisation program, where the individual domain models used in SADA-DeepIDDP are decomposed from the full-chain model predicted by AlphaFold2-local. Figure 6 shows the TM-score for each method on each target, and the detailed results are listed in Supplementary Table S6. SADA-DeepIDDP achieves an average TM-score of 0.84, which is 2.44% and 2.44% higher than that of the original SADA (0.82) and AlphaFold2-CASP14 (0.82). When experimentally resolved individual domain models are input into SADA-DeepIDDP (SADA-DeepIDDP-native), the accuracy of the full-chain structure model increases to 0.88, which is 4.8% higher than that of SADA-DeepIDDP and 7.3% higher than that of AlphaFold2-CASP14.The results show that the assembly method can achieve higher accuracy models if accurate single-domain structures are available.

**Figure 6.**
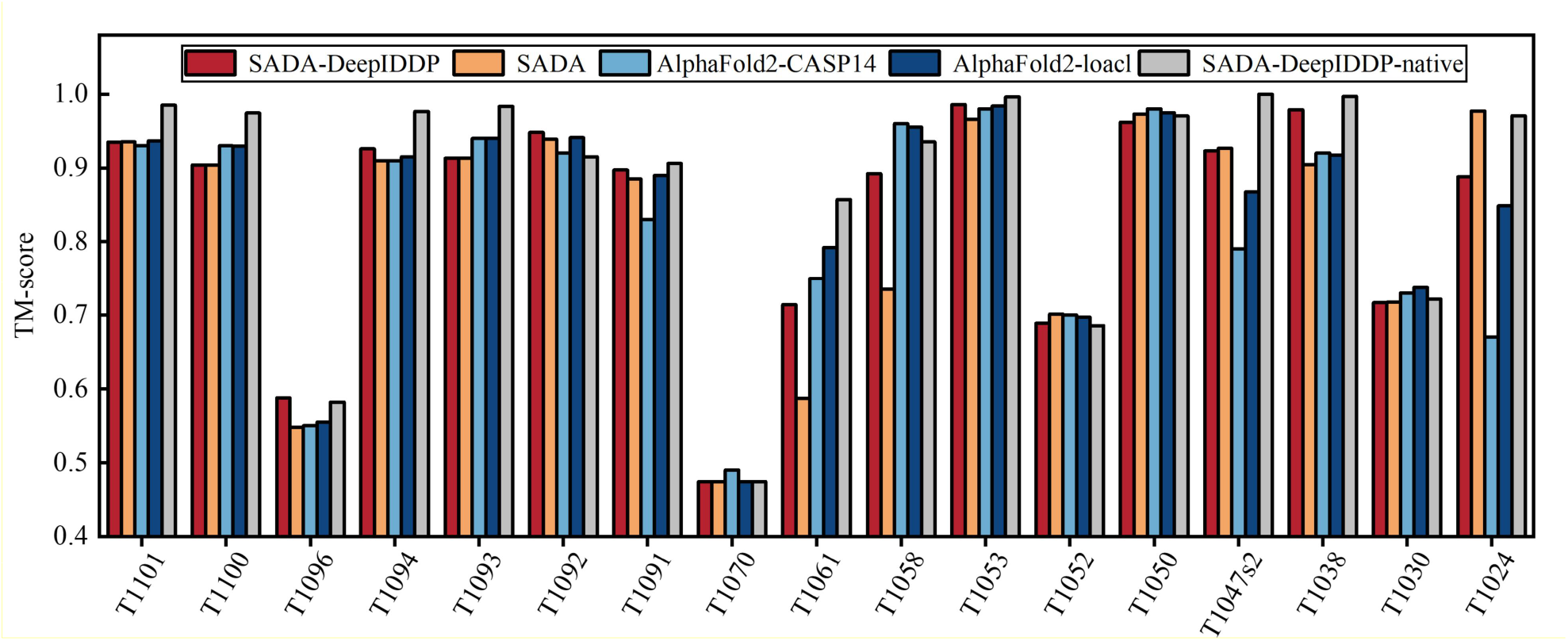
Histogram of TM-score for models predicted by SADA-DeepIDDP, SADA, AlphaFold2-CASP14, AlphaFold2-local and SADA-DeepIDDP-native. SADA-DeepIDDP-native represents using experimental domain.

### 3.5 Ablation experiment

To investigate the contribution of data enhancement and new features to our method. We design ablation experiments. SADA-DeepIDDP^3^ only uses basic MSA features. SADA-DeepIDDP^2^ added inter-domain features on the basis of SADA-DeepIDDP^3^. SADA-DeepIDDP^1^ uses three classes of features. SADA-DeepIDDP is the best version, using three classes of features and the DPMSA strategy. All the versions are trained using the same dataset and parameters. Table 4 summarises the comparative the results. The detailed results are listed in the Supplementary Table S7. The average MAE of SADA-DeepIDDP on all proteins is 2.77 Å, which is 8.4% lower than that of SADA-DeepIDDP^1^(3.03 Å). The average MAE value was reduced by 12.4% when the new features are added to the network (SADA-DeepIDDP^2^ vs. SADA-DeepIDDP^3^). In addition, the top-*L* metrics of SADA-DeepIDDP and SADA-DeepIDDP^2^ improved by 6.6% and 43.7% compared with SADA-DeepIDDP^1^ and SADA-DeepIDDP^3^. Among them, 216 and 261 proteins improved to different degrees, accounting for 60.67% and 73.31% of the total proteins, respectively. According to Figure 7, the average MAE value and the top-*L* metrics of SADA-DeepIDDP progressively improved as the designed inter-domain features and DPMSA strategy are gradually added to our network, which further demonstrates the effectiveness of the designed features and the rationality of the DPMSA strategy.

**Figure 7.**
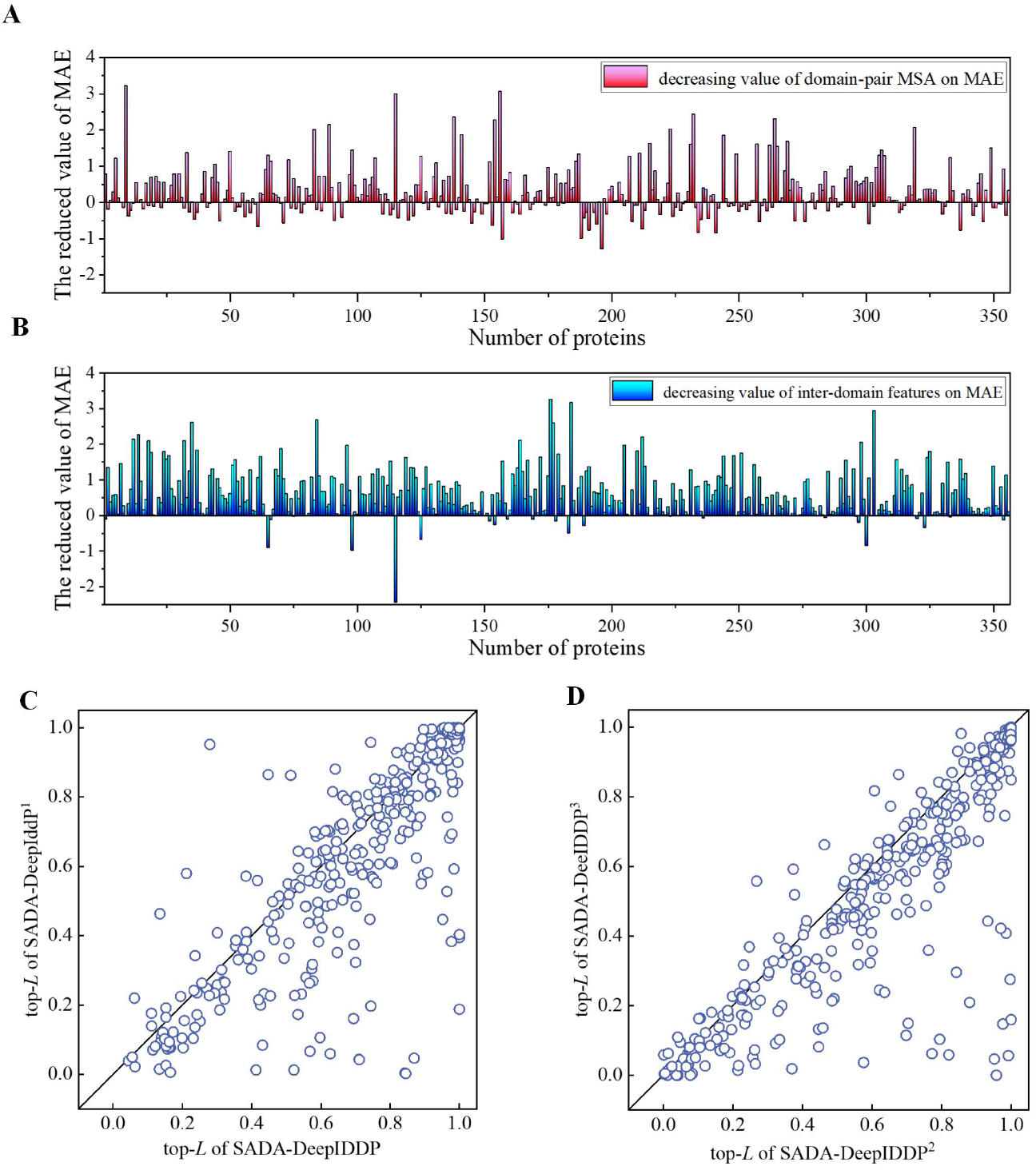
**(A)** Head-to-head top-*L* accuracy comparison of SADA-DeepIDDP and that created by SADA-DeepIDDP^**1**^. **(B)** Head-to-head top-*L* accuracy comparison of SADA-DeepIDDP^**2**^ and that created by SADA-DeepIDDP^**3**^. **(C)** Decreasing in MAE value for inter-domain residue pairs after domain-pair MSA data enhancement for each test protein. **(D)** Decreasing in MAE value for inter-domain residue pairs after the addition of inter-domain features for each test protein.

## 4. Conclusion

We developed an inter-domain distance prediction method DeepIDDP. In DeepIDDP, a non-redundant multi-domain training set was constructed using the CD-HIT and MPDB databases. Meanwhile, a deep residual network with attention mechanism was designed and two new inter-domain features were used to increase the ability to capture the distance of inter-domain residual pairs. Then, a data enhancement strategy called DPMSA was proposed for multi-domain proteins with poor homologous sequences. DeepIDDP outperformed existing distance prediction methods on a benchmark 356 multi-domain protein test set. In addition, the integration of DeepIDDP into our previously proposed assembly method SADA achieved a significant improvement over the original method. Finally, we also reassembled 68 human proteins with AlphaFold2 predictions ≤ 0.80. SADA-DeepIDDP obtained a high average model accuracy, especially for a large number of structural domains. These results indicated that DeepIDDP can effectively capture the residue pair distance relationships between structural domains and further improve the accuracy of multi-structural domain protein assembly methods. In the future, the idea of DeepIDDP can be extended to predict the inter-chain distance of protein complexes, which can help improve the accuracy of protein complex modelling. This is the next step in the research direction of our group.

## Key points

We developed an inter-domain distance prediction method (DeepIDDP). DeepIDDP predicts the distribution of inter-domain distances by constructing a training set of multi-domain proteins. Moreover, it is integrated into our domain assembly method SADA to further improve the accuracy of multi-domain proteins.

In DeepIDDP a network based on the attention mechanism is designed based on multi-domain proteins, and two new inter-domain features and data enhancement strategies are proposed to further improve the ability of the network to capture inter-domain distance.

Experimental results on 68 given human protein and 356 benchmark multi-domain test sets show that, our method can effectively capture the distance relationships of residue pairs between domains in the presence of a high number of domains. And it can be integrated into the domain assembly method to further improve the modelling accuracy of multi-domain models.

## Data and online server

All data needed to evaluate the conclusions are present in the paper and the Supplementary Materials. The online server at http://zhanglab-bioinf.com/DeepIDDP/. All the data utilized for training and testing DeepIDDP can be downloaded at this website.

## References

1. Pearce R, Zhang Y. Deep learning techniques have significantly impacted protein structure prediction and protein design, Curr Opin Struct Biol 2021;68:194–207.

2. Torrisi M, Pollastri G, Le Q. Deep learning methods in protein structure prediction, Comput Struct Biotechnol J 2020;18:1301–1310.

3. Pereira J, Simpkin AJ, Hartmann MD et al. High-accuracy protein structure prediction in CASP14, Proteins 2021;89:1687–1699.

4. AlQuraishi M. Machine learning in protein structure prediction, Curr Opin Chem Biol 2021;65:1–8.

5. Wang S, Sun S, Li Z et al. Accurate De Novo Prediction of Protein Contact Map by Ultra-Deep Learning Model, PLoS Comput Biol 2017;13:e1005324.

6. Jones DT, Buchan DW, Cozzetto D et al. PSICOV: precise structural contact prediction using sparse inverse covariance estimation on large multiple sequence alignments, Bioinformatics 2012;28:184–190.

7. Seemayer S, Gruber M, Soding J. CCMpred--fast and precise prediction of protein residue-residue contacts from correlated mutations, Bioinformatics 2014;30:3128–3130.

8. Rao R, Liu J, Verkuil R et al. MSA Transformer, International Conference on Machine Learning, Vol 139 2021;139.

9. Xu J. Distance-based protein folding powered by deep learning, Proc Natl Acad Sci U S A 2019;116:16856–16865.

10. Yang J, Anishchenko I, Park H et al. Improved protein structure prediction using predicted interresidue orientations, Proc Natl Acad Sci U S A 2020;117:1496–1503.

11. Su H, Wang W, Du Z et al. Improved Protein Structure Prediction Using a New Multi-Scale Network and Homologous Templates, Adv Sci (Weinh) 2021;8:e2102592.

12. Hou J, Wu T, Guo Z et al. The MULTICOM Protein Structure Prediction Server Empowered by Deep Learning and Contact Distance Prediction, Methods Mol Biol 2020;2165:13–26.

13. Shen T, Wu J, Lan H et al. When homologous sequences meet structural decoys: Accurate contact prediction by tFold in CASP14-(tFold for CASP14 contact prediction), Proteins 2021;89:1901–1910.

14. Kolodny R, Li Y, Zhang C et al. Deducing high-accuracy protein contact-maps from a triplet of coevolutionary matrices through deep residual convolutional networks, PLOS Computational Biology 2021;17.

15. Kandathil SM, Greener JG, Jones DT. Prediction of interresidue contacts with DeepMetaPSICOV in CASP13, Proteins 2019;87:1092–1099.

16. Baek M, DiMaio F, Anishchenko I et al. Accurate prediction of protein structures and interactions using a three-track neural network, Science 2021;373:871–876.

17. Jumper J, Evans R, Pritzel A et al. Highly accurate protein structure prediction with AlphaFold, Nature 2021;596:583–589.

18. Kryshtafovych A, Schwede T, Topf M et al. Critical assessment of methods of protein structure prediction (CASP)-Round XIV, Proteins 2021;89:1607–1617.

19. Jones DT, Thornton JM. The impact of AlphaFold2 one year on, Nature methods 2022;19:15.

20. Pearce R, Zhang Y. Toward the solution of the protein structure prediction problem, J Biol Chem 2021;297:100870.

21. Peng CX, Zhou XG, Xia YH et al. Structural analogue-based protein structure domain assembly assisted by deep learning, Bioinformatics 2022;38:4513–4521.

22. Jumper J, Hassabis D. Protein structure predictions to atomic accuracy with AlphaFold, Nat Methods 2022;19:11–12.

23. Xu D, Jaroszewski L, Li Z et al. AIDA: ab initio domain assembly for automated multi-domain protein structure prediction and domain-domain interaction prediction, Bioinformatics 2015;31:2098–2105.

24. Zhou X, Hu J, Zhang C et al. Assembling multidomain protein structures through analogous global structural alignments, Proceedings of the National Academy of Sciences 2019;116:15930–15938.

25. Liu J, He G-X, Zhao K-L et al. De novo protein structure prediction by incremental inter-residue geometries prediction and model quality assessment using deep learning 2022.

26. Fu L, Niu B, Zhu Z et al. CD-HIT: accelerated for clustering the next-generation sequencing data, Bioinformatics 2012;28:3150–3152.

27. Remmert M, Biegert A, Hauser A et al. HHblits: lightning-fast iterative protein sequence searching by HMM-HMM alignment, Nat Methods 2011;9:173–175.

28. Mirdita M, von den Driesch L, Galiez C et al. Uniclust databases of clustered and deeply annotated protein sequences and alignments, Nucleic Acids Res 2017;45:D170–D176.

29. Han JH, Batey S, Nickson AA et al. The folding and evolution of multidomain proteins, Nat Rev Mol Cell Biol 2007;8:319–330.

30. Kabsch W, Sander C. Dictionary of protein secondary structure: pattern recognition of hydrogen-bonded and geometrical features, Biopolymers 1983;22:2577–2637.

31. Leaver-Fay A, Tyka M, Lewis SM et al. ROSETTA3: an object-oriented software suite for the simulation and design of macromolecules, Methods Enzymol 2011;487:545–574.

32. Rohl CA, Strauss CE, Misura KM et al. Protein structure prediction using Rosetta, Methods Enzymol 2004;383:66–93.

33. Henikoff JG, Henikoff S. Using substitution probabilities to improve position-specific scoring matrices, Comput Appl Biosci 1996;12:135–143.

34. Ju F, Zhu J, Shao B et al. CopulaNet: Learning residue co-evolution directly from multiple sequence alignment for protein structure prediction, Nat Commun 2021;12:2535.

35. Xu J, Zhang Y. How significant is a protein structure similarity with TM-score = 0.5?, Bioinformatics 2010;26:889–895.

36. Hwang H, Pierce B, Mintseris J et al. Protein-protein docking benchmark version 3.0, Proteins 2008;73:705–709.

37. Kingsley LJ, Esquivel-Rodríguez J, Yang Y et al. Ranking protein–protein docking results using steered molecular dynamics and potential of mean force calculations, Journal of Computational Chemistry 2016;37:1861–1865.

38. Varadi M, Anyango S, Deshpande M et al. AlphaFold Protein Structure Database: massively expanding the structural coverage of protein-sequence space with high-accuracy models, Nucleic Acids Res 2022;50:D439–D444.

