## Supplemental Materials for "Inter-domain distance prediction based on deep learning for domain assembly"

**Equation S1.** Interface root mean deviation(iRMSD) is defined by

$$iRMSD = \sqrt{\frac{1}{2n}\sum_{i=0}^{2n}\left(x_i^* - x_i\right)^2} \qquad \text{(S1)}$$

Where $n$ is the number of residue pairs with the distance between domains less than 8Å in the ground truth. $x_i^* \in R^3$ and $x_i \in R^3$ is the coordinates of residue in the pairs corresponding to predicted structure and the ground truth after alignment [1].

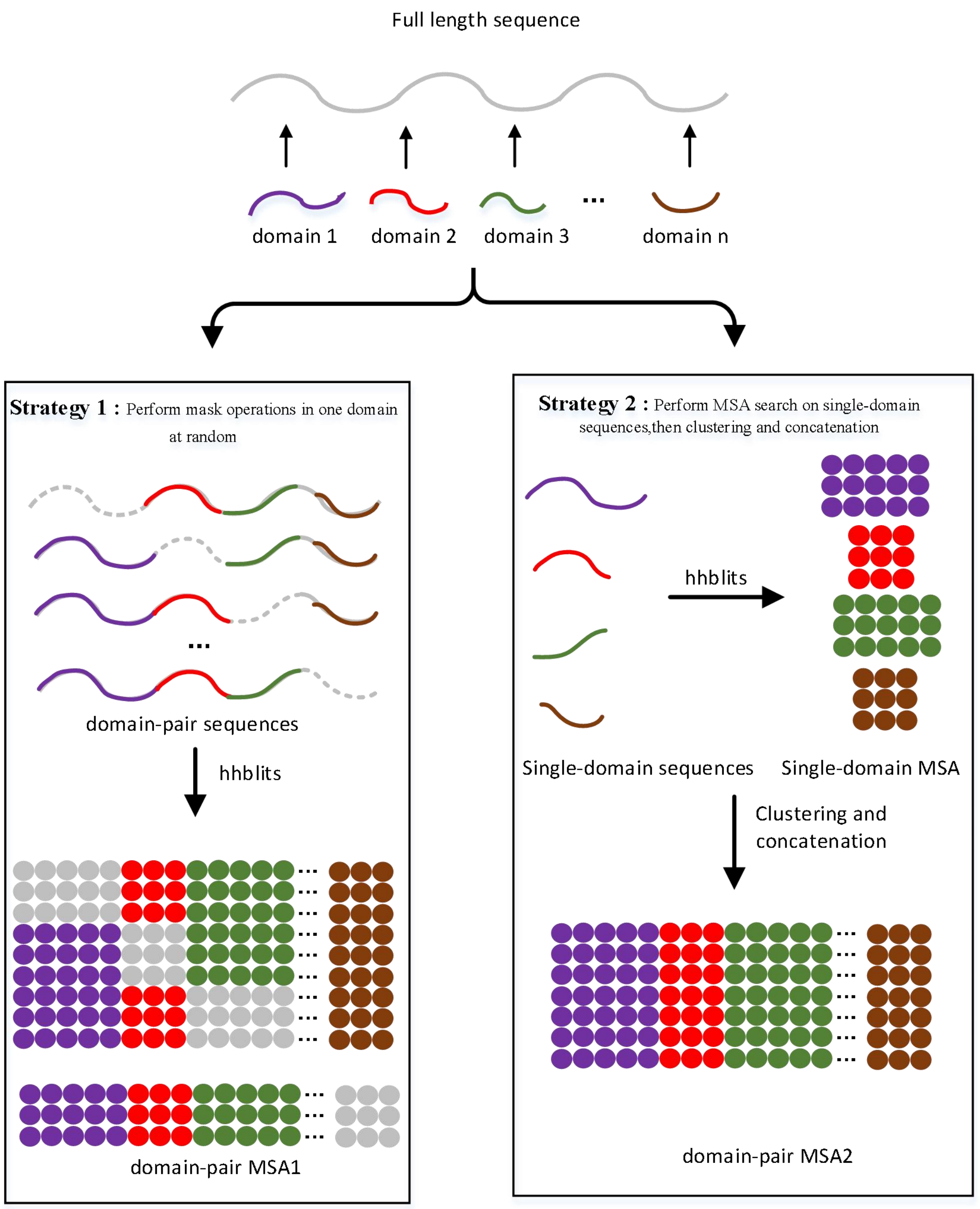


**Figure S1** DPMSA generation process.

**Table S1.** HHblits tool detailed parameters

| parameter | value | Description |
|---|---|---|
| *-id* | 90 | aximum pairwise sequence identity |
| *-cov* | 90、75、50 | minimum coverage with master sequence |
| *-e* | 1E-30、1E-10、1E-6、1E-3 | e-value cutoff for inclusion in result alignment |
| *-mact* | 0.35 | posterior prob threshold for MAC realignment controlling greedi- ness at alignment ends |
| *-neffmax* | 20 | skip further search iterations when diversity Neff of query MSA becomes larger than neffmax |
| *-n* | 4 | number of iterations |
| *-maxseq* | 7500、5000 | Number of sequence bars |
| *-d* | uniclust30、BFD | database name |

**Table S2.** All features used by DeepIDDP

| type | name | shape |
|---|---|---|
| inter-domain feature | IMCP | $L \times L \times 1$ |
| | ICOP | $L \times L \times 1$ |
| single-domain feature | domain information | $L \times 5$ |
| | secondary structure | $L \times 4$ |
| | rosetta enery term | $L \times L \times 13$ |
| MSA feature | target sequence | $L \times 21$ |
| | PSSM | $L \times 21$ |
| | position entropy | $L \times L \times 1$ |
| | inverse covariance matrix | $L \times L \times 441$ |

**Table S3.** Detailed results of DeepIDDP, SADA, trRosettaX, and trRosetta at inter-domain distances

| PDB ID | MAE | | | | Top-*L* | | | |
|---|---|---|---|---|---|---|---|---|
| | DeepIDDP | trRosettaX | trRosetta | SADA | DeepIDDP | trRosettaX | trRosetta | SADA |
| 1bf2A | 2.25 | 3.12 | 3.08 | 3.72 | 0.99 | 0.98 | 0.96 | 0.96 |
| 1bhgA | 2.76 | 3.46 | 3.30 | 3.68 | 0.98 | 0.98 | 0.97 | 0.96 |
| 1bp1A | 2.91 | 3.45 | 3.58 | 3.45 | 0.80 | 0.99 | 0.84 | 0.74 |
| 1c1zA | 1.96 | 3.08 | 2.58 | 3.08 | 0.88 | 0.90 | 0.89 | 0.87 |
| 1cjsA | 1.63 | 2.41 | 2.57 | 2.41 | 0.90 | 0.87 | 0.85 | 0.81 |
| 1cjyA | 4.02 | 4.20 | 4.90 | 4.18 | 0.14 | 0.17 | 0.17 | 0.13 |
| 1ck1A | 1.97 | 3.78 | 3.77 | 3.78 | 1.00 | 0.96 | 0.91 | 0.90 |
| 1d2pA | 3.66 | 4.11 | 2.79 | 4.16 | 0.76 | 0.52 | 0.61 | 0.53 |
| 1ecrA | 1.75 | 1.76 | 2.22 | 1.76 | 0.84 | 0.89 | 0.89 | 0.86 |
| 1efdN | 2.02 | 2.12 | 4.81 | 2.78 | 1.00 | 1.00 | 1.00 | 1.00 |
| 1f5qD | 2.04 | 2.89 | 2.99 | 2.89 | 1.00 | 0.96 | 0.99 | 0.99 |
| 1f7uA | 1.89 | 1.43 | 2.62 | 2.90 | 1.00 | 1.00 | 1.00 | 0.99 |
| 1fa9A | 2.68 | 3.80 | 3.31 | 3.80 | 0.83 | 0.77 | 0.71 | 0.76 |
| 1fjrA | 1.63 | 3.88 | 5.41 | 3.76 | 0.98 | 0.77 | 0.62 | 0.79 |
| 1fx7A | 2.09 | 3.05 | 3.30 | 3.24 | 1.00 | 0.99 | 1.00 | 0.98 |
| 1g87B | 3.61 | 4.36 | 4.45 | 4.35 | 0.48 | 0.57 | 0.29 | 0.18 |
| 1griA | 2.61 | 3.68 | 3.69 | 3.67 | 0.82 | 0.58 | 0.51 | 0.55 |
| 1gu7A | 1.58 | 2.01 | 1.83 | 2.01 | 0.93 | 0.95 | 0.98 | 0.98 |
| 1h88C | 1.23 | 2.49 | 2.56 | 1.90 | 0.99 | 0.99 | 0.96 | 0.93 |
| 1hx6B | 4.44 | 4.41 | 2.36 | 4.40 | 0.06 | 0.04 | 0.11 | 0.09 |
| 1itwA | 3.01 | 3.65 | 3.51 | 3.65 | 0.64 | 0.73 | 0.50 | 0.55 |
| 1iwaA | 2.89 | 4.29 | 4.60 | 3.99 | 0.69 | 0.64 | 0.58 | 0.55 |
| 1jkiA | 2.86 | 3.58 | 3.14 | 3.58 | 0.83 | 0.99 | 0.74 | 0.78 |
| 1k7tA | 1.65 | 3.55 | 4.17 | 3.49 | 0.98 | 0.75 | 0.74 | 0.71 |
| 1kfqA | 2.26 | 3.07 | 3.73 | 3.13 | 1.00 | 1.00 | 0.99 | 0.99 |
| 1ldjA | 2.85 | 2.60 | 2.46 | 2.97 | 0.96 | 0.98 | 0.98 | 0.97 |
| 1m5qH | 2.56 | 3.49 | 2.50 | 3.51 | 0.59 | 0.50 | 0.45 | 0.41 |
| 1m8pB | 2.04 | 3.42 | 2.85 | 3.36 | 0.99 | 0.97 | 0.97 | 0.97 |
| 1mkfA | 3.63 | 4.22 | 4.36 | 4.22 | 0.62 | 0.15 | 0.13 | 0.19 |
| 1mkmB | 1.65 | 2.73 | 1.77 | 2.32 | 0.69 | 0.53 | 0.68 | 0.61 |
| 1n80A | 3.41 | 4.11 | 3.92 | 4.11 | 0.63 | 0.73 | 0.56 | 0.55 |
| 1nh2D | 1.74 | 3.22 | 4.90 | 3.21 | 0.83 | 0.41 | 0.78 | 0.66 |
| 1ni5A | 2.29 | 2.88 | 2.94 | 3.05 | 0.91 | 0.90 | 0.93 | 0.89 |
| 1nyqB | 2.27 | 2.84 | 3.87 | 2.99 | 0.80 | 0.82 | 0.79 | 0.75 |
| 1nzjA | 1.37 | 2.12 | 1.89 | 2.12 | 1.00 | 1.00 | 1.00 | 1.00 |
| 1pprM | 2.51 | 4.11 | 3.95 | 4.31 | 0.92 | 1.00 | 0.15 | 0.05 |
| 1prrA | 2.13 | 4.00 | 2.35 | 3.94 | 0.91 | 0.39 | 0.27 | 0.19 |
| 1q19A | 3.29 | 3.99 | 5.05 | 3.89 | 0.71 | 0.61 | 0.58 | 0.60 |
| 1q25A | 3.37 | 3.95 | 3.95 | 3.96 | 0.69 | 0.53 | 0.42 | 0.40 |
| 1qhdA | 3.70 | 4.69 | 4.67 | 4.69 | 0.70 | 0.28 | 0.18 | 0.20 |
| 1qwrA | 4.30 | 4.38 | 5.08 | 4.37 | 0.26 | 0.24 | 0.25 | 0.26 |
| 1qz9A | 1.72 | 3.09 | 3.03 | 3.09 | 0.99 | 0.97 | 0.98 | 0.97 |

| PDB ID | MAE | | | | Top-*L* | | | |
|---|---|---|---|---|---|---|---|---|
| | DeepIDDP | trRosettaX | trRosetta | SADA | DeepIDDP | trRosettaX | trRosetta | SADA |
| 1r71B | 0.86 | 1.63 | 1.67 | 1.52 | 1.00 | 1.00 | 1.00 | 1.00 |
| 1rh1A | 3.82 | 4.76 | 5.06 | 4.74 | 0.71 | 0.14 | 0.08 | 0.16 |
| 1rktA | 1.48 | 2.79 | 4.10 | 2.13 | 0.99 | 0.85 | 0.92 | 0.90 |
| 1s6lA | 2.81 | 2.95 | 2.73 | 3.34 | 0.81 | 0.64 | 0.51 | 0.50 |
| 1sb7B | 3.61 | 4.11 | 4.10 | 4.11 | 0.51 | 0.41 | 0.41 | 0.37 |
| 1sp3A | 3.61 | 4.24 | 3.12 | 4.42 | 0.92 | 0.96 | 0.91 | 0.91 |
| 1ug9A | 3.51 | 3.86 | 3.85 | 4.12 | 0.70 | 0.79 | 0.52 | 0.58 |
| 1uzjA | 2.44 | 4.15 | 4.15 | 4.18 | 0.98 | 0.86 | 0.85 | 0.86 |
| 1vk1A | 1.93 | 3.45 | 3.34 | 3.45 | 0.93 | 0.63 | 0.61 | 0.63 |
| 1vrmA | 1.53 | 2.11 | 2.10 | 2.11 | 1.00 | 0.99 | 0.99 | 1.00 |
| 1vz6A | 1.75 | 2.91 | 1.94 | 2.81 | 0.87 | 0.69 | 0.66 | 0.70 |
| 1w3aA | 2.87 | 3.58 | 4.01 | 3.57 | 0.63 | 0.35 | 0.30 | 0.34 |
| 1wv3A | 2.15 | 3.91 | 5.60 | 3.96 | 0.92 | 0.38 | 0.15 | 0.23 |
| 1x7pA | 1.56 | 2.66 | 1.90 | 2.73 | 0.95 | 0.98 | 0.98 | 0.94 |
| 1x9yA | 3.58 | 4.35 | 4.39 | 4.35 | 0.76 | 0.29 | 0.17 | 0.42 |
| 1xvuA | 2.63 | 3.09 | 2.97 | 3.09 | 0.30 | 0.33 | 0.27 | 0.27 |
| 1y11A | 3.06 | 3.05 | 1.92 | 3.13 | 0.20 | 0.19 | 0.32 | 0.26 |
| 1yiqA | 3.87 | 4.17 | 3.37 | 4.21 | 0.59 | 0.57 | 0.37 | 0.50 |
| 1yy3A | 3.68 | 3.64 | 3.58 | 3.64 | 0.38 | 0.32 | 0.34 | 0.35 |
| 1z1wA | 2.54 | 2.96 | 2.56 | 2.95 | 0.94 | 0.96 | 0.97 | 0.98 |
| 1z87A | 2.96 | 3.00 | 3.10 | 3.00 | 0.25 | 0.20 | 0.16 | 0.17 |
| 1zbuB | 3.09 | 3.72 | 4.40 | 3.78 | 0.44 | 0.18 | 0.17 | 0.12 |
| 1ze1A | 1.71 | 2.77 | 2.85 | 2.75 | 0.79 | 0.78 | 0.72 | 0.76 |
| 1zpuA | 2.30 | 3.41 | 3.27 | 3.85 | 0.99 | 0.99 | 0.96 | 0.98 |
| 1zy9A | 3.77 | 3.98 | 4.14 | 4.23 | 0.59 | 0.74 | 0.52 | 0.51 |
| 2a1sC | 2.23 | 3.23 | 2.78 | 3.23 | 0.32 | 0.31 | 0.32 | 0.30 |
| 2a3lA | 2.34 | 3.34 | 3.14 | 3.34 | 0.85 | 0.75 | 0.77 | 0.73 |
| 2ablA | 2.00 | 3.62 | 3.64 | 3.52 | 0.81 | 0.41 | 0.48 | 0.45 |
| 2ahvA | 3.02 | 3.43 | 4.55 | 4.15 | 0.77 | 0.82 | 0.72 | 0.77 |
| 2au3A | 2.25 | 2.70 | 2.77 | 2.87 | 0.90 | 0.99 | 0.98 | 0.99 |
| 2b5uA | 2.95 | 3.83 | 3.85 | 3.92 | 0.60 | 0.37 | 0.27 | 0.33 |
| 2bkpA | 2.82 | 4.05 | 2.55 | 3.96 | 0.66 | 0.40 | 0.22 | 0.26 |
| 2bt1A | 3.52 | 4.23 | 4.18 | 4.23 | 0.79 | 1.00 | 0.44 | 0.53 |
| 2bydA | 3.79 | 4.14 | 4.14 | 4.14 | 0.63 | 0.61 | 0.59 | 0.59 |
| 2c1yA | 2.20 | 3.02 | 3.77 | 3.38 | 0.59 | 0.52 | 0.44 | 0.42 |
| 2c43A | 2.40 | 3.15 | 3.12 | 3.15 | 0.85 | 0.87 | 0.85 | 0.86 |
| 2cxcA | 1.37 | 1.27 | 2.62 | 1.73 | 1.00 | 1.00 | 1.00 | 1.00 |
| 2d1cA | 3.44 | 3.61 | 3.76 | 3.73 | 0.35 | 0.31 | 0.24 | 0.22 |
| 2d7iA | 2.97 | 3.09 | 3.97 | 3.17 | 0.14 | 0.23 | 0.16 | 0.17 |
| 2dfyC | 2.40 | 3.72 | 3.63 | 3.72 | 0.84 | 0.49 | 0.51 | 0.51 |
| 2dlaA | 2.08 | 2.79 | 2.72 | 2.79 | 0.96 | 0.78 | 0.72 | 0.79 |
| 2e9hA | 1.56 | 2.56 | 2.07 | 2.92 | 0.99 | 0.98 | 0.99 | 0.94 |
| 2e9xB | 1.74 | 2.15 | 4.10 | 2.35 | 1.00 | 1.00 | 1.00 | 0.99 |
| 2evrA | 3.72 | 3.98 | 5.69 | 3.96 | 0.35 | 0.10 | 0.11 | 0.10 |
| 2ew9A | 3.19 | 3.94 | 2.81 | 3.98 | 0.58 | 0.46 | 0.48 | 0.53 |
| 2ewfA | 3.51 | 3.82 | 3.82 | 3.82 | 0.57 | 0.61 | 0.31 | 0.25 |

| PDB ID | MAE | | | | Top-*L* | | | |
|---|---|---|---|---|---|---|---|---|
| | DeepIDDP | trRosettaX | trRosetta | SADA | DeepIDDP | trRosettaX | trRosetta | SADA |
| 2fd5A | 1.28 | 1.81 | 2.06 | 1.82 | 1.00 | 1.00 | 1.00 | 1.00 |
| 2g3pA | 1.96 | 4.15 | 4.16 | 4.15 | 0.97 | 0.24 | 0.29 | 0.24 |
| 2gg6A | 2.50 | 3.16 | 3.07 | 3.16 | 0.89 | 0.95 | 0.91 | 0.85 |
| 2gh8A | 3.71 | 3.76 | 2.40 | 3.76 | 0.15 | 0.36 | 0.07 | 0.07 |
| 2gsyE | 3.54 | 4.09 | 4.10 | 4.09 | 0.56 | 0.24 | 0.18 | 0.18 |
| 2gt1A | 2.00 | 3.03 | 1.42 | 2.99 | 0.91 | 0.91 | 0.87 | 0.89 |
| 2gzaC | 1.98 | 3.22 | 2.06 | 2.83 | 0.95 | 0.89 | 0.84 | 0.77 |
| 2gzoA | 1.81 | 2.37 | 2.49 | 2.37 | 1.00 | 0.98 | 0.95 | 0.96 |
| 2hjqA | 1.74 | 2.51 | 1.98 | 2.48 | 0.24 | 0.25 | 0.21 | 0.30 |
| 2hwjA | 2.52 | 3.53 | 3.56 | 3.25 | 0.83 | 0.83 | 0.80 | 0.86 |
| 2ii2A | 2.88 | 3.87 | 3.46 | 3.71 | 0.78 | 0.68 | 0.69 | 0.71 |
| 2ijd1 | 2.74 | 2.60 | 3.69 | 2.75 | 0.11 | 0.14 | 0.10 | 0.08 |
| 2iu7A | 1.66 | 3.13 | 4.23 | 3.22 | 0.83 | 0.29 | 0.51 | 0.48 |
| 2iw2A | 3.19 | 3.40 | 4.68 | 3.81 | 0.56 | 0.60 | 0.53 | 0.46 |
| 2j2cA | 3.08 | 3.60 | 3.47 | 3.60 | 0.77 | 0.96 | 0.64 | 0.68 |
| 2jz4A | 2.68 | 2.65 | 4.94 | 2.65 | 0.14 | 0.09 | 0.08 | 0.08 |
| 2kdyA | 2.21 | 4.13 | 3.11 | 3.89 | 0.95 | 0.79 | 0.78 | 0.80 |
| 2kfwA | 2.18 | 2.40 | 2.43 | 2.40 | 0.56 | 0.71 | 0.57 | 0.58 |
| 2kn4A | 1.83 | 2.47 | 6.42 | 2.55 | 0.20 | 0.16 | 0.15 | 0.16 |
| 2l9yA | 2.17 | 3.69 | 3.64 | 3.69 | 0.74 | 0.63 | 0.40 | 0.56 |
| 2mbgA | 1.92 | 3.67 | 1.96 | 3.67 | 0.80 | 0.40 | 0.19 | 0.29 |
| 2nsfA | 1.59 | 2.12 | 3.99 | 2.12 | 0.99 | 1.00 | 0.96 | 0.94 |
| 2ntyB | 2.40 | 3.79 | 3.77 | 3.79 | 0.77 | 0.43 | 0.43 | 0.41 |
| 2nykA | 2.79 | 4.82 | 4.62 | 4.80 | 0.95 | 0.43 | 0.16 | 0.34 |
| 2o6yA | 2.22 | 2.83 | 2.08 | 2.86 | 0.84 | 0.74 | 0.82 | 0.68 |
| 2olsA | 3.59 | 3.36 | 2.90 | 3.63 | 0.61 | 0.87 | 0.73 | 0.75 |
| 2owbA | 1.24 | 2.30 | 3.16 | 2.04 | 1.00 | 1.00 | 1.00 | 1.00 |
| 2piaA | 2.44 | 2.90 | 2.88 | 2.90 | 0.98 | 0.99 | 0.98 | 0.99 |
| 2qfiA | 2.13 | 3.21 | 3.76 | 3.06 | 0.63 | 0.53 | 0.52 | 0.50 |
| 2qp2A | 3.87 | 3.86 | 3.61 | 3.83 | 0.04 | 0.05 | 0.06 | 0.09 |
| 2qygA | 1.83 | 2.46 | 4.62 | 2.89 | 0.86 | 0.74 | 0.63 | 0.59 |
| 2r3vA | 2.68 | 2.91 | 2.83 | 2.91 | 0.90 | 0.99 | 0.88 | 0.93 |
| 2r58A | 1.75 | 3.95 | 3.89 | 3.95 | 0.99 | 1.00 | 0.58 | 0.58 |
| 2r5wB | 2.45 | 3.83 | 2.31 | 3.87 | 0.85 | 0.75 | 0.73 | 0.70 |
| 2r7dA | 2.17 | 3.17 | 2.84 | 3.33 | 0.99 | 0.99 | 0.98 | 0.98 |
| 2ra1A | 3.94 | 3.57 | 4.59 | 4.61 | 0.70 | 0.99 | 0.53 | 0.50 |
| 2ra1A | 3.14 | 3.15 | 4.59 | 4.05 | 0.88 | 0.80 | 0.68 | 0.61 |
| 2uu7A | 2.27 | 2.65 | 5.18 | 2.87 | 0.99 | 0.93 | 0.97 | 0.94 |
| 2uwnA | 1.62 | 2.70 | 2.66 | 2.82 | 0.64 | 0.81 | 0.22 | 0.27 |
| 2v0nA | 3.51 | 4.02 | 4.01 | 4.00 | 0.97 | 0.94 | 0.47 | 0.46 |
| 2v5dA | 2.62 | 3.64 | 3.38 | 3.85 | 0.97 | 0.97 | 0.95 | 0.95 |
| 2vgmA | 2.48 | 2.74 | 2.71 | 2.84 | 0.95 | 0.68 | 0.22 | 0.25 |
| 2w4bA | 3.07 | 3.89 | 2.83 | 4.41 | 0.98 | 0.94 | 0.92 | 0.92 |
| 2w4mA | 1.64 | 2.27 | 2.34 | 2.27 | 0.54 | 0.35 | 0.40 | 0.33 |
| 2wqrB | 2.67 | 3.56 | 3.57 | 3.58 | 0.31 | 0.38 | 0.08 | 0.10 |
| 2x0cA | 2.21 | 3.43 | 3.66 | 3.43 | 0.94 | 0.87 | 0.89 | 1.00 |

| PDB ID | MAE | | | | Top-*L* | | | |
|---|---|---|---|---|---|---|---|---|
| | DeepIDDP | trRosettaX | trRosetta | SADA | DeepIDDP | trRosettaX | trRosetta | SADA |
| 2x7iA | 2.32 | 3.34 | 4.19 | 3.07 | 0.72 | 0.53 | 0.53 | 0.86 |
| 2x8kC | 2.61 | 3.23 | 2.67 | 3.09 | 0.21 | 0.68 | 0.67 | 0.53 |
| 2xt6A | 3.94 | 3.50 | 3.23 | 3.70 | 0.98 | 0.91 | 0.45 | 0.73 |
| 2y25B | 2.19 | 4.70 | 4.71 | 4.90 | 0.81 | 0.61 | 0.64 | 0.50 |
| 2y51A | 2.27 | 3.98 | 3.18 | 3.98 | 1.00 | 0.98 | 0.97 | 0.57 |
| 2yb0E | 1.37 | 2.68 | 2.70 | 2.68 | 0.69 | 0.35 | 0.52 | 0.96 |
| 2yilA | 2.28 | 3.93 | 4.04 | 3.90 | 0.60 | 0.73 | 0.74 | 0.52 |
| 2yk0A | 3.82 | 3.88 | 3.78 | 4.01 | 0.17 | 0.17 | 0.22 | 0.76 |
| 2yrqA | 2.72 | 2.82 | 4.97 | 2.64 | 0.40 | 0.13 | 0.12 | 0.17 |
| 2z86C | 4.05 | 4.09 | 4.09 | 4.09 | 0.76 | 0.93 | 0.91 | 0.12 |
| 2zpaB | 3.86 | 2.90 | 3.00 | 3.13 | 0.59 | 0.76 | 0.63 | 0.92 |
| 2zxcA | 3.49 | 3.48 | 2.48 | 3.91 | 0.60 | 0.30 | 0.28 | 0.59 |
| 2zzqA | 3.96 | 4.15 | 4.11 | 4.15 | 0.06 | 0.05 | 0.05 | 0.29 |
| 3a1iA | 3.13 | 3.17 | 5.28 | 3.14 | 0.92 | 0.86 | 0.80 | 0.05 |
| 3a45A | 2.40 | 4.25 | 4.14 | 3.96 | 0.24 | 0.12 | 0.08 | 0.64 |
| 3a56A | 3.76 | 3.86 | 4.55 | 3.79 | 0.11 | 0.09 | 0.10 | 0.08 |
| 3afoA | 4.55 | 4.56 | 4.58 | 4.56 | 0.74 | 0.64 | 0.61 | 0.09 |
| 3ajvA | 2.73 | 3.31 | 2.98 | 3.20 | 0.79 | 0.85 | 0.87 | 0.61 |
| 3apoA | 4.54 | 2.45 | 3.66 | 2.61 | 0.87 | 0.85 | 0.83 | 1.00 |
| 3aqkA | 1.64 | 2.46 | 2.67 | 1.86 | 0.62 | 0.51 | 0.61 | 0.84 |
| 3arbA | 2.03 | 3.40 | 2.66 | 3.47 | 1.00 | 0.96 | 0.99 | 0.51 |
| 3aujG | 1.60 | 2.76 | 4.11 | 2.46 | 0.82 | 0.58 | 0.42 | 0.99 |
| 3b2zF | 3.10 | 4.00 | 3.25 | 4.00 | 0.21 | 0.22 | 0.22 | 0.25 |
| 3b43A | 3.07 | 3.24 | 2.48 | 3.58 | 0.28 | 0.60 | 0.62 | 0.22 |
| 3b7wA | 4.31 | 3.86 | 3.25 | 4.06 | 0.92 | 0.45 | 0.51 | 0.92 |
| 3bt1U | 2.76 | 4.22 | 4.22 | 4.22 | 1.00 | 0.97 | 1.00 | 0.44 |
| 3bt3A | 1.25 | 2.55 | 4.04 | 2.36 | 0.54 | 0.40 | 0.45 | 0.95 |
| 3bu2A | 2.63 | 2.49 | 2.58 | 2.49 | 0.97 | 0.95 | 0.88 | 0.45 |
| 3c1yA | 1.85 | 3.65 | 3.62 | 3.98 | 0.53 | 0.46 | 0.46 | 0.91 |
| 3c4tA | 2.93 | 3.46 | 2.65 | 3.59 | 0.54 | 0.41 | 0.47 | 0.46 |
| 3craA | 1.90 | 3.31 | 1.96 | 2.98 | 0.93 | 0.46 | 0.40 | 0.42 |
| 3cvzA | 2.38 | 4.35 | 4.32 | 4.35 | 0.95 | 0.91 | 0.60 | 0.51 |
| 3cw2C | 1.73 | 2.82 | 3.39 | 3.35 | 0.81 | 0.54 | 0.51 | 0.57 |
| 3d30A | 3.23 | 4.48 | 3.96 | 4.40 | 0.71 | 0.46 | 0.40 | 0.54 |
| 3dupA | 3.74 | 4.14 | 4.16 | 4.14 | 0.77 | 0.57 | 0.53 | 0.42 |
| 3eo5A | 2.90 | 4.02 | 2.87 | 3.96 | 0.56 | 0.50 | 0.48 | 0.46 |
| 3errA | 3.20 | 3.81 | 4.49 | 3.82 | 0.85 | 0.94 | 0.72 | 0.49 |
| 3eswA | 2.08 | 3.02 | 2.68 | 3.02 | 0.57 | 0.83 | 0.69 | 0.80 |
| 3eukH | 4.07 | 4.32 | 3.93 | 4.32 | 0.58 | 0.50 | 0.20 | 0.77 |
| 3f83A | 3.50 | 4.22 | 4.23 | 4.26 | 0.89 | 0.60 | 0.61 | 0.39 |
| 3fc3A | 2.26 | 4.19 | 4.22 | 4.21 | 1.00 | 1.00 | 1.00 | 0.43 |
| 3fi7A | 1.29 | 1.73 | 2.20 | 1.73 | 1.00 | 1.00 | 1.00 | 0.98 |
| 3fvvA | 1.14 | 1.52 | 1.76 | 1.52 | 0.75 | 0.75 | 0.75 | 1.00 |
| 3g79A | 2.29 | 2.57 | 2.37 | 2.65 | 1.00 | 0.97 | 0.96 | 0.73 |
| 3gbgA | 2.40 | 3.46 | 3.38 | 3.38 | 0.66 | 0.77 | 0.64 | 0.97 |
| 3gf5B | 3.32 | 2.76 | 3.46 | 2.95 | 0.60 | 0.60 | 0.58 | 0.59 |

| PDB ID | MAE | | | | Top-*L* | | | |
|---|---|---|---|---|---|---|---|---|
| | DeepIDDP | trRosettaX | trRosetta | SADA | DeepIDDP | trRosettaX | trRosetta | SADA |
| 3gmsA | 2.96 | 3.53 | 3.44 | 3.53 | 0.42 | 0.78 | 0.62 | 0.56 |
| 3h2tA | 3.39 | 3.41 | 3.64 | 3.78 | 1.00 | 1.00 | 1.00 | 0.59 |
| 3h5cB | 2.59 | 3.80 | 3.16 | 3.08 | 0.79 | 0.84 | 0.81 | 1.00 |
| 3hcsA | 1.82 | 1.27 | 4.99 | 1.31 | 0.36 | 0.16 | 0.14 | 0.78 |
| 3hjlA | 2.18 | 3.76 | 4.22 | 3.73 | 0.06 | 0.09 | 0.09 | 0.17 |
| 3hyiA | 3.40 | 3.39 | 3.00 | 3.36 | 1.00 | 0.98 | 0.98 | 0.09 |
| 3hzzB | 1.54 | 2.53 | 2.38 | 2.53 | 0.67 | 0.62 | 0.51 | 0.96 |
| 3i2dA | 2.80 | 3.25 | 3.84 | 3.32 | 0.85 | 0.85 | 0.92 | 0.55 |
| 3iam2 | 2.51 | 2.49 | 3.19 | 2.76 | 0.61 | 0.69 | 0.71 | 0.84 |
| 3ibjA | 3.03 | 3.19 | 2.72 | 2.83 | 0.95 | 0.98 | 0.98 | 0.66 |
| 3ifrA | 2.44 | 2.80 | 1.73 | 2.80 | 0.72 | 0.69 | 0.67 | 0.97 |
| 3ippB | 3.27 | 4.00 | 3.93 | 3.97 | 0.61 | 0.60 | 0.59 | 0.68 |
| 3isqA | 3.57 | 4.03 | 3.98 | 3.90 | 0.98 | 0.99 | 0.98 | 0.59 |
| 3j7aK | 1.51 | 3.62 | 2.13 | 3.04 | 0.64 | 0.78 | 0.63 | 0.99 |
| 3jymB | 2.71 | 3.48 | 3.24 | 3.27 | 0.45 | 0.86 | 0.34 | 0.71 |
| 3k1rA | 3.51 | 3.56 | 2.26 | 3.89 | 0.59 | 0.49 | 0.41 | 0.35 |
| 3k2iA | 3.74 | 3.98 | 3.02 | 4.13 | 0.94 | 0.87 | 0.83 | 0.50 |
| 3kbgA | 2.72 | 3.55 | 3.53 | 3.55 | 0.96 | 0.85 | 0.83 | 0.87 |
| 3kh5A | 2.36 | 3.68 | 2.41 | 3.92 | 0.75 | 0.34 | 0.35 | 0.84 |
| 3kjpA | 2.32 | 3.70 | 2.49 | 3.64 | 0.60 | 0.52 | 0.55 | 0.31 |
| 3kq4B | 3.48 | 4.10 | 3.05 | 4.03 | 0.48 | 0.24 | 0.19 | 0.46 |
| 3kt1A | 3.89 | 4.16 | 3.80 | 4.19 | 0.94 | 0.64 | 0.62 | 0.16 |
| 3ktmE | 1.59 | 4.16 | 2.54 | 3.98 | 0.57 | 0.47 | 0.50 | 0.58 |
| 3kwlA | 3.43 | 3.19 | 2.92 | 3.87 | 0.99 | 0.95 | 0.92 | 0.47 |
| 3kzwA | 1.27 | 2.68 | 3.60 | 2.74 | 0.12 | 0.09 | 0.10 | 0.94 |
| 3l76A | 3.84 | 3.90 | 2.35 | 3.88 | 0.63 | 0.33 | 0.24 | 0.10 |
| 3ld1A | 3.33 | 4.44 | 3.43 | 4.46 | 0.98 | 1.00 | 0.98 | 0.32 |
| 3lsgA | 1.61 | 1.56 | 2.21 | 1.75 | 0.20 | 0.16 | 0.17 | 0.99 |
| 3m1uA | 4.25 | 4.47 | 4.92 | 4.47 | 0.98 | 0.95 | 0.98 | 0.14 |
| 3mc8A | 1.92 | 3.50 | 2.92 | 2.97 | 0.99 | 0.90 | 0.82 | 0.96 |
| 3me4A | 1.55 | 2.34 | 2.42 | 2.74 | 0.90 | 0.66 | 0.68 | 0.86 |
| 3ml4C | 2.32 | 4.19 | 4.28 | 4.12 | 0.74 | 0.22 | 0.25 | 0.71 |
| 3mw8A | 3.15 | 4.91 | 4.43 | 4.91 | 0.27 | 0.28 | 0.29 | 0.23 |
| 3mwcA | 4.14 | 4.40 | 4.40 | 4.40 | 0.85 | 0.52 | 0.43 | 0.30 |
| 3mx2B | 2.64 | 3.46 | 3.38 | 4.09 | 0.13 | 0.11 | 0.13 | 0.59 |
| 3mzfA | 4.23 | 4.29 | 4.74 | 4.31 | 0.88 | 0.25 | 0.55 | 0.14 |
| 3njaB | 2.26 | 3.41 | 3.64 | 3.06 | 0.54 | 0.27 | 0.32 | 0.17 |
| 3npfA | 3.79 | 4.21 | 4.21 | 4.18 | 0.53 | 0.47 | 0.46 | 0.28 |
| 3nqiA | 3.09 | 3.81 | 2.23 | 3.77 | 0.38 | 0.34 | 0.26 | 0.44 |
| 3nsjA | 4.06 | 4.35 | 2.21 | 4.35 | 0.16 | 0.06 | 0.06 | 0.22 |
| 3nt8A | 3.29 | 3.41 | 3.74 | 3.41 | 0.99 | 1.00 | 0.99 | 0.11 |
| 3ntkA | 1.64 | 1.75 | 2.93 | 1.75 | 1.00 | 1.00 | 0.99 | 0.99 |
| 3oaaG | 2.58 | 2.81 | 2.75 | 2.81 | 0.92 | 0.98 | 0.93 | 0.99 |
| 3ob8A | 3.32 | 3.89 | 3.21 | 4.02 | 0.99 | 0.99 | 1.00 | 0.94 |
| 3og5A | 2.07 | 2.38 | 2.36 | 2.32 | 0.45 | 0.54 | 0.26 | 0.99 |
| 3oh0A | 4.16 | 4.21 | 3.34 | 4.41 | 0.92 | 0.83 | 0.43 | 0.28 |

| PDB ID | MAE | | | | Top-*L* | | | |
|---|---|---|---|---|---|---|---|---|
| | DeepIDDP | trRosettaX | trRosetta | SADA | DeepIDDP | trRosettaX | trRosetta | SADA |
| 3opfB | 3.04 | 4.10 | 2.69 | 4.14 | 0.76 | 0.81 | 0.58 | 0.33 |
| 3orjA | 3.06 | 3.82 | 3.71 | 3.79 | 0.47 | 0.28 | 0.14 | 0.59 |
| 3p53A | 3.98 | 4.00 | 2.50 | 3.86 | 0.59 | 0.30 | 0.25 | 0.15 |
| 3pcsB | 3.63 | 4.04 | 3.65 | 4.05 | 0.98 | 0.98 | 0.98 | 0.26 |
| 3plaA | 1.63 | 2.54 | 2.27 | 2.22 | 0.74 | 0.90 | 0.90 | 0.96 |
| 3po3S | 2.86 | 1.64 | 2.58 | 1.72 | 0.83 | 0.80 | 0.80 | 0.91 |
| 3ptyA | 3.00 | 3.66 | 3.72 | 3.79 | 0.77 | 0.74 | 0.64 | 0.73 |
| 3pxpA | 4.15 | 4.16 | 4.33 | 4.16 | 0.14 | 0.20 | 0.35 | 0.62 |
| 3qavA | 1.78 | 1.64 | 1.25 | 1.77 | 0.97 | 1.00 | 1.00 | 0.33 |
| 3qe9Y | 2.20 | 1.51 | 3.31 | 3.66 | 1.00 | 1.00 | 0.95 | 1.00 |
| 3qf4B | 2.37 | 2.91 | 4.40 | 3.06 | 0.31 | 0.38 | 0.30 | 0.99 |
| 3qjjA | 1.84 | 1.34 | 3.82 | 3.35 | 1.00 | 1.00 | 0.99 | 0.28 |
| 3qjoA | 3.28 | 2.98 | 3.85 | 4.01 | 0.92 | 0.99 | 0.80 | 0.98 |
| 3qphA | 2.57 | 3.80 | 3.81 | 3.83 | 0.81 | 0.76 | 0.42 | 0.81 |
| 3qtdA | 3.05 | 3.00 | 5.24 | 3.45 | 0.86 | 0.94 | 0.93 | 0.51 |
| 3qyeA | 2.59 | 3.45 | 2.95 | 2.99 | 0.99 | 0.98 | 1.00 | 0.94 |
| 3r05A | 4.96 | 3.51 | 1.67 | 3.72 | 0.95 | 0.97 | 0.95 | 0.99 |
| 3r6bA | 1.66 | 2.16 | 3.10 | 2.70 | 0.57 | 0.63 | 0.63 | 0.95 |
| 3rfyA | 1.99 | 3.52 | 3.41 | 3.59 | 0.85 | 0.73 | 0.71 | 0.58 |
| 3rh7A | 3.07 | 4.00 | 2.97 | 4.00 | 0.74 | 0.49 | 0.49 | 0.71 |
| 3rimA | 3.49 | 3.75 | 3.57 | 3.93 | 0.67 | 0.68 | 0.65 | 0.50 |
| 3rrpA | 2.10 | 2.89 | 2.57 | 2.82 | 0.94 | 0.93 | 1.00 | 0.69 |
| 3rwxA | 2.53 | 3.63 | 4.24 | 3.62 | 0.43 | 0.12 | 0.15 | 0.97 |
| 3sb4A | 3.02 | 3.81 | 4.19 | 3.83 | 0.56 | 0.16 | 0.24 | 0.12 |
| 3seoB | 2.80 | 3.84 | 3.91 | 3.96 | 0.65 | 0.34 | 0.38 | 0.29 |
| 3soaA | 2.33 | 2.92 | 2.72 | 2.79 | 0.98 | 0.94 | 0.94 | 0.31 |
| 3spgA | 2.44 | 2.09 | 3.07 | 3.06 | 0.31 | 0.36 | 0.31 | 0.89 |
| 3swjA | 2.82 | 4.11 | 3.83 | 4.11 | 0.72 | 0.54 | 0.46 | 0.31 |
| 3t58B | 4.28 | 3.91 | 4.24 | 4.29 | 0.32 | 0.49 | 0.15 | 0.30 |
| 3t7jA | 1.21 | 2.95 | 3.72 | 3.90 | 1.00 | 0.97 | 0.95 | 0.23 |
| 3tixD | 3.12 | 4.30 | 4.25 | 4.29 | 0.57 | 0.52 | 0.27 | 0.92 |
| 3tp9A | 2.52 | 3.04 | 2.73 | 3.03 | 0.89 | 0.95 | 0.90 | 0.42 |
| 3u07C | 2.73 | 3.85 | 1.47 | 3.81 | 0.79 | 0.75 | 0.75 | 0.92 |
| 3u0kA | 3.93 | 4.22 | 4.26 | 4.24 | 0.30 | 0.18 | 0.15 | 0.70 |
| 3u0oB | 2.08 | 3.04 | 4.80 | 3.09 | 0.97 | 0.87 | 0.86 | 0.14 |
| 3u9gA | 3.06 | 4.60 | 5.15 | 4.52 | 0.98 | 0.84 | 0.72 | 0.87 |
| 3ua3A | 3.46 | 4.12 | 3.71 | 4.05 | 0.81 | 0.73 | 0.77 | 0.68 |
| 3ub1D | 3.00 | 4.32 | 3.23 | 4.70 | 0.52 | 0.36 | 0.31 | 0.75 |
| 3ubhA | 2.35 | 3.14 | 1.81 | 3.15 | 0.70 | 0.60 | 0.59 | 0.27 |
| 3uitD | 1.95 | 2.63 | 3.37 | 2.67 | 0.23 | 0.12 | 0.11 | 0.59 |
| 3uj0A | 3.06 | 3.49 | 3.34 | 3.48 | 0.63 | 0.44 | 0.47 | 0.11 |
| 3uo3A | 2.84 | 4.30 | 3.43 | 4.17 | 0.66 | 0.34 | 0.33 | 0.45 |
| 3v7oB | 1.56 | 2.61 | 3.61 | 2.21 | 0.23 | 0.18 | 0.25 | 0.32 |
| 3vlaA | 4.48 | 4.54 | 4.61 | 4.65 | 0.41 | 0.14 | 0.20 | 0.25 |
| 3vn4A | 3.26 | 3.83 | 3.98 | 4.09 | 0.85 | 0.72 | 0.68 | 0.18 |
| 3vr8B | 2.20 | 3.68 | 1.45 | 2.74 | 0.96 | 0.87 | 0.94 | 0.64 |

| PDB ID | MAE | | | | Top-*L* | | | |
|---|---|---|---|---|---|---|---|---|
| | DeepIDDP | trRosettaX | trRosetta | SADA | DeepIDDP | trRosettaX | trRosetta | SADA |
| 3vsmA | 3.92 | 4.43 | 4.37 | 4.41 | 0.53 | 0.66 | 0.29 | 0.32 |
| 3vstA | 3.60 | 3.91 | 3.78 | 3.88 | 0.62 | 0.53 | 0.53 | 0.52 |
| 3w1bA | 3.55 | 3.85 | 3.85 | 3.99 | 0.47 | 0.50 | 0.42 | 0.42 |
| 3w2wA | 3.32 | 3.46 | 1.88 | 3.95 | 0.51 | 0.89 | 0.77 | 0.85 |
| 3wkuA | 2.00 | 3.67 | 2.34 | 3.69 | 0.62 | 0.44 | 0.41 | 0.16 |
| 3zh9B | 2.54 | 3.16 | 3.11 | 3.09 | 0.82 | 0.83 | 0.80 | 0.79 |
| 3zniA | 2.95 | 4.03 | 2.19 | 4.07 | 0.85 | 0.70 | 0.33 | 0.28 |
| 3zvmA | 3.25 | 3.41 | 4.46 | 3.44 | 0.24 | 0.35 | 0.33 | 0.24 |
| 4acoA | 4.10 | 4.16 | 2.06 | 4.17 | 0.15 | 0.09 | 0.11 | 0.09 |
| 4aimA | 3.26 | 3.22 | 1.73 | 3.50 | 0.65 | 0.69 | 0.69 | 0.76 |
| 4ak1A | 4.08 | 4.21 | 2.94 | 4.24 | 0.24 | 0.08 | 0.09 | 0.10 |
| 4alzA | 3.25 | 4.35 | 4.34 | 4.35 | 0.57 | 0.26 | 0.26 | 0.26 |
| 4ap5A | 2.80 | 3.36 | 2.25 | 3.83 | 0.70 | 0.74 | 0.63 | 0.62 |
| 4aq1A | 3.97 | 4.05 | 2.08 | 4.08 | 0.16 | 0.10 | 0.12 | 0.15 |
| 4aqfB | 3.52 | 4.26 | 4.24 | 4.26 | 0.65 | 0.31 | 0.24 | 0.26 |
| 4ax8A | 3.85 | 3.85 | 3.97 | 4.01 | 0.15 | 0.23 | 0.24 | 0.21 |
| 4axdA | 3.71 | 3.84 | 2.84 | 4.16 | 0.58 | 0.68 | 0.55 | 0.50 |
| 4b21A | 2.70 | 3.62 | 3.64 | 3.55 | 0.71 | 0.43 | 0.47 | 0.46 |
| 4b3iA | 3.62 | 3.78 | 3.68 | 3.79 | 0.46 | 0.47 | 0.47 | 0.46 |
| 4bd9B | 1.90 | 3.31 | 3.99 | 3.75 | 0.99 | 1.00 | 0.98 | 0.93 |
| 4bfiB | 1.64 | 2.49 | 2.99 | 2.51 | 0.88 | 0.78 | 0.78 | 0.80 |
| 4bt9B | 2.09 | 2.21 | 3.17 | 2.12 | 0.17 | 0.17 | 0.19 | 0.18 |
| 4c0aB | 2.17 | 2.59 | 2.91 | 2.95 | 0.99 | 1.00 | 1.00 | 0.99 |
| 4c0sA | 3.59 | 4.18 | 3.97 | 4.08 | 0.66 | 0.41 | 0.71 | 0.61 |
| 4cczA | 3.15 | 3.71 | 3.50 | 3.71 | 0.64 | 0.76 | 0.73 | 0.61 |
| 4d0nB | 1.59 | 2.17 | 3.71 | 3.79 | 0.86 | 0.90 | 0.80 | 0.82 |
| 4d1iG | 3.11 | 3.62 | 3.68 | 3.96 | 0.58 | 0.49 | 0.47 | 0.48 |
| 4dimA | 1.61 | 2.66 | 2.56 | 3.00 | 1.00 | 1.00 | 0.93 | 0.98 |
| 4dj3A | 2.78 | 3.75 | 2.11 | 3.88 | 0.63 | 0.23 | 0.20 | 0.20 |
| 4dqaA | 3.16 | 3.40 | 5.04 | 3.53 | 0.16 | 0.08 | 0.08 | 0.08 |
| 4dt4A | 3.08 | 5.38 | 5.37 | 5.37 | 0.89 | 0.36 | 0.36 | 0.39 |
| 4dtfA | 4.16 | 4.97 | 5.09 | 5.20 | 0.84 | 0.78 | 0.55 | 0.65 |
| 4eo3A | 2.71 | 2.57 | 4.46 | 3.01 | 0.15 | 0.17 | 0.16 | 0.15 |
| 4eogA | 2.88 | 4.79 | 2.37 | 5.01 | 0.98 | 0.83 | 0.73 | 0.80 |
| 4etxA | 2.13 | 3.89 | 3.51 | 3.93 | 0.89 | 0.20 | 0.25 | 0.19 |
| 4ewtA | 2.61 | 3.40 | 2.93 | 3.12 | 0.66 | 0.61 | 0.70 | 0.63 |
| 4f23A | 3.64 | 3.11 | 4.25 | 4.27 | 0.50 | 0.69 | 0.12 | 0.15 |
| 4fe9A | 3.99 | 4.11 | 3.10 | 4.07 | 0.28 | 0.32 | 0.23 | 0.19 |
| 4fguA | 2.81 | 3.44 | 2.83 | 3.91 | 0.84 | 0.95 | 0.86 | 0.86 |
| 4fkcA | 1.36 | 2.47 | 2.65 | 2.74 | 0.91 | 0.79 | 0.82 | 0.78 |
| 4fxkC | 3.35 | 3.76 | 2.09 | 3.82 | 0.42 | 0.16 | 0.15 | 0.14 |
| 4fzbC | 2.25 | 2.56 | 2.76 | 2.68 | 0.88 | 0.88 | 0.88 | 0.87 |
| 4g1pA | 2.47 | 3.24 | 2.98 | 3.19 | 0.81 | 0.83 | 0.70 | 0.73 |
| 4gbyA | 1.82 | 2.50 | 2.62 | 2.57 | 1.00 | 0.98 | 0.97 | 0.99 |
| 4gfqA | 2.59 | 2.96 | 3.26 | 3.19 | 0.75 | 0.61 | 0.61 | 0.66 |
| 4ggmX | 3.24 | 3.53 | 2.57 | 3.49 | 0.51 | 0.34 | 0.35 | 0.34 |

| PDB ID | MAE | | | | Top-*L* | | | |
|---|---|---|---|---|---|---|---|---|
| | DeepIDDP | trRosettaX | trRosetta | SADA | DeepIDDP | trRosettaX | trRosetta | SADA |
| 4gslA | 2.77 | 3.50 | 4.00 | 3.43 | 0.17 | 0.18 | 0.17 | 0.17 |
| 4gyjA | 3.14 | 3.43 | 3.46 | 3.65 | 0.76 | 0.79 | 0.78 | 0.78 |
| 4h2aA | 3.93 | 4.23 | 2.65 | 4.22 | 0.63 | 0.48 | 0.41 | 0.42 |
| 4h3tA | 3.18 | 3.75 | 4.18 | 3.86 | 0.46 | 0.47 | 0.45 | 0.45 |
| 4hmoA | 3.01 | 3.17 | 1.71 | 3.78 | 0.84 | 0.86 | 0.81 | 0.81 |
| 4hvzA | 2.07 | 2.59 | 2.58 | 2.56 | 0.76 | 0.50 | 0.55 | 0.53 |
| 4i5sB | 1.87 | 2.73 | 1.62 | 2.85 | 0.70 | 0.68 | 0.70 | 0.64 |
| 4ie6A | 4.07 | 4.22 | 6.05 | 4.39 | 0.49 | 0.43 | 0.29 | 0.17 |
| 4iggB | 3.60 | 3.61 | 1.86 | 3.75 | 0.71 | 0.83 | 0.73 | 0.76 |
| 4il6B | 4.37 | 4.62 | 4.63 | 4.63 | 0.39 | 0.39 | 0.33 | 0.32 |
| 4indA | 3.87 | 4.12 | 4.12 | 4.13 | 0.45 | 0.21 | 0.14 | 0.18 |
| 4j9vA | 2.90 | 3.06 | 1.52 | 3.12 | 0.97 | 1.00 | 1.00 | 1.00 |
| 4jdzB | 3.67 | 3.76 | 3.76 | 3.77 | 0.38 | 0.25 | 0.17 | 0.21 |
| 4jxkA | 2.25 | 3.17 | 2.98 | 3.00 | 0.92 | 0.84 | 0.82 | 0.82 |
| 4k3bA | 2.84 | 3.42 | 1.82 | 2.86 | 0.52 | 0.52 | 0.61 | 0.58 |
| 4kc3B | 3.15 | 4.24 | 4.29 | 4.25 | 0.72 | 0.36 | 0.27 | 0.33 |
| 4kikB | 3.41 | 4.04 | 4.00 | 4.05 | 0.66 | 0.62 | 0.56 | 0.55 |
| 4kwuA | 4.16 | 4.11 | 2.91 | 4.14 | 0.11 | 0.24 | 0.15 | 0.15 |
| 4l5gA | 2.67 | 3.60 | 1.69 | 2.87 | 0.89 | 0.77 | 0.84 | 0.80 |
| 4lmfA | 2.46 | 4.47 | 4.35 | 4.41 | 0.85 | 0.70 | 0.65 | 0.63 |
| 4lpqA | 3.03 | 3.83 | 1.47 | 4.30 | 0.78 | 0.61 | 0.26 | 0.33 |
| 4lziA | 3.15 | 4.06 | 4.09 | 4.07 | 0.80 | 0.39 | 0.38 | 0.40 |
| 4m00A | 3.15 | 4.22 | 1.67 | 4.43 | 0.71 | 0.42 | 0.36 | 0.46 |
| 4m8mB | 3.96 | 4.05 | 4.06 | 4.06 | 0.39 | 0.26 | 0.14 | 0.22 |
| 4m8rA | 3.53 | 3.72 | 3.83 | 3.72 | 0.42 | 0.08 | 0.05 | 0.19 |
| 4m9pA | 3.43 | 3.99 | 4.00 | 4.02 | 0.70 | S0.48 | 0.51 | 0.46 |
| 4mzyA | 3.80 | 4.09 | 4.08 | 4.12 | 0.43 | 0.52 | 0.47 | 0.43 |
| 4n06B | 2.62 | 3.13 | 3.64 | 3.39 | 0.77 | 0.78 | 0.71 | 0.74 |
| 4nj5A | 3.53 | 4.04 | 2.12 | 4.12 | 0.76 | 0.56 | 0.42 | 0.45 |
| 4onyA | 3.84 | 4.19 | 4.06 | 4.19 | 0.29 | 0.32 | 0.33 | 0.31 |
| 4opaB | 2.00 | 4.02 | 3.01 | 4.12 | 0.74 | 0.37 | 0.26 | 0.23 |
| 4pt5A | 2.09 | 4.10 | 4.00 | 4.43 | 0.98 | 0.91 | 0.68 | 0.86 |
| 4pyhA | 4.28 | 4.73 | 4.70 | 4.68 | 0.51 | 0.20 | 0.19 | 0.20 |
| 4qkuB | 2.07 | 2.65 | 4.39 | 3.12 | 0.99 | 0.99 | 0.95 | 0.97 |
| 4rg1A | 3.86 | 4.22 | 4.22 | 4.23 | 0.78 | 0.74 | 0.69 | 0.71 |
| 4up9A | 3.29 | 3.85 | 3.55 | 4.23 | 0.32 | 0.30 | 0.29 | 0.33 |
| 4uwhA | 3.23 | 3.50 | 3.37 | 3.61 | 0.79 | 0.78 | 0.72 | 0.75 |
| 4w7sA | 3.11 | 3.43 | 4.77 | 3.43 | 0.15 | 0.17 | 0.15 | 0.14 |

**Table S4.** SADA-DeepIDDP and SADA compare detailed results on multi-domain benchmark set

| PDB ID | TM-score | | iRMSD | | PDB ID | TM-score | | iRMSD | |
|---|---|---|---|---|---|---|---|---|---|
| | SADA-DeepIDDP | SADA | SADA-DeepIDDP | SADA | | SADA-DeepIDDP | SADA | SADA-DeepIDDP | SADA |
| 1bf2A | 0.97 | 0.99 | 0.48 | 0.15 | 1pprM | 0.99 | 0.57 | 1.33 | 14.39 |
| 1bhgA | 0.99 | 1.00 | 4.37 | 4.25 | 1prrA | 0.64 | 0.60 | 4.48 | 4.07 |
| 1bp1A | 0.97 | 0.99 | 0.81 | 0.71 | 1q19A | 0.97 | 0.99 | 1.15 | 1.39 |
| 1c1zA | 0.57 | 0.46 | 1.96 | 2.53 | 1q25A | 0.92 | 0.49 | 2.36 | 1.15 |
| 1cjsA | 0.94 | 0.84 | 2.33 | 2.69 | 1qhdA | 0.92 | 0.90 | 1.59 | 2.14 |
| 1cjyA | 0.81 | 0.81 | 9.09 | 8.72 | 1qwrA | 0.78 | 1.00 | 1.06 | 0.05 |
| 1ck1A | 1.00 | 0.99 | 0.23 | 0.51 | 1qz9A | 0.99 | 0.99 | 0.76 | 0.59 |
| 1d2pA | 0.55 | 0.36 | 2.59 | 5.32 | 1r71B | 0.91 | 0.88 | 0.68 | 1.28 |
| 1ecrA | 0.88 | 0.88 | 2.76 | 2.66 | 1rh1A | 0.72 | 0.88 | 0.90 | 1.19 |
| 1efdN | 0.97 | 0.97 | 0.87 | 0.65 | 1rktA | 0.83 | 0.97 | 1.42 | 1.34 |
| 1f5qD | 0.99 | 0.99 | 0.72 | 0.36 | 1s6lA | 0.96 | 0.76 | 5.22 | 4.95 |
| 1f7uA | 0.97 | 0.97 | 1.16 | 0.41 | 1sb7B | 0.94 | 0.87 | 1.18 | 1.35 |
| 1fa9A | 1.00 | 1.00 | 0.25 | 0.02 | 1sp3A | 0.98 | 1.00 | 1.63 | 0.93 |
| 1fjrA | 0.89 | 0.86 | 0.88 | 0.89 | 1ug9A | 0.68 | 0.65 | 11.76 | 4.87 |
| 1fx7A | 0.97 | 0.87 | 1.01 | 0.50 | 1uzjA | 0.54 | 0.78 | 2.44 | 7.66 |
| 1g87B | 0.93 | 0.76 | 1.70 | 3.65 | 1vk1A | 0.79 | 0.85 | 5.40 | 3.14 |
| 1griA | 0.87 | 0.68 | 2.38 | 3.75 | 1vrmA | 0.99 | 0.99 | 0.72 | 0.17 |
| 1gu7A | 0.99 | 1.00 | 0.30 | 0.41 | 1vz6A | 0.99 | 0.95 | 5.12 | 4.43 |
| 1h88C | 0.44 | 0.71 | 3.73 | 4.80 | 1w3aA | 0.98 | 0.62 | 7.98 | 6.85 |
| 1hx6B | 0.71 | 0.98 | 2.26 | 0.19 | 1wv3A | 0.95 | 0.93 | 4.40 | 2.47 |
| 1itwA | 1.00 | 0.99 | 0.44 | 0.94 | 1x7pA | 0.99 | 0.99 | 5.30 | 0.66 |
| 1iwaA | 0.99 | 0.99 | 0.70 | 0.60 | 1x9yA | 0.98 | 0.63 | 1.88 | 10.89 |
| 1jkiA | 0.99 | 1.00 | 0.69 | 0.67 | 1xvuA | 0.95 | 0.98 | 1.24 | 0.76 |
| 1k7tA | 0.40 | 0.67 | 3.85 | 3.32 | 1y11A | 0.75 | 0.60 | 1.86 | 2.68 |
| 1kfqA | 0.91 | 0.97 | 1.29 | 1.14 | 1yiqA | 0.92 | 0.88 | 1.13 | 1.52 |
| 1ldjA | 0.66 | 0.62 | 2.77 | 2.78 | 1yy3A | 0.74 | 0.70 | 2.04 | 2.13 |
| 1m5qH | 0.58 | 0.58 | 3.26 | 4.03 | 1z1wA | 0.99 | 0.93 | 0.75 | 1.44 |
| 1m8pB | 0.40 | 0.68 | 6.42 | 2.78 | 1z87A | 0.68 | 0.67 | 3.09 | 3.59 |
| 1mkfA | 0.89 | 0.99 | 1.74 | 0.96 | 1zbuB | 0.90 | 0.79 | 2.52 | 4.32 |
| 1mkmB | 0.70 | 0.70 | 4.43 | 4.58 | 1ze1A | 0.99 | 0.99 | 0.75 | 0.78 |
| 1n80A | 0.98 | 1.00 | 0.12 | 0.14 | 1zpuA | 0.98 | 0.99 | 4.93 | 3.10 |
| 1nh2D | 0.56 | 0.58 | 3.58 | 3.74 | 1zy9A | 0.86 | 0.92 | 0.75 | 0.53 |
| 1ni5A | 0.65 | 0.75 | 6.85 | 4.17 | 2a1sC | 0.87 | 0.86 | 2.35 | 2.34 |
| 1nyqB | 0.85 | 0.87 | 1.47 | 0.97 | 2a3lA | 0.97 | 0.97 | 0.88 | 1.69 |
| 1nzjA | 1.00 | 0.99 | 0.48 | 0.70 | 2ablA | 0.71 | 0.70 | 2.81 | 3.45 |

| PDB ID | TM-score | | iRMSD | | PDB ID | TM-score | | iRMSD | |
|---|---|---|---|---|---|---|---|---|---|
| | SADA-DeepIDDP | SADA | SADA-DeepIDDP | SADA | | SADA-DeepIDDP | SADA | SADA-DeepIDDP | SADA |
| 2ahvA | 0.81 | 0.82 | 4.06 | 4.08 | 2l9yA | 0.92 | 0.69 | 1.53 | 2.48 |
| 2au3A | 0.84 | 0.81 | 7.14 | 4.86 | 2mbgA | 0.97 | 0.78 | 1.42 | 5.88 |
| 2b5uA | 0.67 | 0.48 | 2.54 | 1.06 | 2nsfA | 0.97 | 0.94 | 1.69 | 1.34 |
| 2bkpA | 0.66 | 0.83 | 3.87 | 1.06 | 2ntyB | 0.76 | 0.82 | 1.92 | 3.12 |
| 2bt1A | 0.99 | 0.99 | 0.94 | 0.86 | 2nykA | 0.98 | 0.95 | 1.05 | 1.31 |
| 2bydA | 0.80 | 0.98 | 2.36 | 0.50 | 2o6yA | 0.97 | 0.99 | 0.82 | 0.66 |
| 2c1yA | 0.85 | 0.56 | 1.70 | 3.38 | 2olsA | 0.69 | 0.49 | 3.67 | 4.21 |
| 2c43A | 0.71 | 0.93 | 2.63 | 0.89 | 2owbA | 0.98 | 0.98 | 0.80 | 1.06 |
| 2cxcA | 0.66 | 0.96 | 2.66 | 0.48 | 2piaA | 0.95 | 0.86 | 0.72 | 4.38 |
| 2d1cA | 0.89 | 0.84 | 4.39 | 3.87 | 2qfiA | 0.93 | 0.87 | 5.97 | 2.29 |
| 2d7iA | 0.74 | 0.74 | 2.62 | 3.01 | 2qp2A | 0.74 | 0.74 | 2.72 | 2.99 |
| 2dfyC | 0.68 | 0.66 | 2.01 | 2.98 | 2qygA | 0.98 | 0.97 | 0.98 | 0.74 |
| 2dlaA | 0.97 | 1.00 | 0.34 | 0.69 | 2r3vA | 0.99 | 1.00 | 0.65 | 0.59 |
| 2e9hA | 0.74 | 0.97 | 3.77 | 0.92 | 2r58A | 0.99 | 0.65 | 0.16 | 2.71 |
| 2e9xB | 0.97 | 0.99 | 1.59 | 1.09 | 2r5wB | 0.98 | 0.80 | 0.45 | 1.12 |
| 2evrA | 0.77 | 0.99 | 2.08 | 0.93 | 2r7dA | 0.97 | 0.98 | 1.17 | 1.81 |
| 2ew9A | 0.54 | 0.54 | 3.19 | 4.49 | 2ra1A | 0.50 | 0.49 | 3.14 | 8.12 |
| 2ewfA | 0.87 | 0.49 | 6.22 | 8.72 | 2uu7A | 0.97 | 0.99 | 0.60 | 1.15 |
| 2fd5A | 0.77 | 0.99 | 1.10 | 0.27 | 2uwnA | 0.57 | 0.77 | 1.12 | 1.43 |
| 2g3pA | 0.89 | 0.63 | 2.33 | 4.39 | 2v0nA | 0.64 | 0.45 | 3.37 | 0.99 |
| 2gg6A | 1.00 | 0.99 | 0.83 | 0.34 | 2v5dA | 0.86 | 0.69 | 3.58 | 5.11 |
| 2gh8A | 0.91 | 0.72 | 1.99 | 4.50 | 2vgmA | 0.92 | 0.76 | 3.82 | 13.90 |
| 2gsyE | 1.00 | 0.92 | 1.87 | 6.26 | 2w4bA | 0.86 | 0.83 | 4.78 | 7.31 |
| 2gt1A | 0.98 | 0.97 | 1.57 | 1.74 | 2w4mA | 0.86 | 0.89 | 3.02 | 2.84 |
| 2gzaC | 0.97 | 0.98 | 1.34 | 1.08 | 2wqrB | 0.47 | 0.49 | 2.41 | 3.63 |
| 2gzoA | 0.90 | 0.93 | 2.91 | 1.09 | 2x0cA | 0.68 | 0.66 | 1.91 | 2.84 |
| 2hjqA | 0.53 | 0.55 | 3.50 | 3.11 | 2x7iA | 0.96 | 0.98 | 0.64 | 0.99 |
| 2hwjA | 0.91 | 0.83 | 1.13 | 2.35 | 2x8kC | 0.90 | 0.98 | 1.67 | 0.46 |
| 2ii2A | 0.84 | 0.87 | 1.03 | 0.96 | 2xt6A | 0.61 | 0.54 | 4.21 | 5.28 |
| 2ijd1 | 0.77 | 0.77 | 1.78 | 2.31 | 2y25B | 0.73 | 0.57 | 2.31 | 9.22 |
| 2iu7A | 0.62 | 0.60 | 4.06 | 6.63 | 2y51A | 1.00 | 0.99 | 0.50 | 0.53 |
| 2iw2A | 0.90 | 0.98 | 1.23 | 0.90 | 2yb0E | 0.95 | 0.97 | 0.57 | 0.81 |
| 2j2cA | 0.99 | 0.96 | 6.00 | 6.52 | 2yilA | 0.64 | 0.63 | 2.80 | 2.81 |
| 2jz4A | 0.59 | 0.59 | 1.06 | 1.25 | 2yk0A | 0.92 | 0.72 | 1.87 | 2.96 |
| 2kdyA | 0.98 | 0.59 | 0.32 | 3.25 | 2yrqA | 0.53 | 0.52 | 2.07 | 1.54 |
| 2kfwA | 0.77 | 0.78 | 3.95 | 4.67 | 2z86C | 0.70 | 0.69 | 4.00 | 4.47 |
| 2kn4A | 0.62 | 0.65 | 1.58 | 1.61 | 2zpaB | 0.90 | 0.90 | 1.82 | 1.79 |

| PDB ID | TM-score | | iRMSD | | PDB ID | TM-score | | iRMSD | |
|---|---|---|---|---|---|---|---|---|---|
| | SADA-DeepIDDP | SADA | SADA-DeepIDDP | SADA | | SADA-DeepIDDP | SADA | SADA-DeepIDDP | SADA |
| 2zxcA | 0.96 | 0.82 | 0.53 | 3.53 | 3gmsA | 0.97 | 1.00 | 0.53 | 0.35 |
| 2zzqA | 0.71 | 0.46 | 3.97 | 20.67 | 3h2tA | 0.88 | 0.81 | 7.35 | 7.45 |
| 3a1iA | 0.90 | 0.93 | 0.95 | 1.24 | 3h5cB | 0.92 | 0.97 | 1.51 | 1.22 |
| 3a45A | 0.98 | 0.99 | 1.19 | 0.57 | 3hcsA | 0.72 | 0.80 | 4.75 | 1.21 |
| 3a56A | 0.67 | 0.66 | 2.27 | 4.65 | 3hjlA | 0.43 | 0.36 | 3.06 | 6.62 |
| 3afoA | 0.64 | 0.67 | 3.12 | 3.45 | 3hyiA | 0.69 | 0.71 | 3.24 | 3.78 |
| 3ajvA | 0.60 | 0.98 | 4.46 | 0.44 | 3hzzB | 1.00 | 1.00 | 0.51 | 0.42 |
| 3apoA | 0.52 | 0.45 | 4.35 | 3.12 | 3i2dA | 0.92 | 0.86 | 1.03 | 1.18 |
| 3aqkA | 0.95 | 0.96 | 0.95 | 0.52 | 3iam2 | 0.73 | 0.92 | 1.75 | 0.69 |
| 3arbA | 0.94 | 0.94 | 1.46 | 1.32 | 3ibjA | 0.58 | 0.58 | 2.73 | 2.71 |
| 3aujG | 0.71 | 0.99 | 3.77 | 0.48 | 3ifrA | 0.99 | 0.99 | 0.93 | 0.80 |
| 3b2zF | 0.96 | 0.76 | 1.83 | 6.73 | 3ippB | 0.96 | 0.74 | 0.90 | 4.87 |
| 3b43A | 0.36 | 0.36 | 5.29 | 5.61 | 3isqA | 1.00 | 1.00 | 0.50 | 0.29 |
| 3b7wA | 0.83 | 0.83 | 7.41 | 7.31 | 3j7aK | 0.95 | 0.96 | 0.65 | 0.79 |
| 3bt1U | 0.88 | 0.61 | 1.20 | 3.86 | 3jymB | 0.44 | 0.43 | 5.17 | 4.22 |
| 3bt3A | 0.64 | 0.95 | 3.66 | 0.87 | 3k1rA | 0.83 | 0.66 | 4.53 | 9.35 |
| 3bu2A | 0.91 | 0.91 | 0.84 | 0.54 | 3k2iA | 0.78 | 0.78 | 2.76 | 2.56 |
| 3c1yA | 0.90 | 0.63 | 1.61 | 3.70 | 3kbgA | 0.44 | 0.62 | 3.51 | 4.46 |
| 3c4tA | 0.89 | 0.74 | 2.98 | 4.90 | 3kh5A | 0.96 | 0.94 | 0.94 | 1.43 |
| 3craA | 0.94 | 0.61 | 1.56 | 3.27 | 3kjpA | 0.98 | 0.91 | 1.49 | 1.27 |
| 3cvzA | 0.85 | 0.87 | 3.89 | 4.12 | 3kq4B | 0.43 | 0.36 | 7.51 | 8.69 |
| 3cw2C | 0.71 | 0.70 | 1.70 | 1.46 | 3kt1A | 0.90 | 0.91 | 1.87 | 7.20 |
| 3d30A | 0.93 | 0.96 | 1.52 | 0.79 | 3ktmE | 0.87 | 0.63 | 1.86 | 2.12 |
| 3dupA | 0.90 | 0.98 | 1.57 | 0.50 | 3kwlA | 0.76 | 0.58 | 3.06 | 4.59 |
| 3eo5A | 0.54 | 0.64 | 5.84 | 5.16 | 3kzwA | 0.99 | 1.00 | 0.58 | 0.42 |
| 3errA | 0.86 | 0.79 | 1.28 | 1.83 | 3l76A | 0.80 | 0.89 | 3.46 | 2.65 |
| 3eswA | 0.97 | 0.89 | 0.46 | 0.82 | 3ld1A | 0.98 | 0.70 | 6.45 | 9.82 |
| 3eukH | 0.90 | 0.92 | 1.58 | 1.33 | 3lsgA | 0.84 | 0.86 | 1.12 | 0.88 |
| 3f83A | 0.39 | 0.46 | 3.22 | 4.01 | 3m1uA | 0.93 | 0.98 | 1.33 | 0.81 |
| 3fc3A | 0.45 | 0.54 | 7.86 | 9.36 | 3mc8A | 0.75 | 0.83 | 6.91 | 4.83 |
| 3fi7A | 0.88 | 0.99 | 0.63 | 0.72 | 3me4A | 0.94 | 0.90 | 0.63 | 0.34 |
| 3fvvA | 0.98 | 0.98 | 6.03 | 6.54 | 3ml4C | 0.95 | 0.65 | 1.45 | 5.55 |
| 3g79A | 0.89 | 0.90 | 1.24 | 1.01 | 3mw8A | 0.68 | 0.75 | 1.48 | 1.59 |
| 3gbgA | 0.98 | 0.64 | 1.74 | 5.13 | 3mwcA | 0.97 | 1.00 | 0.72 | 0.27 |
| 3gf5B | 0.44 | 0.41 | 5.15 | 6.59 | 3mx2B | 0.92 | 0.92 | 3.45 | 4.84 |

| PDB ID | TM-score | | iRMSD | | PDB ID | TM-score | | iRMSD | |
|---|---|---|---|---|---|---|---|---|---|
| | SADA-DeepIDDP | SADA | SADA-DeepIDDP | SADA | | SADA-DeepIDDP | SADA | SADA-DeepIDDP | SADA |
| 3mzfA | 0.87 | 0.93 | 2.04 | 1.33 | 3soaA | 0.97 | 0.73 | 1.55 | 1.89 |
| 3njaB | 0.61 | 0.61 | 2.10 | 1.65 | 3spgA | 0.99 | 0.98 | 1.31 | 0.46 |
| 3npfA | 0.82 | 0.93 | 2.31 | 1.07 | 3swjA | 0.93 | 0.71 | 1.59 | 8.70 |
| 3nqiA | 0.88 | 0.94 | 2.95 | 0.83 | 3t58B | 0.93 | 0.58 | 6.41 | 14.24 |
| 3nsjA | 0.84 | 0.82 | 2.81 | 2.82 | 3t7jA | 0.91 | 0.97 | 6.05 | 0.75 |
| 3nt8A | 0.73 | 0.76 | 5.04 | 3.01 | 3tixD | 0.66 | 0.63 | 4.52 | 1.46 |
| 3ntkA | 0.97 | 0.98 | 1.52 | 1.25 | 3tp9A | 0.93 | 0.87 | 1.75 | 2.39 |
| 3oaaG | 0.99 | 0.98 | 0.86 | 1.47 | 3u07C | 0.89 | 0.88 | 1.30 | 1.07 |
| 3ob8A | 0.91 | 0.97 | 1.47 | 2.06 | 3u0kA | 0.70 | 0.61 | 1.56 | 2.28 |
| 3og5A | 0.89 | 0.97 | 1.23 | 0.77 | 3u0oB | 0.99 | 0.99 | 2.08 | 0.83 |
| 3oh0A | 0.92 | 0.97 | 2.15 | 3.94 | 3u9gA | 0.98 | 0.77 | 1.12 | 3.95 |
| 3opfB | 0.92 | 0.68 | 2.08 | 6.15 | 3ua3A | 0.61 | 0.58 | 16.41 | 8.74 |
| 3orjA | 0.91 | 0.66 | 2.56 | 1.52 | 3ub1D | 0.59 | 0.62 | 5.71 | 4.78 |
| 3p53A | 0.52 | 0.37 | 5.03 | 8.80 | 3ubhA | 0.74 | 0.68 | 6.52 | 3.18 |
| 3pcsB | 0.96 | 0.86 | 2.50 | 3.37 | 3uitD | 0.86 | 0.57 | 1.08 | 2.59 |
| 3plaA | 0.66 | 0.65 | 2.86 | 6.70 | 3uj0A | 0.90 | 0.94 | 0.93 | 1.19 |
| 3po3S | 0.57 | 0.96 | 2.62 | 1.01 | 3uo3A | 0.87 | 0.99 | 2.69 | 0.31 |
| 3ptyA | 0.98 | 0.75 | 0.53 | 1.64 | 3v7oB | 0.65 | 0.65 | 2.30 | 1.94 |
| 3pvlA | 0.89 | 0.68 | 1.75 | 5.38 | 3vlaA | 1.00 | 1.00 | 0.00 | 0.00 |
| 3pxpA | 0.85 | 0.74 | 2.67 | 3.50 | 3vn4A | 0.92 | 0.47 | 8.06 | 10.15 |
| 3qavA | 0.88 | 0.98 | 1.46 | 1.21 | 3vr8B | 1.00 | 1.00 | 0.24 | 0.32 |
| 3qe9Y | 0.95 | 0.94 | 1.02 | 2.50 | 3vsmA | 0.82 | 0.80 | 1.56 | 1.01 |
| 3qf4B | 0.93 | 0.93 | 1.14 | 1.13 | 3vstA | 0.65 | 1.00 | 1.70 | 0.35 |
| 3qjjA | 0.97 | 0.97 | 0.81 | 1.01 | 3w1bA | 0.71 | 0.72 | 2.44 | 3.30 |
| 3qjoA | 0.92 | 0.80 | 1.75 | 1.06 | 3w2wA | 0.86 | 0.85 | 1.63 | 2.18 |
| 3qphA | 0.71 | 0.48 | 1.37 | 3.22 | 3wkuA | 0.94 | 0.76 | 0.88 | 5.66 |
| 3qtdA | 1.00 | 1.00 | 0.47 | 0.63 | 3zh9B | 0.71 | 0.62 | 3.86 | 4.05 |
| 3qyeA | 0.97 | 0.99 | 0.51 | 3.29 | 3zniA | 0.81 | 0.41 | 1.70 | 14.91 |
| 3r05A | 0.32 | 0.32 | 6.60 | 6.66 | 3zvmA | 0.78 | 0.59 | 2.54 | 6.63 |
| 3r6bA | 0.55 | 0.54 | 4.12 | 3.27 | 4acoA | 0.89 | 0.91 | 16.25 | 4.86 |
| 3rfyA | 0.98 | 0.95 | 0.68 | 0.92 | 4aimA | 0.84 | 0.91 | 2.16 | 1.20 |
| 3rh7A | 0.97 | 0.71 | 2.14 | 5.84 | 4ak1A | 0.46 | 0.29 | 5.07 | 9.43 |
| 3rimA | 0.90 | 0.98 | 0.54 | 0.07 | 4alzA | 0.37 | 0.69 | 7.21 | 7.80 |
| 3rrpA | 0.75 | 0.98 | 1.39 | 0.40 | 4ap5A | 0.98 | 0.99 | 5.29 | 0.67 |
| 3rwxA | 0.57 | 0.60 | 3.69 | 3.29 | 4aq1A | 0.50 | 0.45 | 7.69 | 8.26 |
| 3sb4A | 0.83 | 0.61 | 2.08 | 7.42 | 4aqfB | 0.96 | 0.88 | 1.26 | 3.71 |
| 3seoB | 0.80 | 0.77 | 1.81 | 1.07 | 4ax8A | 0.61 | 0.64 | 3.56 | 3.86 |

| PDB ID | TM-score | | iRMSD | | PDB ID | TM-score | | iRMSD | |
|---|---|---|---|---|---|---|---|---|---|
| | SADA-DeepIDDP | SADA | SADA-DeepIDDP | SADA | | SADA-DeepIDDP | SADA | SADA-DeepIDDP | SADA |
| 4axdA | 0.94 | 0.99 | 1.10 | 0.44 | 4ie6A | 0.69 | 0.70 | 2.76 | 2.12 |
| 4b21A | 0.95 | 0.99 | 1.04 | 0.29 | 4iggB | 0.46 | 0.31 | 5.38 | 4.89 |
| 4b3iA | 0.94 | 0.94 | 8.07 | 3.99 | 4il6B | 0.82 | 0.73 | 3.85 | 7.27 |
| 4bd9B | 0.56 | 0.61 | 5.22 | 5.51 | 4indA | 0.71 | 0.46 | 5.21 | 13.16 |
| 4bfiB | 0.82 | 0.83 | 1.27 | 0.94 | 4j9vA | 0.96 | 0.87 | 2.12 | 2.38 |
| 4bt9B | 0.60 | 0.61 | 2.16 | 1.43 | 4jdzB | 0.59 | 0.70 | 3.58 | 2.18 |
| 4c0aB | 0.88 | 0.57 | 1.62 | 0.55 | 4jxkA | 0.98 | 0.98 | 0.70 | 0.46 |
| 4c0sA | 0.64 | 0.63 | 4.31 | 5.62 | 4k3bA | 0.64 | 0.56 | 3.73 | 3.19 |
| 4cczA | 0.59 | 0.96 | 1.13 | 2.02 | 4kc3B | 0.77 | 0.76 | 1.98 | 1.78 |
| 4d0nB | 0.93 | 0.91 | 1.58 | 0.89 | 4kikB | 0.88 | 0.73 | 1.37 | 3.28 |
| 4d1iG | 0.94 | 0.96 | 2.69 | 2.38 | 4kwuA | 0.57 | 0.78 | 3.41 | 3.26 |
| 4dimA | 0.91 | 0.89 | 0.91 | 4.77 | 4l5gA | 0.75 | 0.97 | 5.86 | 1.47 |
| 4dj3A | 0.92 | 0.98 | 0.90 | 1.05 | 4lmfA | 0.58 | 0.49 | 4.97 | 5.53 |
| 4dqaA | 0.64 | 0.64 | 0.49 | 0.12 | 4lpqA | 0.85 | 0.91 | 5.43 | 0.97 |
| 4dt4A | 0.91 | 0.98 | 2.11 | 1.19 | 4lziA | 0.94 | 0.67 | 3.34 | 10.09 |
| 4dtfA | 1.00 | 0.95 | 0.39 | 0.28 | 4m00A | 0.51 | 0.52 | 3.22 | 6.89 |
| 4eo3A | 0.57 | 0.58 | 0.95 | 1.53 | 4m8mB | 0.87 | 0.87 | 3.68 | 9.61 |
| 4eogA | 0.99 | 0.98 | 1.82 | 1.71 | 4m8rA | 0.84 | 0.77 | 9.19 | 14.43 |
| 4etxA | 0.99 | 0.55 | 6.05 | 9.47 | 4m9pA | 0.59 | 0.51 | 5.20 | 7.64 |
| 4ewtA | 0.87 | 0.84 | 1.11 | 1.44 | 4mzyA | 1.00 | 1.00 | 0.20 | 0.19 |
| 4f23A | 0.98 | 1.00 | 0.33 | 0.62 | 4n06B | 0.99 | 0.99 | 0.70 | 1.34 |
| 4fe9A | 0.48 | 0.47 | 3.31 | 3.94 | 4nj5A | 0.99 | 0.69 | 2.39 | 19.53 |
| 4fguA | 0.93 | 0.92 | 2.73 | 1.81 | 4onyA | 0.87 | 0.81 | 1.59 | 1.85 |
| 4fkcA | 0.94 | 0.94 | 1.09 | 0.97 | 4opaB | 0.78 | 0.71 | 4.70 | 9.16 |
| 4fxkC | 0.98 | 0.97 | 1.83 | 2.43 | 4pt5A | 0.90 | 0.91 | 1.15 | 1.93 |
| 4fzbC | 0.95 | 0.99 | 1.80 | 0.75 | 4pyhA | 0.86 | 0.77 | 7.77 | 8.61 |
| 4g1pA | 1.00 | 0.93 | 0.67 | 1.21 | 4qkuB | 0.97 | 1.00 | 5.13 | 0.58 |
| 4gbyA | 0.98 | 1.00 | 1.09 | 0.72 | 4rg1A | 0.84 | 0.77 | 4.22 | 4.41 |
| 4gfqA | 0.69 | 0.76 | 3.44 | 3.60 | 4up9A | 0.75 | 0.96 | 7.50 | 9.11 |
| 4ggmX | 0.86 | 0.61 | 7.04 | 8.12 | 4uwhA | 0.99 | 0.99 | 1.39 | 0.88 |
| 4gslA | 0.61 | 0.55 | 4.48 | 4.72 | 4w7sA | 0.64 | 0.64 | 5.10 | 4.73 |
| 4gyjA | 1.00 | 1.00 | 0.35 | 0.23 | | | | | |
| 4h2aA | 0.82 | 0.61 | 2.02 | 4.72 | | | | | |
| 4h3tA | 0.94 | 0.98 | 1.12 | 0.73 | | | | | |
| 4hmoA | 0.98 | 0.99 | 6.14 | 0.32 | | | | | |
| 4hvzA | 0.81 | 0.75 | 1.79 | 2.59 | | | | | |
| 4i5sB | 0.59 | 0.55 | 4.73 | 4.60 | | | | | |

**Table S5.** Detailed results of a full-chain modeling comparison between SADA-DeepidDP, SADA, and AlphaFold2 for 68 human multidomain proteins

| PDB ID | TM-score | | | iRMSD | | |
|---|---|---|---|---|---|---|
| | SADA-DeepIDDP | SADA | AlphaFold2 | SADA-DeepIDDP | SADA | AlphaFold2 |
| 1b3uA | 0.82 | 0.83 | 0.72 | 0.83 | 1.15 | 1.94 |
| 1dt9A | 0.95 | 0.59 | 0.81 | 1.00 | 2.35 | 8.87 |
| 1ggzA | 0.60 | 0.60 | 0.59 | 0.61 | 0.61 | 0.61 |
| 1griA | 0.60 | 0.52 | 0.46 | 4.32 | 5.40 | 4.16 |
| 1nmvA | 0.65 | 0.66 | 0.66 | 1.76 | 1.43 | 1.42 |
| 1oqyA | 0.28 | 0.29 | 0.22 | 12.37 | 8.02 | 8.70 |
| 1q8kA | 0.86 | 0.63 | 0.60 | 2.92 | 3.94 | 5.35 |
| 1qbkB | 0.78 | 0.80 | 0.79 | 3.24 | 3.18 | 4.09 |
| 1s8oA | 1.00 | 1.00 | 0.72 | 0.04 | 0.30 | 2.30 |
| 1st0A | 0.98 | 0.99 | 0.78 | 0.05 | 1.24 | 1.78 |
| 1ytqA | 0.96 | 0.96 | 0.49 | 0.58 | 0.58 | 5.63 |
| 1zzaA | 0.38 | 0.41 | 0.34 | 6.73 | 5.48 | 6.63 |
| 2ar7A | 0.83 | 0.82 | 0.75 | 7.11 | 7.47 | 7.41 |
| 2dybA | 0.73 | 0.68 | 0.71 | 5.65 | 9.36 | 6.29 |
| 2eyzA | 0.38 | 0.33 | 0.32 | 10.03 | 12.20 | 10.12 |
| 2gf5A | 0.69 | 0.68 | 0.48 | 5.20 | 5.53 | 7.56 |
| 2hxyA | 0.53 | 0.53 | 0.53 | 6.88 | 6.70 | 6.81 |
| 2kdoA | 0.61 | 0.55 | 0.43 | 3.38 | 5.80 | 6.66 |
| 2kn6A | 0.45 | 0.45 | 0.45 | 11.74 | 11.55 | 12.31 |
| 2kr0A | 0.31 | 0.31 | 0.31 | 21.10 | 25.66 | 26.77 |
| 2l4hA | 0.67 | 0.70 | 0.75 | 7.82 | 7.92 | 7.60 |
| 2l6lA | 0.36 | 0.39 | 0.38 | 6.60 | 6.81 | 6.10 |
| 2looA | 0.36 | 0.36 | 0.36 | 5.43 | 5.26 | 5.39 |
| 2mphA | 0.49 | 0.49 | 0.50 | 18.30 | 19.98 | 20.18 |
| 2mzhA | 0.75 | 0.74 | 0.74 | 2.25 | 2.24 | 2.36 |

| PDB ID | TM-score | | | iRMSD | | |
|---|---|---|---|---|---|---|
| | SADA-DeepIDDP | SADA | AlphaFold2 | SADA-DeepIDDP | SADA | AlphaFold2 |
| 2nn6I | 0.35 | 0.30 | 0.28 | 2.63 | 1.74 | 2.31 |
| 2p01A | 0.77 | 0.77 | 0.77 | 10.88 | 11.90 | 13.51 |
| 2wzbA | 0.95 | 0.96 | 0.67 | 5.32 | 5.29 | 5.34 |
| 2ymbC | 0.75 | 0.78 | 0.69 | 1.94 | 1.84 | 1.78 |
| 3ajmA | 0.63 | 0.66 | 0.62 | 1.27 | 1.27 | 1.27 |
| 3j0aA | 0.61 | 0.56 | 0.61 | 6.02 | 6.09 | 6.01 |
| 3t5oA | 0.72 | 0.70 | 0.55 | 2.03 | 5.71 | 2.08 |
| 3vklA | 0.73 | 0.70 | 0.53 | 2.21 | 1.73 | 2.47 |
| 4bitA | 0.95 | 0.97 | 0.75 | 5.80 | 5.50 | 5.73 |
| 4cooA | 0.37 | 0.38 | 0.42 | 2.26 | 2.26 | 10.26 |
| 4d1eA | 0.59 | 0.46 | 0.50 | 4.62 | 4.21 | 4.44 |
| 4kfzA | 0.87 | 0.92 | 0.67 | 2.83 | 3.17 | 2.89 |
| 4mspA | 0.89 | 0.89 | 0.78 | 0.64 | 0.64 | 1.64 |
| 5c19A | 0.82 | 0.81 | 0.73 | 1.92 | 1.77 | 2.02 |
| 5edmA | 0.40 | 0.39 | 0.40 | 3.09 | 1.85 | 6.64 |
| 5ivw2 | 0.43 | 0.43 | 0.43 | 11.63 | 11.63 | 11.63 |
| 5t7cA | 0.67 | 0.58 | 0.58 | 8.06 | 8.08 | 7.87 |
| 5u6gA | 0.69 | 0.68 | 0.70 | 1.48 | 1.54 | 2.37 |
| 5xwmA | 0.71 | 0.65 | 0.65 | 3.13 | 3.24 | 3.26 |
| 5ztfA | 0.63 | 0.62 | 0.62 | 5.28 | 6.44 | 6.61 |
| 6fonA | 0.64 | 0.63 | 0.67 | 8.23 | 8.22 | 8.03 |
| 6hhjA | 0.58 | 0.57 | 0.45 | 5.16 | 7.03 | 7.20 |
| 6jt0B | 0.42 | 0.39 | 0.51 | 5.63 | 6.57 | 13.54 |
| 6kvgA | 0.70 | 0.69 | 0.70 | 3.79 | 4.26 | 4.08 |
| 6nr8A | 0.63 | 0.68 | 0.73 | 5.10 | 6.44 | 4.94 |
| 6r6hB | 0.70 | 0.70 | 0.70 | 1.55 | 1.97 | 2.82 |
| 6s7pA | 0.93 | 0.94 | 0.73 | 6.80 | 7.60 | 9.28 |

| PDB ID | TM-score | | | iRMSD | | |
|---|---|---|---|---|---|---|
| | SADA-DeepIDDP | SADA | AlphaFold2 | SADA-DeepIDDP | SADA | AlphaFold2 |
| 6sn1B | 0.77 | 0.81 | 0.70 | 0.67 | 0.60 | 2.97 |
| 6tgbB | 0.80 | 0.79 | 0.80 | 2.93 | 2.41 | 3.04 |
| 6ulgF | 0.78 | 0.77 | 0.70 | 3.21 | 3.40 | 3.48 |
| 6wm2P | 0.90 | 0.75 | 0.75 | 2.71 | 2.39 | 3.33 |
| 6y4lA | 0.77 | 0.71 | 0.77 | 2.36 | 3.45 | 3.55 |
| 7a09J | 0.91 | 0.79 | 0.76 | 2.06 | 2.65 | 2.68 |
| 7a5oA | 0.79 | 0.81 | 0.60 | 2.66 | 2.93 | 2.82 |
| 7cccB | 0.43 | 0.44 | 0.43 | 0.51 | 0.86 | 0.57 |
| 7dmeA | 0.60 | 0.60 | 0.60 | 2.94 | 2.94 | 2.94 |
| 7emf0 | 0.83 | 0.85 | 0.64 | 5.18 | 5.18 | 5.18 |
| 7kzpF | 0.88 | 0.88 | 0.59 | 4.04 | 4.66 | 8.24 |
| 7kzpL | 0.85 | 0.85 | 0.57 | 1.30 | 1.99 | 2.08 |
| 7kzpQ | 0.46 | 0.44 | 0.64 | 1.57 | 2.67 | 3.67 |
| 7lbmv | 0.80 | 0.76 | 0.75 | 2.25 | 2.33 | 0.80 |
| 7lbmy | 0.73 | 0.75 | 0.70 | 1.04 | 1.04 | 1.04 |
| 7lsyY | | | | 9.11 | 7.74 | 7.28 |

**Table S6.** Detailed results in 17 CASP14 multi-domain proteins

| PDB ID | TM-score | | | | |
|---|---|---|---|---|---|
| | SADA-DeepIDDP | SADA | AlphaFold2-CASP14 | AlphaFold2-local | SADA-DeepIDDP-native |
| T1024 | 0.89 | 0.98 | 0.67 | 0.85 | 0.97 |
| T1030 | 0.72 | 0.72 | 0.73 | 0.74 | 0.72 |
| T1038 | 0.98 | 0.90 | 0.92 | 0.92 | 1.00 |
| T1047s2 | 0.92 | 0.93 | 0.79 | 0.87 | 1.00 |
| T1050 | 0.96 | 0.97 | 0.98 | 0.97 | 0.97 |
| T1052 | 0.69 | 0.70 | 0.7 | 0.70 | 0.69 |
| T1053 | 0.99 | 0.97 | 0.98 | 0.98 | 1.00 |
| T1058 | 0.89 | 0.74 | 0.96 | 0.96 | 0.94 |
| T1061 | 0.71 | 0.59 | 0.75 | 0.79 | 0.86 |
| T1070 | 0.47 | 0.47 | 0.49 | 0.47 | 0.47 |
| T1091 | 0.90 | 0.88 | 0.83 | 0.89 | 0.91 |
| T1092 | 0.95 | 0.94 | 0.92 | 0.94 | 0.92 |
| T1093 | 0.91 | 0.91 | 0.94 | 0.94 | 0.98 |
| T1094 | 0.93 | 0.91 | 0.91 | 0.92 | 0.98 |
| T1096 | 0.59 | 0.55 | 0.55 | 0.55 | 0.58 |
| T1100 | 0.90 | 0.90 | 0.93 | 0.93 | 0.97 |
| T1101 | 0.94 | 0.94 | 0.93 | 0.94 | 0.99 |
| Average | 0.84 | 0.82 | 0.82 | 0.84 | 0.88 |

**Table S7.** Detailed results of contribution of inter-domain features and data enhancement

| PDB ID | MAE | | | | Top-*L* | | | |
|---|---|---|---|---|---|---|---|---|
| | DeepIDDP | DeepIDDP[1] | DeepIDDP[2] | DeepIDDP[3] | DeepIDDP | DeepIDDP[1] | DeepIDDP[2] | DeepIDDP[3] |
| 1bf2A | 2.25 | 3.03 | 4.32 | 4.21 | 0.99 | 0.98 | 0.15 | 0.71 |
| 1bhgA | 2.76 | 2.56 | 2.99 | 4.33 | 0.98 | 0.98 | 0.95 | 0.14 |
| 1bp1A | 2.91 | 2.97 | 4.10 | 4.46 | 0.80 | 0.80 | 0.62 | 0.37 |
| 1c1zA | 1.96 | 2.24 | 3.25 | 3.81 | 0.88 | 0.80 | 0.78 | 0.75 |
| 1cjsA | 1.63 | 2.84 | 3.12 | 3.71 | 0.90 | 0.58 | 0.54 | 0.55 |
| 1cjyA | 4.02 | 4.14 | 4.33 | 4.35 | 0.14 | 0.11 | 0.16 | 0.46 |
| 1ck1A | 1.97 | 1.98 | 2.67 | 4.12 | 1.00 | 0.98 | 0.99 | 0.73 |
| 1d2pA | 3.66 | 3.52 | 4.16 | 4.43 | 0.76 | 0.85 | 0.70 | 0.50 |
| 1ecrA | 1.75 | 4.98 | 4.95 | 4.95 | 0.84 | 0.00 | 0.00 | 0.51 |
| 1efdN | 2.02 | 1.63 | 1.88 | 2.22 | 1.00 | 1.00 | 1.00 | 0.59 |
| 1f5qD | 2.04 | 1.82 | 2.28 | 3.01 | 1.00 | 1.00 | 1.00 | 0.22 |
| 1f7uA | 1.89 | 1.86 | 2.26 | 4.39 | 1.00 | 1.00 | 1.00 | 0.27 |
| 1fa9A | 2.68 | 3.23 | 3.66 | 4.00 | 0.83 | 0.68 | 0.61 | 0.01 |
| 1fjrA | 1.63 | 1.63 | 2.01 | 4.28 | 0.98 | 0.99 | 0.99 | 0.10 |
| 1fx7A | 2.09 | 2.16 | 2.27 | 3.22 | 1.00 | 0.99 | 0.98 | 0.95 |
| 1g87B | 3.61 | 3.43 | 4.30 | 4.45 | 0.48 | 0.51 | 0.45 | 0.51 |
| 1griA | 2.61 | 3.13 | 3.79 | 4.24 | 0.82 | 0.80 | 0.61 | 0.14 |
| 1gu7A | 1.58 | 1.48 | 1.52 | 3.61 | 0.93 | 0.96 | 0.97 | 0.20 |
| 1h88C | 1.23 | 1.92 | 2.09 | 3.86 | 0.99 | 0.96 | 0.89 | 0.62 |
| 1hx6B | 4.44 | 4.33 | 4.45 | 4.46 | 0.06 | 0.22 | 0.24 | 0.08 |
| 1itwA | 3.01 | 3.73 | 3.86 | 3.86 | 0.64 | 0.58 | 0.04 | 0.21 |
| 1iwaA | 2.89 | 3.46 | 3.87 | 4.37 | 0.69 | 0.59 | 0.62 | 0.14 |
| 1jkiA | 2.86 | 2.71 | 3.68 | 4.03 | 0.83 | 0.87 | 0.68 | 0.28 |
| 1k7tA | 1.65 | 2.21 | 2.20 | 3.99 | 0.98 | 0.86 | 0.90 | 0.41 |
| 1kfqA | 2.26 | 2.28 | 2.68 | 4.25 | 1.00 | 1.00 | 0.98 | 0.46 |
| 1ldjA | 2.85 | 2.95 | 2.81 | 4.50 | 0.96 | 0.95 | 0.95 | 0.06 |
| 1m5qH | 2.56 | 3.03 | 3.35 | 4.08 | 0.59 | 0.38 | 0.36 | 0.94 |
| 1m8pB | 2.04 | 2.82 | 3.86 | 4.38 | 0.99 | 0.94 | 0.85 | 0.34 |
| 1mkfA | 3.63 | 3.69 | 4.26 | 4.28 | 0.62 | 0.49 | 0.21 | 1.00 |
| 1mkmB | 1.65 | 2.43 | 3.31 | 4.28 | 0.69 | 0.52 | 0.45 | 0.91 |
| 1n80A | 3.41 | 3.54 | 4.06 | 4.37 | 0.63 | 0.60 | 0.36 | 0.24 |
| 1nh2D | 1.74 | 1.57 | 1.47 | 3.56 | 0.83 | 0.79 | 0.84 | 0.11 |
| 1ni5A | 2.29 | 3.66 | 3.89 | 4.39 | 0.91 | 0.58 | 0.56 | 0.64 |
| 1nyqB | 2.27 | 2.00 | 2.83 | 4.07 | 0.80 | 0.80 | 0.71 | 0.45 |
| 1nzjA | 1.37 | 1.39 | 1.66 | 4.28 | 1.00 | 1.00 | 1.00 | 0.88 |
| 1pprM | 2.51 | 2.04 | 4.20 | 4.37 | 0.92 | 0.93 | 0.44 | 0.07 |
| 1prrA | 2.13 | 1.85 | 2.39 | 4.22 | 0.91 | 0.96 | 0.83 | 0.13 |
| 1q19A | 3.29 | 3.30 | 3.40 | 3.75 | 0.71 | 0.72 | 0.64 | 0.05 |
| 1q25A | 3.37 | 3.61 | 4.00 | 4.06 | 0.69 | 0.60 | 0.41 | 0.42 |
| 1qhdA | 3.70 | 4.54 | 4.89 | 4.89 | 0.70 | 0.32 | 0.09 | 0.62 |
| 1qwrA | 4.30 | 4.16 | 4.29 | 4.49 | 0.26 | 0.26 | 0.25 | 0.16 |
| 1qz9A | 1.72 | 1.69 | 2.00 | 3.11 | 0.99 | 0.99 | 0.98 | 0.88 |

| PDB ID | MAE | | | | Top-*L* | | | |
|---|---|---|---|---|---|---|---|---|
| | DeepIDDP | DeepIDDP[1] | DeepIDDP[2] | DeepIDDP[3] | DeepIDDP | DeepIDDP[1] | DeepIDDP[2] | DeepIDDP[3] |
| 1r71B | 0.86 | 1.54 | 1.45 | 2.76 | 1.00 | 1.00 | 1.00 | 0.52 |
| 1rh1A | 3.82 | 4.87 | 4.96 | 4.96 | 0.71 | 0.04 | 0.00 | 0.96 |
| 1rktA | 1.48 | 2.03 | 2.21 | 3.25 | 0.99 | 1.00 | 0.93 | 0.25 |
| 1s6lA | 2.81 | 2.29 | 2.66 | 3.43 | 0.81 | 0.78 | 0.67 | 0.40 |
| 1sb7B | 3.61 | 3.61 | 3.75 | 4.31 | 0.51 | 0.52 | 0.54 | 0.41 |
| 1sp3A | 3.61 | 3.70 | 4.16 | 4.63 | 0.92 | 0.91 | 0.87 | 0.72 |
| 1ug9A | 3.51 | 3.84 | 3.88 | 4.24 | 0.70 | 0.80 | 0.75 | 0.66 |
| 1uzjA | 2.44 | 3.84 | 3.81 | 4.43 | 0.98 | 0.81 | 0.83 | 0.98 |
| 1vk1A | 1.93 | 2.04 | 2.66 | 4.07 | 0.93 | 0.91 | 0.80 | 0.80 |
| 1vrmA | 1.53 | 1.28 | 1.47 | 3.04 | 1.00 | 1.00 | 1.00 | 0.58 |
| 1vz6A | 1.75 | 1.62 | 1.98 | 2.94 | 0.87 | 0.89 | 0.89 | 0.18 |
| 1w3aA | 2.87 | 2.74 | 3.81 | 4.07 | 0.63 | 0.62 | 0.51 | 0.10 |
| 1wv3A | 2.15 | 2.40 | 3.06 | 4.13 | 0.92 | 0.81 | 0.59 | 0.01 |
| 1x7pA | 1.56 | 1.15 | 1.45 | 1.82 | 0.95 | 1.00 | 0.99 | 0.20 |
| 1x9yA | 3.58 | 3.64 | 3.94 | 4.37 | 0.76 | 0.72 | 0.74 | 0.73 |
| 1xvuA | 2.63 | 2.34 | 2.61 | 3.88 | 0.30 | 0.41 | 0.42 | 0.38 |
| 1y11A | 3.06 | 3.16 | 3.29 | 3.43 | 0.20 | 0.10 | 0.09 | 0.54 |
| 1yiqA | 3.87 | 3.81 | 4.13 | 4.23 | 0.59 | 0.69 | 0.58 | 0.12 |
| 1yy3A | 3.68 | 3.01 | 3.21 | 4.27 | 0.38 | 0.57 | 0.44 | 0.64 |
| 1z1wA | 2.54 | 2.79 | 2.51 | 4.17 | 0.94 | 0.97 | 0.94 | 0.61 |
| 1z87A | 2.96 | 3.18 | 3.37 | 3.69 | 0.25 | 0.15 | 0.10 | 0.06 |
| 1zbuB | 3.09 | 3.99 | 4.10 | 4.10 | 0.44 | 0.23 | 0.17 | 0.02 |
| 1ze1A | 1.71 | 3.01 | 4.04 | 3.12 | 0.79 | 0.70 | 0.11 | 0.73 |
| 1zpuA | 2.30 | 3.44 | 4.27 | 4.15 | 0.99 | 0.97 | 0.42 | 0.65 |
| 1zy9A | 3.77 | 4.02 | 4.15 | 4.31 | 0.59 | 0.55 | 0.55 | 0.34 |
| 2a1sC | 2.23 | 2.43 | 3.40 | 4.70 | 0.32 | 0.27 | 0.20 | 0.01 |
| 2a3lA | 2.34 | 2.47 | 3.05 | 4.15 | 0.85 | 0.80 | 0.68 | 0.19 |
| 2ablA | 2.00 | 1.97 | 1.94 | 3.81 | 0.81 | 0.88 | 0.90 | 0.03 |
| 2ahvA | 3.02 | 2.45 | 3.01 | 4.04 | 0.77 | 0.83 | 0.87 | 0.19 |
| 2au3A | 2.25 | 2.40 | 2.65 | 3.26 | 0.90 | 0.88 | 0.94 | 0.91 |
| 2b5uA | 2.95 | 4.12 | 4.19 | 4.29 | 0.60 | 0.11 | 0.08 | 0.77 |
| 2bkpA | 2.82 | 2.66 | 3.15 | 3.62 | 0.66 | 0.80 | 0.73 | 0.01 |
| 2bt1A | 3.52 | 4.17 | 4.43 | 4.41 | 0.79 | 0.84 | 0.30 | 0.21 |
| 2bydA | 3.79 | 3.61 | 3.98 | 4.67 | 0.63 | 0.70 | 0.66 | 0.91 |
| 2c1yA | 2.20 | 2.63 | 3.27 | 3.75 | 0.59 | 0.44 | 0.33 | 0.45 |
| 2c43A | 2.40 | 2.10 | 3.21 | 4.16 | 0.85 | 0.93 | 0.89 | 0.02 |
| 2cxcA | 1.37 | 1.31 | 1.49 | 2.47 | 1.00 | 1.00 | 1.00 | 0.32 |
| 2d1cA | 3.44 | 3.82 | 3.96 | 3.96 | 0.35 | 0.37 | 0.02 | 0.79 |
| 2d7iA | 2.97 | 3.15 | 3.33 | 3.39 | 0.14 | 0.09 | 0.10 | 0.99 |
| 2dfyC | 2.40 | 2.60 | 3.12 | 4.20 | 0.84 | 0.83 | 0.80 | 0.08 |
| 2dlaA | 2.08 | 4.10 | 4.47 | 4.89 | 0.96 | 0.53 | 0.46 | 0.01 |
| 2e9hA | 1.56 | 1.35 | 1.44 | 4.12 | 0.99 | 1.00 | 1.00 | 0.12 |
| 2e9xB | 1.74 | 2.46 | 2.31 | 3.42 | 1.00 | 0.99 | 0.99 | 0.00 |
| 2evrA | 3.72 | 3.48 | 3.25 | 3.92 | 0.35 | 0.39 | 0.31 | 0.23 |

| PDB ID | MAE | | | | Top-*L* | | | |
|---|---|---|---|---|---|---|---|---|
| | DeepIDDP | DeepIDDP[1] | DeepIDDP[2] | DeepIDDP[3] | DeepIDDP | DeepIDDP[1] | DeepIDDP[2] | DeepIDDP[3] |
| 2ew9A | 3.19 | 3.91 | 3.47 | 4.14 | 0.58 | 0.27 | 0.56 | 0.17 |
| 2ewfA | 3.51 | 3.50 | 3.72 | 3.97 | 0.57 | 0.56 | 0.46 | 0.35 |
| 2fd5A | 1.28 | 3.43 | 3.35 | 3.66 | 1.00 | 0.96 | 0.95 | 0.38 |
| 2g3pA | 1.96 | 2.38 | 3.30 | 4.39 | 0.97 | 0.91 | 0.67 | 0.29 |
| 2gg6A | 2.50 | 2.00 | 2.57 | 3.61 | 0.89 | 0.96 | 0.86 | 0.63 |
| 2gh8A | 3.71 | 3.72 | 3.79 | 3.83 | 0.15 | 0.19 | 0.06 | 0.24 |
| 2gsyE | 3.54 | 4.08 | 4.24 | 4.25 | 0.56 | 0.28 | 0.21 | 0.12 |
| 2gt1A | 2.00 | 1.58 | 1.73 | 2.59 | 0.91 | 0.94 | 0.97 | 0.63 |
| 2gzaC | 1.98 | 1.99 | 4.01 | 4.30 | 0.95 | 0.99 | 0.41 | 0.83 |
| 2gzoA | 1.81 | 1.84 | 2.06 | 4.02 | 1.00 | 0.97 | 0.97 | 0.47 |
| 2hjqA | 1.74 | 2.50 | 3.03 | 3.73 | 0.24 | 0.17 | 0.17 | 0.47 |
| 2hwjA | 2.52 | 3.96 | 4.32 | 3.34 | 0.83 | 0.82 | 0.06 | 0.90 |
| 2ii2A | 2.88 | 3.34 | 3.51 | 3.61 | 0.78 | 0.71 | 0.66 | 0.58 |
| 2ijd1 | 2.74 | 2.87 | 3.19 | 3.22 | 0.11 | 0.07 | 0.06 | 0.40 |
| 2iu7A | 1.66 | 1.70 | 1.69 | 2.78 | 0.83 | 0.83 | 0.83 | 0.52 |
| 2iw2A | 3.19 | 3.40 | 3.51 | 4.11 | 0.56 | 0.48 | 0.44 | 0.85 |
| 2j2cA | 3.08 | 3.73 | 3.50 | 4.09 | 0.77 | 0.65 | 0.77 | 0.10 |
| 2jz4A | 2.68 | 2.85 | 3.04 | 3.11 | 0.14 | 0.12 | 0.03 | 0.88 |
| 2kdyA | 2.21 | 2.69 | 3.54 | 4.14 | 0.95 | 0.93 | 0.74 | 0.40 |
| 2kfwA | 2.18 | 2.87 | 3.07 | 4.23 | 0.56 | 0.48 | 0.45 | 0.46 |
| 2kn4A | 1.83 | 3.05 | 3.14 | 3.47 | 0.20 | 0.08 | 0.07 | 0.06 |
| 2l9yA | 2.17 | 2.54 | 2.98 | 4.29 | 0.74 | 0.60 | 0.56 | 0.23 |
| 2mbgA | 1.92 | 2.14 | 3.73 | 3.81 | 0.80 | 0.76 | 0.36 | 0.88 |
| 2nsfA | 1.59 | 1.27 | 1.79 | 2.88 | 0.99 | 1.00 | 0.98 | 0.91 |
| 2ntyB | 2.40 | 2.63 | 3.98 | 4.24 | 0.77 | 0.71 | 0.47 | 0.62 |
| 2nykA | 2.79 | 2.86 | 4.15 | 4.99 | 0.95 | 0.88 | 0.87 | 0.16 |
| 2o6yA | 2.22 | 1.86 | 2.56 | 4.08 | 0.84 | 0.70 | 0.64 | 0.02 |
| 2olsA | 3.59 | 3.39 | 3.52 | 4.12 | 0.61 | 0.65 | 0.68 | 0.42 |
| 2owbA | 1.24 | 4.24 | 4.34 | 1.90 | 1.00 | 0.19 | 0.17 | 0.31 |
| 2piaA | 2.44 | 2.01 | 2.50 | 3.01 | 0.98 | 1.00 | 0.99 | 0.46 |
| 2qfiA | 2.13 | 2.19 | 3.03 | 3.74 | 0.63 | 0.60 | 0.57 | 0.00 |
| 2qp2A | 3.87 | 3.94 | 3.94 | 3.94 | 0.04 | 0.04 | 0.00 | 0.64 |
| 2qygA | 1.83 | 2.10 | 2.58 | 4.20 | 0.86 | 0.74 | 0.67 | 0.08 |
| 2r3vA | 2.68 | 2.18 | 2.72 | 3.42 | 0.90 | 0.99 | 0.96 | 0.87 |
| 2r58A | 1.75 | 1.93 | 2.96 | 4.30 | 0.99 | 0.94 | 0.76 | 0.30 |
| 2r5wB | 2.45 | 2.07 | 2.39 | 3.72 | 0.85 | 0.85 | 0.93 | 0.81 |
| 2r7dA | 2.17 | 2.65 | 2.90 | 3.96 | 0.99 | 0.96 | 0.93 | 0.92 |
| 2ra1A | 3.94 | 4.42 | 4.92 | 5.01 | 0.70 | 0.64 | 0.24 | 0.04 |
| 2uu7A | 3.14 | 4.42 | 4.92 | 4.24 | 0.88 | 0.90 | 0.89 | 0.85 |
| 2uwnA | 2.27 | 2.24 | 2.72 | 3.47 | 0.99 | 0.96 | 0.95 | 0.38 |
| 2v0nA | 1.62 | 1.92 | 2.47 | 3.82 | 0.64 | 0.64 | 0.63 | 0.04 |
| 2v5dA | 3.51 | 3.30 | 3.98 | 4.19 | 0.97 | 0.74 | 0.72 | 0.34 |
| 2vgmA | 2.62 | 2.72 | 3.26 | 4.13 | 0.97 | 0.68 | 0.57 | 0.36 |
| 2w4bA | 2.48 | 3.20 | 3.94 | 4.13 | 0.95 | 0.45 | 0.13 | 0.12 |

| PDB ID | MAE | | | | Top-*L* | | | |
|---|---|---|---|---|---|---|---|---|
| | DeepIDDP | DeepIDDP[1] | DeepIDDP[2] | DeepIDDP[3] | DeepIDDP | DeepIDDP[1] | DeepIDDP[2] | DeepIDDP[3] |
| 2w4mA | 1.64 | 1.52 | 1.64 | 2.60 | 0.98 | 0.98 | 0.96 | 0.78 |
| 2wqrB | 2.67 | 2.84 | 3.41 | 3.81 | 0.54 | 0.54 | 0.45 | 0.00 |
| 2x0cA | 2.21 | 2.82 | 3.47 | 4.29 | 0.31 | 0.24 | 0.21 | 0.02 |
| 2x7iA | 2.32 | 2.02 | 2.13 | 2.75 | 0.94 | 0.96 | 0.95 | 0.06 |
| 2x8kC | 2.61 | 3.34 | 3.81 | 4.15 | 0.72 | 0.58 | 0.48 | 0.63 |
| 2xt6A | 3.94 | 3.62 | 3.72 | 4.47 | 0.21 | 0.58 | 0.60 | 0.15 |
| 2y25B | 2.19 | 4.55 | 5.17 | 5.47 | 0.98 | 0.59 | 0.46 | 0.44 |
| 2y51A | 2.27 | 2.06 | 3.01 | 3.95 | 0.81 | 0.61 | 0.82 | 0.52 |
| 2yb0E | 1.37 | 1.49 | 2.77 | 3.61 | 1.00 | 1.00 | 0.85 | 0.68 |
| 2yilA | 2.28 | 4.14 | 4.32 | 4.50 | 0.69 | 0.16 | 0.14 | 0.90 |
| 2yk0A | 3.82 | 3.57 | 3.86 | 4.11 | 0.60 | 0.69 | 0.63 | 0.45 |
| 2yrqA | 2.72 | 3.20 | 3.59 | 3.88 | 0.17 | 0.12 | 0.12 | 0.95 |
| 2z86C | 4.05 | 4.13 | 4.14 | 4.17 | 0.40 | 0.30 | 0.32 | 0.00 |
| 2zpaB | 3.86 | 3.28 | 3.68 | 4.04 | 0.76 | 0.81 | 0.78 | 0.10 |
| 2zxcA | 3.49 | 3.23 | 3.83 | 3.97 | 0.59 | 0.61 | 0.58 | 0.17 |
| 2zzqA | 3.96 | 4.05 | 4.19 | 4.23 | 0.60 | 0.42 | 0.28 | 0.20 |
| 3a1iA | 3.13 | 3.13 | 3.15 | 3.25 | 0.06 | 0.05 | 0.09 | 0.03 |
| 3a45A | 2.40 | 2.07 | 2.32 | 2.98 | 0.92 | 1.00 | 0.98 | 0.03 |
| 3a56A | 3.76 | 3.71 | 3.87 | 3.87 | 0.24 | 0.23 | 0.15 | 0.06 |
| 3afoA | 4.55 | 4.51 | 4.58 | 4.64 | 0.11 | 0.14 | 0.18 | 0.45 |
| 3ajvA | 2.73 | 3.84 | 3.96 | 3.80 | 0.74 | 0.20 | 0.13 | 0.19 |
| 3apoA | 4.54 | 3.90 | 3.95 | 4.53 | 0.79 | 0.79 | 0.76 | 0.71 |
| 3aqkA | 1.64 | 3.92 | 3.99 | 3.73 | 0.87 | 0.05 | 0.03 | 0.10 |
| 3arbA | 2.03 | 2.17 | 2.89 | 3.52 | 0.62 | 0.54 | 0.51 | 0.80 |
| 3aujG | 1.60 | 4.66 | 4.63 | 5.03 | 1.00 | 0.40 | 0.40 | 0.37 |
| 3b2zF | 3.10 | 2.09 | 2.58 | 4.10 | 0.82 | 0.93 | 0.79 | 0.72 |
| 3b43A | 3.07 | 3.70 | 3.80 | 4.15 | 0.21 | 0.15 | 0.10 | 0.12 |
| 3b7wA | 4.31 | 4.92 | 4.52 | 4.43 | 0.28 | 0.95 | 0.05 | 0.36 |
| 3bt1U | 2.76 | 3.59 | 4.05 | 4.20 | 0.92 | 0.80 | 0.60 | 0.78 |
| 3bt3A | 1.25 | 0.97 | 1.05 | 2.21 | 1.00 | 1.00 | 1.00 | 0.59 |
| 3bu2A | 2.63 | 2.65 | 3.23 | 4.07 | 0.54 | 0.56 | 0.32 | 0.29 |
| 3c1yA | 1.85 | 1.79 | 3.05 | 4.38 | 0.97 | 0.99 | 0.87 | 0.44 |
| 3c4tA | 2.93 | 2.61 | 2.20 | 4.31 | 0.53 | 0.55 | 0.59 | 0.24 |
| 3craA | 1.90 | 2.08 | 2.40 | 3.63 | 0.54 | 0.59 | 0.57 | 0.32 |
| 3cvzA | 2.38 | 3.13 | 4.19 | 4.65 | 0.93 | 0.81 | 0.60 | 0.42 |
| 3cw2C | 1.73 | 1.99 | 2.15 | 3.70 | 0.95 | 0.92 | 0.94 | 0.63 |
| 3d30A | 3.23 | 3.02 | 3.42 | 3.95 | 0.81 | 0.82 | 0.73 | 0.55 |
| 3dupA | 3.74 | 3.68 | 4.33 | 4.22 | 0.71 | 0.59 | 0.32 | 0.35 |
| 3eo5A | 2.90 | 3.03 | 3.70 | 4.43 | 0.77 | 0.75 | 0.62 | 0.07 |
| 3errA | 3.20 | 3.49 | 4.19 | 4.26 | 0.56 | 0.48 | 0.36 | 0.19 |
| 3eswA | 2.08 | 2.40 | 2.67 | 4.30 | 0.85 | 0.77 | 0.78 | 0.17 |
| 3eukH | 4.07 | 4.07 | 4.53 | 4.64 | 0.57 | 0.56 | 0.48 | 0.30 |
| 3f83A | 3.50 | 3.42 | 4.25 | 4.40 | 0.58 | 0.70 | 0.50 | 0.87 |
| 3fc3A | 2.26 | 3.23 | 3.31 | 4.42 | 0.89 | 0.74 | 0.81 | 0.12 |

| PDB ID | MAE | | | | Top-*L* | | | |
|---|---|---|---|---|---|---|---|---|
| | DeepIDDP | DeepIDDP[1] | DeepIDDP[2] | DeepIDDP[3] | DeepIDDP | DeepIDDP[1] | DeepIDDP[2] | DeepIDDP[3] |
| 3fi7A | 1.29 | 1.41 | 1.43 | 4.70 | 1.00 | 1.00 | 1.00 | 0.98 |
| 3fvvA | 1.14 | 1.27 | 1.55 | 4.15 | 1.00 | 1.00 | 1.00 | 0.76 |
| 3g79A | 2.29 | 3.07 | 3.22 | 3.06 | 0.75 | 0.56 | 0.51 | 0.99 |
| 3gbgA | 2.40 | 2.31 | 2.59 | 4.31 | 1.00 | 1.00 | 0.98 | 0.68 |
| 3gf5B | 3.32 | 3.84 | 3.73 | 4.20 | 0.66 | 0.50 | 0.42 | 0.27 |
| 3gmsA | 2.96 | 2.93 | 3.20 | 4.03 | 0.60 | 0.62 | 0.67 | 0.32 |
| 3h2tA | 3.39 | 3.92 | 4.02 | 4.02 | 0.42 | 0.21 | 0.01 | 0.76 |
| 3h5cB | 2.59 | 3.48 | 4.42 | 3.92 | 1.00 | 1.00 | 0.16 | 0.42 |
| 3hcsA | 1.82 | 2.18 | 2.02 | 5.19 | 0.79 | 0.73 | 0.73 | 0.42 |
| 3hjlA | 2.18 | 2.60 | 3.64 | 4.05 | 0.36 | 0.33 | 0.18 | 0.00 |
| 3hyiA | 3.40 | 4.54 | 4.79 | 4.82 | 0.06 | 0.02 | 0.02 | 0.79 |
| 3hzzB | 1.54 | 2.88 | 3.22 | 3.99 | 1.00 | 0.90 | 0.83 | 0.56 |
| 3i2dA | 2.80 | 1.81 | 2.99 | 4.09 | 0.67 | 0.79 | 0.66 | 0.22 |
| 3iam2 | 2.51 | 2.07 | 2.74 | 2.44 | 0.85 | 0.92 | 0.80 | 0.02 |
| 3ibjA | 3.03 | 2.75 | 3.28 | 4.53 | 0.61 | 0.70 | 0.73 | 0.00 |
| 3ifrA | 2.44 | 1.68 | 2.01 | 3.38 | 0.95 | 1.00 | 0.96 | 0.00 |
| 3ippB | 3.27 | 3.26 | 3.54 | 3.79 | 0.72 | 0.72 | 0.65 | 0.17 |
| 3isqA | 3.57 | 3.30 | 3.42 | 4.09 | 0.61 | 0.64 | 0.65 | 0.01 |
| 3j7aK | 1.51 | 0.90 | 0.99 | 1.63 | 0.98 | 1.00 | 1.00 | 0.93 |
| 3jymB | 2.71 | 2.72 | 3.38 | 3.99 | 0.64 | 0.67 | 0.46 | 0.91 |
| 3k1rA | 3.51 | 2.22 | 3.18 | 4.10 | 0.45 | 0.86 | 0.77 | 0.07 |
| 3k2iA | 3.74 | 3.84 | 4.12 | 4.21 | 0.59 | 0.47 | 0.41 | 0.22 |
| 3kbgA | 2.72 | 2.39 | 3.13 | 3.84 | 0.94 | 0.91 | 0.79 | 0.40 |
| 3kh5A | 2.36 | 2.70 | 3.18 | 3.45 | 0.96 | 0.96 | 0.91 | 0.00 |
| 3kjpA | 2.32 | 2.76 | 3.40 | 3.96 | 0.75 | 0.67 | 0.63 | 0.01 |
| 3kq4B | 3.48 | 3.51 | 3.81 | 4.22 | 0.60 | 0.57 | 0.55 | 0.38 |
| 3kt1A | 3.89 | 3.93 | 4.16 | 4.17 | 0.48 | 0.46 | 0.31 | 0.27 |
| 3ktmE | 1.59 | 2.15 | 2.69 | 3.11 | 0.94 | 0.94 | 0.81 | 0.77 |
| 3kwlA | 3.43 | 3.65 | 3.86 | 4.21 | 0.57 | 0.44 | 0.25 | 0.20 |
| 3kzwA | 1.27 | 1.19 | 1.72 | 3.69 | 0.99 | 0.98 | 0.94 | 0.17 |
| 3l76A | 3.84 | 3.89 | 3.93 | 3.96 | 0.12 | 0.08 | 0.07 | 0.06 |
| 3ld1A | 3.33 | 4.60 | 4.61 | 4.61 | 0.63 | 0.06 | 0.05 | 0.93 |
| 3lsgA | 1.61 | 1.08 | 1.04 | 1.75 | 0.98 | 1.00 | 1.00 | 0.44 |
| 3m1uA | 4.25 | 4.16 | 4.48 | 4.49 | 0.20 | 0.22 | 0.22 | 0.03 |
| 3mc8A | 1.92 | 1.84 | 2.17 | 3.99 | 0.98 | 0.98 | 0.97 | 0.84 |
| 3me4A | 1.55 | 2.90 | 3.13 | 3.44 | 0.99 | 0.84 | 0.81 | 0.03 |
| 3ml4C | 2.32 | 1.59 | 2.15 | 4.34 | 0.90 | 1.00 | 0.97 | 0.76 |
| 3mw8A | 3.15 | 2.93 | 3.91 | 5.28 | 0.74 | 0.81 | 0.75 | 0.72 |
| 3mwcA | 4.14 | 4.17 | 4.32 | 4.55 | 0.27 | 0.25 | 0.37 | 0.04 |
| 3mx2B | 2.64 | 4.26 | 4.24 | 4.24 | 0.85 | 0.00 | 0.06 | 0.33 |
| 3mzfA | 4.23 | 4.58 | 4.61 | 4.60 | 0.13 | 0.01 | 0.06 | 0.17 |
| 3njaB | 2.26 | 3.12 | 3.34 | 4.32 | 0.88 | 0.72 | 0.73 | 0.05 |
| 3npfA | 3.79 | 3.87 | 4.09 | 4.28 | 0.54 | 0.23 | 0.32 | 0.16 |
| 3nqiA | 3.09 | 2.75 | 3.39 | 3.86 | 0.53 | 0.64 | 0.62 | 0.35 |

| PDB ID | MAE | | | | Top-*L* | | | |
|---|---|---|---|---|---|---|---|---|
| | DeepIDDP | DeepIDDP[1] | DeepIDDP[2] | DeepIDDP[3] | DeepIDDP | DeepIDDP[1] | DeepIDDP[2] | DeepIDDP[3] |
| 3nsjA | 4.06 | 4.07 | 4.56 | 4.66 | 0.38 | 0.41 | 0.33 | 0.02 |
| 3nt8A | 3.29 | 3.44 | 3.50 | 3.51 | 0.16 | 0.07 | 0.03 | 0.03 |
| 3ntkA | 1.64 | 2.17 | 2.42 | 2.67 | 0.99 | 0.98 | 0.93 | 0.90 |
| 3oaaG | 2.58 | 4.61 | 4.73 | 4.71 | 1.00 | 0.39 | 0.31 | 0.71 |
| 3ob8A | 3.32 | 2.93 | 3.17 | 4.07 | 0.92 | 0.92 | 0.93 | 0.41 |
| 3og5A | 2.07 | 1.93 | 2.26 | 2.67 | 0.99 | 0.98 | 0.95 | 0.54 |
| 3oh0A | 4.16 | 4.41 | 4.44 | 4.44 | 0.45 | 0.23 | 0.17 | 0.00 |
| 3opfB | 3.04 | 2.81 | 3.47 | 4.19 | 0.92 | 0.95 | 0.88 | 0.38 |
| 3orjA | 3.06 | 3.02 | 3.66 | 4.11 | 0.76 | 0.77 | 0.64 | 0.72 |
| 3p53A | 3.98 | 4.02 | 4.17 | 4.26 | 0.47 | 0.46 | 0.14 | 0.04 |
| 3pcsB | 3.63 | 3.93 | 4.15 | 4.16 | 0.59 | 0.50 | 0.28 | 0.11 |
| 3plaA | 1.63 | 3.23 | 3.50 | 4.79 | 0.98 | 0.69 | 0.71 | 0.63 |
| 3po3S | 2.86 | 5.29 | 5.29 | 5.29 | 0.74 | 0.96 | 0.00 | 0.03 |
| 3ptyA | 3.00 | 2.85 | 3.49 | 4.30 | 0.83 | 0.81 | 0.64 | 0.96 |
| 3pvlA | 4.15 | 3.32 | 3.43 | 4.24 | 0.77 | 0.61 | 0.47 | 0.47 |
| 3pxpA | 1.78 | 1.30 | 1.36 | 1.47 | 0.14 | 0.46 | 0.66 | 0.05 |
| 3qavA | 2.20 | 2.59 | 4.66 | 4.59 | 0.97 | 1.00 | 0.98 | 0.39 |
| 3qe9Y | 2.37 | 2.72 | 2.84 | 3.80 | 1.00 | 1.00 | 0.27 | 0.17 |
| 3qf4B | 1.84 | 1.39 | 1.55 | 2.41 | 0.31 | 0.19 | 0.19 | 0.53 |
| 3qjjA | 3.28 | 3.45 | 3.78 | 4.18 | 1.00 | 1.00 | 1.00 | 0.95 |
| 3qjoA | 2.57 | 2.78 | 3.38 | 3.96 | 0.92 | 0.84 | 0.83 | 0.62 |
| 3qphA | 3.05 | 2.21 | 3.25 | 3.95 | 0.81 | 0.75 | 0.63 | 0.10 |
| 3qtdA | 2.59 | 2.42 | 2.70 | 3.81 | 0.86 | 0.94 | 0.86 | 0.72 |
| 3qyeA | 4.96 | 4.95 | 4.01 | 5.07 | 0.99 | 1.00 | 1.00 | 0.93 |
| 3r05A | 1.66 | 3.51 | 3.86 | 5.53 | 0.95 | 0.96 | 0.91 | 0.89 |
| 3r6bA | 1.99 | 2.09 | 3.07 | 3.91 | 0.57 | 0.30 | 0.30 | 0.01 |
| 3rfyA | 3.07 | 2.96 | 3.91 | 4.32 | 0.85 | 0.86 | 0.67 | 0.26 |
| 3rh7A | 3.49 | 3.45 | 3.58 | 4.36 | 0.74 | 0.76 | 0.64 | 0.97 |
| 3rimA | 2.10 | 2.00 | 2.60 | 4.28 | 0.67 | 0.71 | 0.65 | 0.02 |
| 3rrpA | 2.53 | 3.86 | 3.91 | 3.91 | 0.94 | 0.93 | 0.90 | 0.41 |
| 3rwxA | 3.02 | 2.76 | 3.76 | 3.86 | 0.43 | 0.08 | 0.00 | 0.00 |
| 3sb4A | 2.80 | 2.69 | 2.73 | 4.47 | 0.56 | 0.64 | 0.51 | 0.18 |
| 3seoB | 2.33 | 2.27 | 2.65 | 2.75 | 0.65 | 0.72 | 0.68 | 0.09 |
| 3soaA | 2.44 | 2.24 | 3.58 | 4.06 | 0.98 | 0.99 | 0.96 | 0.72 |
| 3spgA | 2.82 | 2.72 | 3.76 | 4.27 | 0.31 | 0.30 | 0.29 | 0.19 |
| 3swjA | 4.28 | 4.33 | 4.34 | 4.34 | 0.72 | 0.72 | 0.59 | 0.37 |
| 3t58B | 1.21 | 1.26 | 1.68 | 3.09 | 0.32 | 0.22 | 0.03 | 0.03 |
| 3t7jA | 3.12 | 4.72 | 4.75 | 4.77 | 1.00 | 1.00 | 1.00 | 0.01 |
| 3tixD | 2.52 | 1.98 | 2.69 | 3.76 | 0.57 | 0.07 | 0.02 | 0.00 |
| 3tp9A | 2.73 | 2.63 | 3.37 | 3.67 | 0.89 | 0.97 | 0.87 | 0.20 |
| 3u07C | 3.93 | 4.01 | 4.26 | 4.32 | 0.79 | 0.81 | 0.75 | 0.80 |
| 3u0kA | 2.08 | 1.84 | 2.18 | 2.68 | 0.30 | 0.26 | 0.05 | 0.71 |
| 3u0oB | 3.06 | 4.63 | 4.80 | 4.79 | 0.97 | 0.96 | 0.94 | 0.89 |
| 3u9gA | 4.06 | 4.07 | 4.56 | 4.66 | 0.98 | 0.38 | 0.28 | 0.86 |

| PDB ID | MAE | | | | Top-$L$ | | | |
|---|---|---|---|---|---|---|---|---|
| | DeepIDDP | DeepIDDP[1] | DeepIDDP[2] | DeepIDDP[3] | DeepIDDP | DeepIDDP[1] | DeepIDDP[2] | DeepIDDP[3] |
| 3ua3A | 3.46 | 3.33 | 3.81 | 4.39 | 0.81 | 0.84 | 0.67 | 0.44 |
| 3ub1D | 3.00 | 5.31 | 5.32 | 5.32 | 0.52 | 0.01 | 0.01 | 0.18 |
| 3ubhA | 2.35 | 3.89 | 3.66 | 4.30 | 0.70 | 0.48 | 0.42 | 0.40 |
| 3uitD | 1.95 | 1.98 | 2.84 | 3.11 | 0.23 | 0.24 | 0.17 | 0.06 |
| 3uj0A | 3.06 | 3.17 | 3.54 | 3.72 | 0.63 | 0.55 | 0.45 | 0.25 |
| 3uo3A | 2.84 | 3.70 | 3.80 | 4.33 | 0.66 | 0.55 | 0.62 | 0.24 |
| 3v7oB | 1.56 | 3.24 | 3.74 | 4.14 | 0.23 | 0.10 | 0.10 | 0.50 |
| 3vlaA | 4.48 | 4.81 | 4.80 | 4.79 | 0.41 | 0.01 | 0.00 | 0.39 |
| 3vn4A | 3.26 | 3.90 | 4.56 | 4.62 | 0.85 | 0.70 | 0.56 | 0.78 |
| 3vr8B | 2.20 | 1.68 | 1.90 | 2.38 | 0.96 | 1.00 | 0.98 | 0.62 |
| 3vsmA | 3.92 | 4.49 | 4.51 | 4.50 | 0.53 | 0.17 | 0.11 | 0.07 |
| 3vstA | 3.60 | 4.01 | 4.19 | 4.20 | 0.62 | 0.70 | 0.15 | 0.02 |
| 3w1bA | 3.55 | 3.61 | 3.97 | 4.17 | 0.47 | 0.39 | 0.31 | 0.23 |
| 3w2wA | 3.32 | 2.79 | 3.21 | 4.16 | 0.51 | 0.86 | 0.80 | 0.45 |
| 3wkuA | 2.00 | 2.02 | 2.83 | 3.86 | 0.62 | 0.63 | 0.55 | 0.44 |
| 3zh9B | 2.54 | 2.57 | 3.36 | 3.83 | 0.82 | 0.79 | 0.54 | 0.37 |
| 3zniA | 2.95 | 3.20 | 3.97 | 4.18 | 0.85 | 0.79 | 0.58 | 0.24 |
| 3zvmA | 3.25 | 3.07 | 3.50 | 3.61 | 0.24 | 0.34 | 0.32 | 0.64 |
| 4acoA | 4.10 | 4.15 | 4.20 | 4.20 | 0.15 | 0.16 | 0.07 | 0.72 |
| 4aimA | 3.26 | 3.77 | 3.77 | 4.04 | 0.65 | 0.35 | 0.36 | 0.75 |
| 4ak1A | 4.08 | 4.41 | 4.41 | 4.42 | 0.24 | 0.14 | 0.09 | 0.09 |
| 4alzA | 3.25 | 4.09 | 4.60 | 4.54 | 0.57 | 0.26 | 0.07 | 0.06 |
| 4ap5A | 2.80 | 2.55 | 2.88 | 4.11 | 0.70 | 0.78 | 0.75 | 0.14 |
| 4aq1A | 3.97 | 4.08 | 4.12 | 4.14 | 0.16 | 0.09 | 0.05 | 0.03 |
| 4aqfB | 3.52 | 3.96 | 4.34 | 4.46 | 0.65 | 0.41 | 0.27 | 0.09 |
| 4ax8A | 3.85 | 3.96 | 3.96 | 4.03 | 0.15 | 0.10 | 0.16 | 0.26 |
| 4axdA | 3.71 | 3.59 | 3.96 | 4.16 | 0.58 | 0.57 | 0.42 | 0.08 |
| 4b21A | 2.70 | 2.64 | 2.93 | 3.97 | 0.71 | 0.77 | 0.75 | 0.38 |
| 4b3iA | 3.62 | 3.64 | 3.74 | 4.07 | 0.46 | 0.41 | 0.44 | 0.25 |
| 4bd9B | 1.90 | 2.56 | 3.01 | 4.56 | 0.99 | 0.98 | 0.75 | 0.94 |
| 4bfiB | 1.64 | 2.55 | 3.05 | 3.58 | 0.88 | 0.62 | 0.59 | 0.48 |
| 4bt9B | 2.09 | 3.08 | 3.26 | 3.45 | 0.17 | 0.08 | 0.08 | 0.12 |
| 4c0aB | 2.17 | 2.03 | 3.12 | 4.42 | 0.99 | 0.99 | 0.97 | 0.46 |
| 4c0sA | 3.59 | 4.16 | 4.33 | 4.30 | 0.66 | 0.62 | 0.24 | 0.46 |
| 4cczA | 3.15 | 3.63 | 4.04 | 3.84 | 0.64 | 0.88 | 0.21 | 0.99 |
| 4d0nB | 1.59 | 2.09 | 2.87 | 4.92 | 0.86 | 0.80 | 0.78 | 0.18 |
| 4d1iG | 3.11 | 3.69 | 3.81 | 4.26 | 0.58 | 0.32 | 0.33 | 0.26 |
| 4dimA | 1.61 | 2.29 | 4.23 | 3.38 | 1.00 | 0.99 | 0.06 | 0.62 |
| 4dj3A | 2.78 | 2.19 | 2.99 | 4.04 | 0.63 | 0.82 | 0.68 | 0.01 |
| 4dqaA | 3.16 | 3.70 | 3.70 | 3.70 | 0.16 | 0.08 | 0.00 | 0.75 |
| 4dt4A | 3.08 | 2.98 | 2.73 | 5.68 | 0.89 | 0.86 | 0.98 | 0.33 |
| 4dtfA | 4.16 | 5.12 | 5.26 | 5.26 | 0.84 | 0.77 | 0.06 | 0.24 |
| 4eo3A | 2.71 | 4.00 | 3.93 | 4.08 | 0.15 | 0.02 | 0.02 | 0.99 |
| 4eogA | 2.88 | 4.32 | 4.98 | 5.27 | 0.98 | 0.84 | 0.75 | 0.29 |

| PDB ID | MAE | | | | Top-*L* | | | |
|---|---|---|---|---|---|---|---|---|
| | DeepIDDP | DeepIDDP[1] | DeepIDDP[2] | DeepIDDP[3] | DeepIDDP | DeepIDDP[1] | DeepIDDP[2] | DeepIDDP[3] |
| 4etxA | 2.13 | 3.43 | 3.97 | 4.11 | 0.89 | 0.55 | 0.46 | 0.35 |
| 4ewtA | 2.61 | 2.67 | 3.35 | 3.74 | 0.66 | 0.65 | 0.68 | 0.54 |
| 4f23A | 3.64 | 3.77 | 4.35 | 4.47 | 0.50 | 0.49 | 0.22 | 0.13 |
| 4fe9A | 3.99 | 4.04 | 4.20 | 4.23 | 0.28 | 0.23 | 0.23 | 0.08 |
| 4fguA | 2.81 | 2.86 | 3.60 | 4.15 | 0.84 | 0.85 | 0.77 | 0.04 |
| 4fkcA | 1.36 | 1.42 | 1.79 | 3.36 | 0.91 | 0.91 | 0.85 | 0.09 |
| 4fxkC | 3.35 | 3.07 | 4.05 | 4.18 | 0.42 | 0.56 | 0.38 | 0.90 |
| 4fzbC | 2.25 | 2.05 | 2.23 | 3.52 | 0.88 | 0.90 | 0.90 | 0.19 |
| 4g1pA | 2.47 | 2.37 | 3.04 | 3.72 | 0.81 | 0.79 | 0.67 | 0.07 |
| 4gbyA | 1.82 | 1.96 | 2.16 | 3.28 | 1.00 | 1.00 | 1.00 | 0.00 |
| 4gfqA | 2.59 | 3.05 | 3.20 | 3.96 | 0.75 | 0.57 | 0.52 | 0.00 |
| 4ggmX | 3.24 | 3.44 | 3.34 | 4.20 | 0.51 | 0.38 | 0.52 | 0.25 |
| 4gslA | 2.77 | 4.83 | 4.89 | 4.89 | 0.17 | 0.01 | 0.01 | 0.45 |
| 4gyjA | 3.14 | 3.18 | 4.32 | 4.23 | 0.76 | 0.79 | 0.10 | 0.77 |
| 4h2aA | 3.93 | 4.02 | 4.21 | 4.29 | 0.63 | 0.49 | 0.42 | 0.25 |
| 4h3tA | 3.18 | 3.18 | 3.60 | 4.22 | 0.46 | 0.50 | 0.48 | 0.38 |
| 4hmoA | 3.01 | 3.36 | 3.41 | 3.06 | 0.84 | 0.76 | 0.78 | 0.05 |
| 4hvzA | 2.07 | 2.44 | 2.71 | 4.33 | 0.76 | 0.77 | 0.65 | 0.17 |
| 4i5sB | 1.87 | 2.24 | 2.56 | 4.36 | 0.70 | 0.52 | 0.56 | 0.00 |
| 4ie6A | 4.07 | 4.32 | 4.51 | 4.51 | 0.49 | 0.33 | 0.10 | 0.00 |
| 4iggB | 3.60 | 3.95 | 4.11 | 4.20 | 0.71 | 0.04 | 0.11 | 0.00 |
| 4il6B | 4.37 | 4.40 | 4.67 | 4.77 | 0.39 | 0.33 | 0.39 | 0.29 |
| 4indA | 3.87 | 3.89 | 4.18 | 4.22 | 0.45 | 0.44 | 0.24 | 0.45 |
| 4j9vA | 2.90 | 2.58 | 3.41 | 4.31 | 0.97 | 1.00 | 0.96 | 0.86 |
| 4jdzB | 3.67 | 3.55 | 3.83 | 3.87 | 0.38 | 0.36 | 0.35 | 0.16 |
| 4jxkA | 2.25 | 2.28 | 2.52 | 4.02 | 0.92 | 0.92 | 0.93 | 0.02 |
| 4k3bA | 2.84 | 4.06 | 4.12 | 4.08 | 0.52 | 0.23 | 0.22 | 0.00 |
| 4kc3B | 3.15 | 3.46 | 3.72 | 4.35 | 0.72 | 0.64 | 0.60 | 0.22 |
| 4kikB | 3.41 | 3.41 | 3.48 | 4.19 | 0.66 | 0.72 | 0.75 | 0.23 |
| 4kwuA | 4.16 | 4.17 | 4.19 | 4.20 | 0.11 | 0.18 | 0.08 | 0.01 |
| 4l5gA | 2.67 | 1.90 | 1.89 | 3.46 | 0.89 | 0.94 | 0.94 | 0.13 |
| 4lmfA | 2.46 | 2.69 | 3.84 | 4.86 | 0.85 | 0.84 | 0.75 | 0.34 |
| 4lpqA | 3.03 | 3.18 | 3.23 | 4.41 | 0.78 | 0.68 | 0.86 | 0.26 |
| 4lziA | 3.15 | 3.49 | 3.74 | 4.21 | 0.80 | 0.61 | 0.60 | 0.13 |
| 4m00A | 3.15 | 3.24 | 4.50 | 4.74 | 0.71 | 0.76 | 0.49 | 0.29 |
| 4m8m | 3.96 | 3.59 | 4.03 | 4.14 | 0.39 | 0.38 | 0.30 | 0.00 |
| 4m8rA | 3.53 | 3.41 | 3.83 | 3.86 | 0.42 | 0.34 | 0.20 | 0.49 |
| 4m9pA | 3.43 | 3.96 | 3.96 | 4.14 | 0.70 | 0.37 | 0.59 | 0.21 |
| 4mzyA | 3.80 | 4.59 | 4.65 | 4.68 | 0.43 | 0.20 | 0.23 | 0.78 |
| 4n06B | 2.62 | 2.08 | 2.72 | 2.80 | 0.77 | 0.85 | 0.76 | 0.00 |
| 4nj5A | 3.53 | 3.86 | 4.03 | 4.22 | 0.76 | 0.55 | 0.50 | 0.98 |
| 4onyA | 3.84 | 3.85 | 3.99 | 4.21 | 0.29 | 0.23 | 0.26 | 0.17 |
| 4opaB | 2.00 | 3.49 | 4.66 | 4.64 | 0.74 | 0.45 | 0.08 | 0.72 |
| 4pt5A | 2.09 | 1.95 | 3.02 | 4.41 | 0.98 | 1.00 | 0.96 | 0.81 |

| PDB ID | MAE | | | | Top-*L* | | | |
|---|---|---|---|---|---|---|---|---|
| | DeepIDDP | DeepIDDP[1] | DeepIDDP[2] | DeepIDDP[3] | DeepIDDP | DeepIDDP[1] | DeepIDDP[2] | DeepIDDP[3] |
| 4pyhA | 4.28 | 4.12 | 4.62 | 4.89 | 4pyhA | 0.51 | 0.55 | 0.44 |
| 4qkuB | 2.07 | 2.03 | 2.57 | 2.75 | 4qkuB | 0.99 | 0.99 | 0.96 |
| 4rg1A | 3.86 | 3.81 | 4.22 | 5.02 | 4rg1A | 0.78 | 0.80 | 0.65 |
| 4up9A | 3.29 | 4.21 | 4.57 | 4.43 | 4up9A | 0.32 | 0.26 | 0.03 |
| 4uwhA | 3.23 | 2.87 | 2.91 | 4.04 | 4uwhA | 0.79 | 0.81 | 0.83 |
| 4w7sA | 3.11 | 3.43 | 3.68 | 3.77 | 4w7sA | 0.15 | 0.08 | 0.05 |

### References

[1] W. Kabsch. A solution for the best rotation to relate two sets of vectors. Acta Crystallographica Section A: Crystal Physics, Diffraction, Theoretical and General Crystallography, 32(5):922–923,1976.
